# NEK9 ablation rewires docetaxel resistance through induction of ERK-mediated cancer cell pyroptosis

**DOI:** 10.1101/2024.09.10.612209

**Authors:** Shamima Azma Ansari, Sibasish Mohanty, Pallavi Mohapatra, Rachna Rath, Dillip Muduli, Saroj Kumar Das Majumdar, Rajeeb K. Swain, Rupesh Dash

## Abstract

Docetaxel alone or in combination with other drugs is the most common chemotherapy regimen for several neoplasms including advanced OSCC. Unfortunately, chemoresistance leads to relapse and continued tumor growth. It is therefore important to explore the causative factors for docetaxel resistance. In this study, we performed a CRISPR-based kinome screening that identified Never In Mitosis Gene-A Related Kinase-9 (NEK9) as a major player of docetaxel resistance in OSCC, prostate, and pancreatic cancer lines. NEK9 expression was upregulated in tumor samples of chemotherapy non-responders compared to responder OSCC patients. Our validation data suggests selectively knocking out NEK9 sensitizes cancer cells to docetaxel. Mechanistically, we found that ablation of NEK9 induces DNA damage, activating ERK(p-T202/Y204) that leads to Gasdermin-E mediated Cancer Cell pyroptosis. The in-vitro kinase activity assay identified fostamatinib as a potent inhibitor of NEK9. The xenograft data suggest that fostamatinib restores docetaxel sensitivity and facilitates a significant reduction of tumor burden. Overall, our data suggests a novel combination of fostamatinib and docetaxel needs further clinical investigation in advanced OSCC.

## 1. Introduction

Taxanes are microtubule-stabilizing diterpenes isolated from trees belonging to *Taxus brevifolia* species. The clinically approved antineoplastic includes naturally occurring Paclitaxel and semi-synthetic derivatives like Docetaxel and Cabazitaxel (Dumontet & Jordan, 2010; Wani *et al*, 1971). Docetaxel, either as a standalone treatment or in conjunction with other chemotherapy medications, is the most often employed chemotherapy regimen for individuals diagnosed with breast, lung, and prostate cancer (Tan *et al*, 2012). Cisplatin, 5 Fluorouracil (5FU), and Docetaxel together termed as TPF regimen is commonly prescribed for locally advanced Head and neck squamous cell carcinomas (HNSCC) (Gau *et al*, 2019). Docetaxel binds to β-tubulin with high affinity hence resulting in polymerization and stabilization of microtubules in the cells. This leads to blockage in the G1/M phase of the cell cycle followed by induction of apoptosis (Schiff *et al*, 1979). Additionally, Docetaxel is known to modulate the expression of cell cycle protein p27 and anti-apoptotic protein Bcl-2 (Mhaidat *et al*, 2007). However, chemoresistance leads to relapse, continued tumor growth, and metastatic spread. Drug resistance to Docetaxel could be attributed to impaired apoptosis, elevated expression of efflux transporters, and altered DNA damage responses (Hsieh *et al*, 2023). Drug resistance is one of the major factors for treatment failure in (Oral squamous cell carcinoma) OSCC and prostate carcinomas. Hence, it is important to identify the causative factor (s) that contribute significantly to drug resistance.

Kinases have a crucial role in cell signaling and are involved in nearly every cancer trait including growth, proliferation, angiogenesis, metastasis, evasion of antitumor immune responses, and drug resistance. More than 800 kinases exist in the human cells out of which 538 kinases are known to regulate different cell signalling. In HNSCC, kinases are also reported to regulate acquired drug resistance. Jin *et al*., in 2018 performed an RNAi-based kinome screening in HNSCC to find microtubule-associated serine/threonine kinase 1 (MAST1) as an important regulator of Cisplatin resistance. They found that MAST1 activates MAPK signaling in a cRaf-independent manner to drive Cisplatin resistance (Jin *et al*, 2018). We have performed a CRISPR-based kinome screening in 5FU resistant OSCC lines to identify Misshapen Like Kinase 1 (MINK1) as a key player in driving 5FU resistance. Mechanistically, we found that MINK1 modulates AKT phosphorylation at Ser473, which enables p-MDM2 (Ser166) mediated degradation of p53 (Mohanty *et al*, 2022). However, the potential role of kinases in Docetaxel-resistant OSCC is yet to be explored.

In this study, we have performed a CRISPR-based kinome screening in Docetaxel inherent resistant cells and identified NEK9, which plays a significant role in Docetaxel resistance. In *Homo sapiens*, NEKs, Never In Mitosis A (NIMA) related kinases family consists of 11 kinases (NEK1-NEK11) which have structural homology with the NIMA kinase of *Aspergillus nidulans*. Structurally, the unique feature of NIMA proteins involves the presence of an N-terminal catalytic domain that has serine/threonine kinase activity, the only exception is NEK10 having the centrally located catalytic domain (Panchal & Evan Prince, 2023). Other than the catalytic domain, NEKs have coiled–coiled domain and PEST sequences that participate in ubiquitin-dependent proteolysis. All NEKs have a hallmark His-Arg-Asp (HRD) motif in their catalytic domain, which is positively controlled by phosphorylation, they have a serine/threonine residue within the activation loop (Fry *et al*, 2012). Although the functional role of NEKs is poorly characterized, they are reported to have a potential role in cell division. Particularly, NEK9 is an upstream activator of NEK6 and NEK7 and these three NEKs play roles in generating mitotic spindle (Belham *et al*, 2003). The kinase activity of NEK9 is also reported to promote metastasis. In cancer-associated fibroblasts, NEK9 interacts with roundabout guidance receptor 1 and phosphorylates tripartite motif containing 28 (TRIM28) and cortactin (CTTN) to facilitate SLIT2-mediated metastasis (Lu *et al*, 2023). In gastric cancer cells, NEK9 activates RhoA by phosphorylation of Rho/Rac guanine nucleotide exchange factor 2 to regulate cell motility (Lu *et al*, 2021). In this study, we found that selectively knocking out NEK9 sensitizes cancer cells to Docetaxel and the kinase activity of NEK9 is crucial for augmenting resistance against Docetaxel. We found that knocking out NEK9 followed by treatment of Docetaxel induces DNA damage that activates Extracellular signal-regulated kinase, ERK (p-T202/Y204), which in turn induces Gasdermin-E (GSDME) mediated cancer cell pyroptosis in drug-resistant cells. We also identified Fostamatinib as a potent inhibitor of NEK9 activity.

## 2. Results

### 2.1 CRISPR-based kinome screening reveals NEK9 as a novel regulator of Docetaxel resistance in cancer cells

The Docetaxel sensitivity pattern in OSCC lines was determined by performing Docetaxel response assays (Fig. S01A-D). According to the results, SCC4 is the least sensitive line (IC50:5.17nM) while H357 is the most sensitive line (IC50:1.311nM) for Docetaxel. Based on this information, SCC4 was selected for kinome screening. A CRISPR-based kinome-wide screening was performed utilizing a ready-to-infect lentiviral sgRNA library knocking out 840 kinases individually with a total of 3214 sgRNA (up to 4 sgRNA per gene target) to explore for kinase(s) which play a vital role in driving Docetaxel resistance. For this purpose, SCC4 lines stably expressing Cas9 were generated using a lentivirus approach (Fig. S01E) which exhibited drug drug-resistant phenotype comparable to that of the parental SCC4 cell line (Fig. S01F). Additionally, we evaluated the polybrene and puromycin tolerance concentration in Cas9 overexpressing clones (Fig. S01G-H). For primary screening, Cas9 expressing SCC4 lines (Fig. S01E) were infected with lentiviruses, individually knocking out 840 kinases, followed by treatment with sub-lethal dose (2nM) of Docetaxel for 48h, and the cell viability was observed in high content analyzer using fluorescent live/dead dye (Fig. 1A). The screening protocol was optimized using appropriate positive control and negative control as described in our earlier study (Mohanty *et al*., 2022). In brief, SCC4 cells with HPRT knockout showed resistance to drug 6-thioguanine which was kept as positive control throughout the screening (Fig S01I). From primary screening, 316 of 840 kinase genes were selected for further analysis by rejecting 524 for their lethality (cell viability <65%) (Fig. 1B). 316 selected knockouts, when analyzed for cell viability in Docetaxel treated condition, fetched only 53 targets with significant cell death (Cell viability <30%) (Fig. 1B marked in orange shade). For secondary screening, those 53 candidates were knocked out followed by treatment of Docetaxel, and cell viability was analyzed in high content analyzer in four different inherently Docetaxel-resistant cancer cells including OSCC lines SCC4 and SCC9, pancreatic cancer line MIA PaCa-2 and prostate cancer line DU145 (Fig. 1C, S02A-C). MIA PaCa-2 and DU145 cells were selected in secondary screening due to their resistant pattern to Docetaxel (Fig. S02A). Docetaxel sensitivity of Cas9 overexpressing cell lines was estimated and was comparable with the parental lines (Fig. S02D). Puromycin sensitivity and polybrene tolerance in Cas9 overexpressing clones were also evaluated prior to performing secondary screening (Fig. S02E-F). Never in mitosis gene A (NIMA)-related kinase 9 (NEK9) was identified as topmost target in the secondary screen with least cell viability and Dual specificity tyrosine-phosphorylation-regulation kinase 3 (DYRK3) being the second most target (Fig. 1D). Overall, our kinome screening data suggests that NEK9 is a potential kinase for sensitizing four different cancer lines to Docetaxel (Fig. 1D and 1E). Due to higher inherent Docetaxel resistance, most of the further knockout experiments were accomplished in MIA PaCa-2 and DU145 cell lines.

**Fig. 1.**
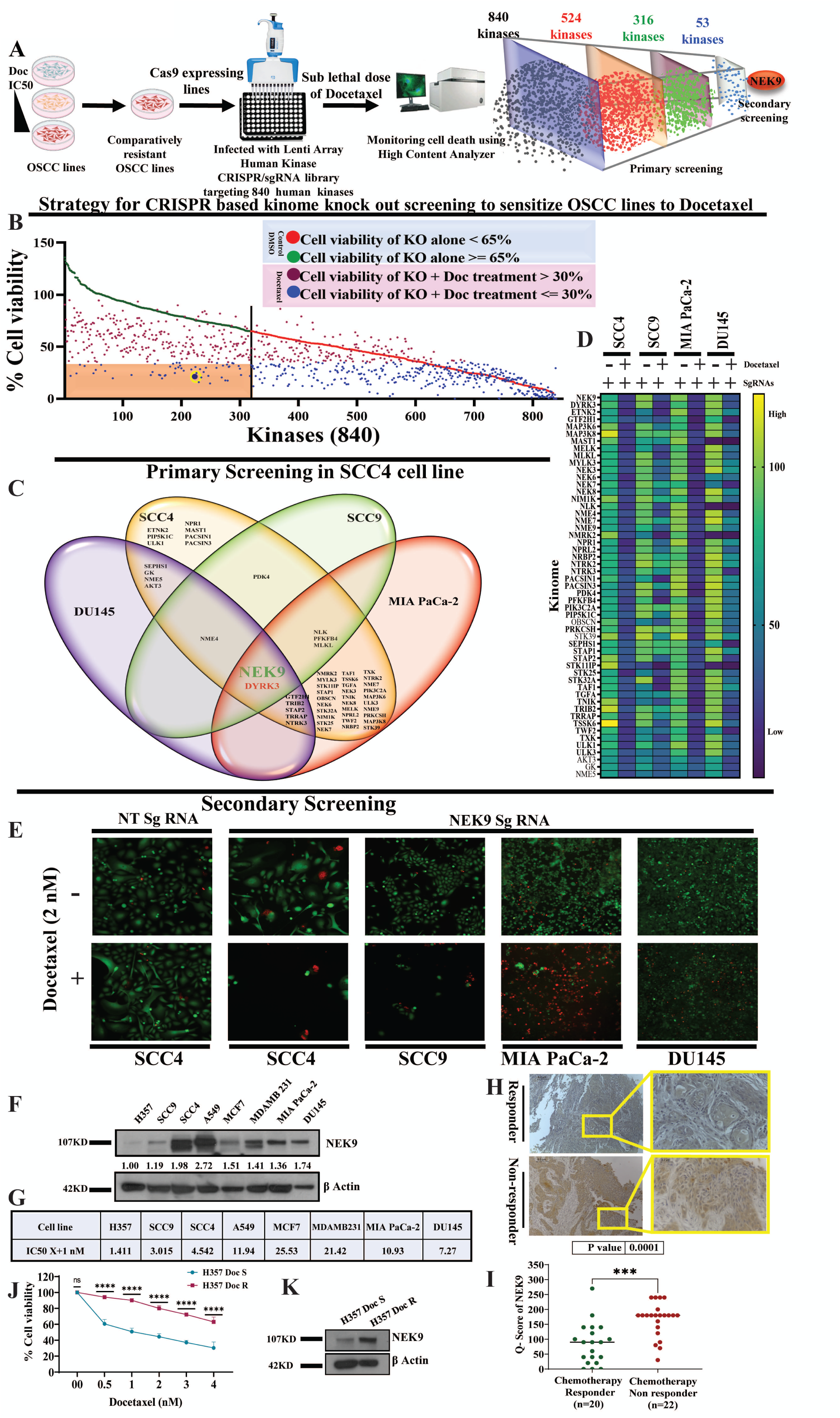
CRISPR-based Kinome screening identified NEK9 as a potential driver of Docetaxel resistance. **A.** Schematic presentation of approach for kinome screening to discover the potential kinase responsible for Docetaxel resistance in OSCC. **B.** Primary CRISPR-Cas9-based kinome screening was performed in the SCC4 cell line to target 840 human kinases. Red and green dots represent the cell viability of each kinase knockout without drug treatment of which red dots represent the rejected lethal knockouts (n=524) with cell viability (< 65%), and green dots (n=316) for cell viability (>65%). Similarly, the magenta and blue dots are the representatives of the same 840 kinase knockouts in Docetaxel treated (sublethal dose 2nM) condition. Kinase knockouts after drug treatment showing cell viability >30% are depicted in magenta colour (n=419). Knockouts that significantly induced cell death (cell viability <30%) upon Docetaxel treatment are depicted in blue dots (n=421). From primary screening, we selected 53 kinases that were not lethal (represented in the orange box) when knocked out and in untreated condition but the cell viability decreased significantly when treated with Docetaxel. **C.** Secondary screening was performed in four indicated cell lines for the top 53 kinases from primary screening of which only two kinases (NEK9 and DYRK3) were common for all the four cell lines. NEK9 was selected as a potential target purely based on the lowest survival fraction among the two candidates. **D.** Heat map shows the % cell viability of 53 kinase knockouts in the presence and absence of Docetaxel for the four cell lines. **E.** Representative images from high content analyzer for NEK9 knockout with and without Docetaxel treatment for all the cell lines, and negative control for SCC4. **F. G.** Protein expression of NEK9 indicated against respective Docetaxel IC50s of different cancer lines. **F** panel represents **t**he band intensity of NEK9 expression and G panel represents the Docetaxel IC50s of indicated cell lines. **H.** Representative IHC staining of NEK9 in OSCC patient tumors (Scale bar: 0.2μM). **I.** IHC scoring for NEK9 from panel (H) (Q Score□=□Staining Intensity□×□% of Staining), (Median, n□=□20 for chemotherapy-responder and n□=□22 for chemotherapy-non-responder) ***P<0.001 by 2-tailed Student’s t-test, **J.** Indicated cells were treated with Docetaxel in dose-dependent manner for 48h and cell viability was determined by MTT assay (n=3 and 2-way ANOVA, ****P<0.0001). **K.** NEK9 expression was analyzed in H357 Docetaxel resistant (H357 DocR) Vs its sensitive counterpart (H357 DocS) by immunoblotting (n=3).

### 2.2 NEK9 expression correlates with Docetaxel resistance of diverse cancer cell lines and OSCC tumors

In order to investigate the potential correlation between NEK9 expression and Docetaxel sensitivity, we conducted an MTT experiment and immunoblotting. Our results indicate that the expression of NEK9 in several cancer cell lines is positively correlated with their resistance to Docetaxel (Fig 1F and 1G), with the exception of MCF7. In order to illustrate the clinical significance of our discoveries, we examined the expression of NEK9 in clinical samples obtained from OSCC patients who had undergone the TPF regimen (neoadjuvant chemotherapy, NACT). The IHC analysis revealed significantly elevated NEK9 expression in tumor tissue samples from chemotherapy non-responders (n=22) compared to responders (n=20) OSCC patients (Fig. 1H and 1I). To explore the NEK9 expression status in the acquired Docetaxel resistance model, we generated Docetaxel-resistant OSCC cell lines with consistent exposure to sublethal doses of Docetaxel for a prolonged period of time (Fig. 1J). The immunoblotting data suggests that NEK9 expression is significantly elevated in Docetaxel-resistant cells (H357 DocR) as compared to their sensitive counterpart (H357 DocS) (Fig 1 K).

### 2.3 Docetaxel resistance can be overcome by depleting NEK9 in cancer cells

To confirm the potential role of NEK9 in Docetaxel resistance, NEK9 stable knockout clones were generated with two different sgRNA using a lentiviral approach (Fig. S03A). MTT and colony-forming assay suggest that cell viability and proliferation were significantly reduced when NEK9-depleted cells were exposed to Docetaxel (Fig 2A and S03B). We also found enhanced cell death in NEK9-depleted cells in response to Docetaxel treatment (Fig S03C). Similarly, elevated expression of cleaved PARP and activated H2AX was observed in NEK9-depleted cells treated with Docetaxel suggesting induction of cell death (Fig 2B). Furthermore, NEK9 knockout sensitized PDC1 cells to Docetaxel, which were isolated from a TP chemotherapy-treated non-responder OSCC patient (Maji *et al*, 2019) (Fig. 2A, 2B, S03B-C). PARP cleavage and activated H2AX in SCC4 NEK9 KO cells upon drug treatment revealed similar phenotypes (Fig. S03D). Similar results were obtained when NEK9 was stably knocked down in DU145 cell lines (Fig. S03E-H). Furthermore, the appearance of compact microtubule bundling in NEK9-depleted cells upon drug treatment indicated efficient Docetaxel impact on cells (Fig. 2C). To evaluate if our *in-vitro* data holds true in *in-vivo* experiments, xenograft tumors using NEK9 WT and NEK9 KO PDC1 cells were generated in the flank of nude mice and the mice were subjected to Docetaxel treatment (5mg/kg) twice per week. Treatment with Docetaxel (5mg/kg) significantly reduced the tumor burden in the NEK9 KO group but not in the NEK9 WT group (Fig. 2D-F). Next, we also used the Zebrafish (*Danio rerio*) [Tg(fli1: EGFP)] tumor xenograft model to further validate our findings. Here, we found that tumor growth was significantly reduced in the NEK9 KO group when treated with Docetaxel (5μM) (Fig. 2G and 2H). The zoomed images revealed reduced cancer cell migration in NEK9-depleted groups. These data imply that NEK9 imparts Docetaxel resistance and that ablating NEK9 sensitizes cancer cells to Docetaxel.

**Fig. 2.**
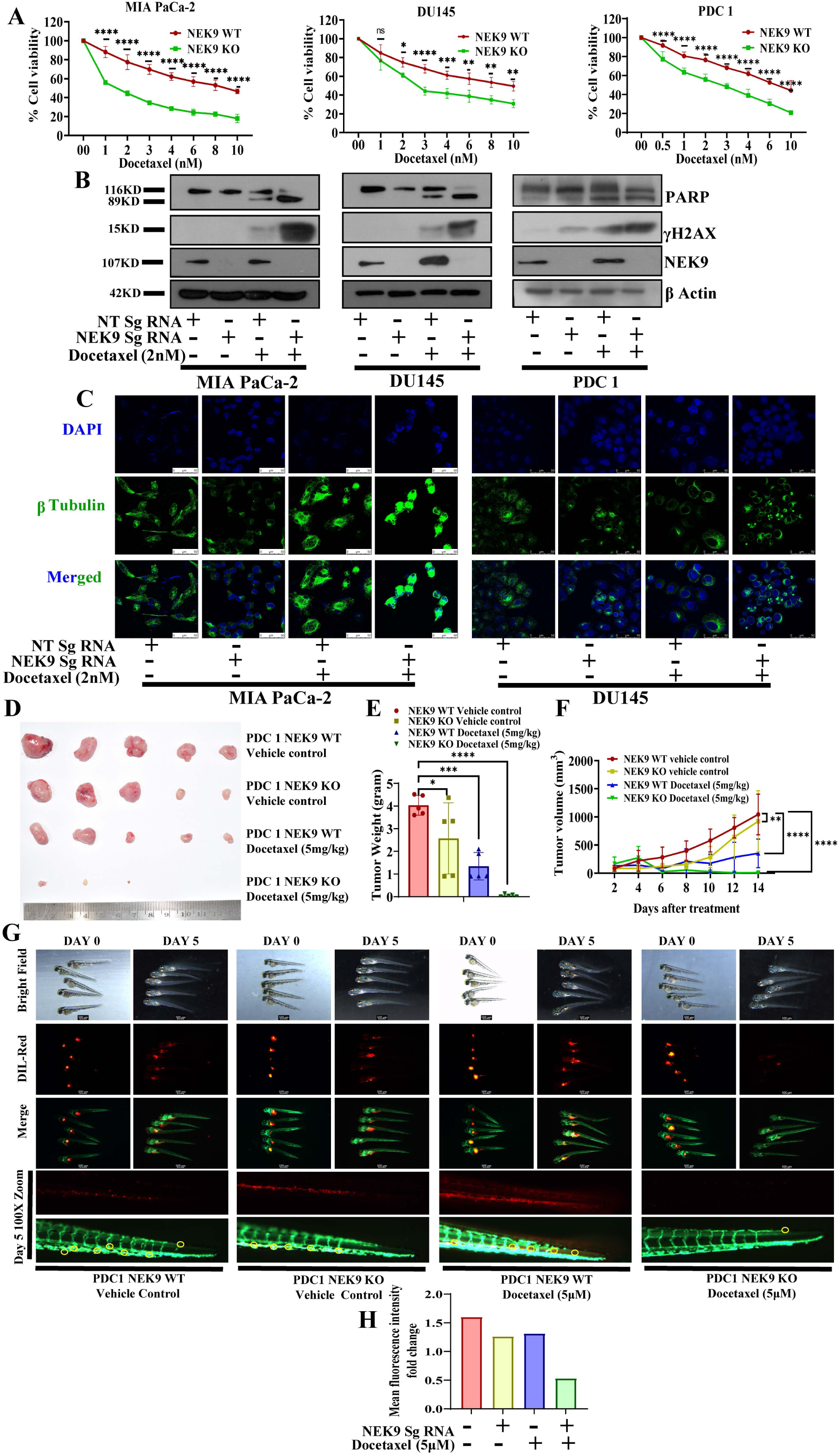
Ablation of NEK9 sensitizes cancer cells to Docetaxel. **A.** Indicated cells were treated with Docetaxel for 48h in a dose-dependent manner and cell viability was determined by MTT assay (n=3 and 2-way ANOVA, ****P<0.0001). **B.** Indicated cells were treated with Docetaxel or vehicle control for 48h. Cell lysates were collected and immunoblotting was performed with indicated antibodies. (n=3). **C.** Tubulin bundling was analyzed in the indicated cells after 48 hrs of Docetaxel treatment by immunostaining for β tubulin. These results are the representatives of at least three independent experiments performed with duplicates. **D**. Patient-derived cells (PDC1) from chemotherapy (TP) nonresponder OSCC patient tumor was established earlier. PDC1 NEK9 WT and NEK9 KO cells were implanted subcutaneously in athymic male nude mice, followed by intraperitoneal Docetaxel (5mg/kg) for the indicated time period. At the end of the experiment, mice were sacrificed and tumors were isolated and photographed (n=5). **E**. The bar diagram presents the tumor weight of each indicated group at the end of the experiment (mean±SEM, n=5, Ordinary one-way ANOVA, ****P<0.0001). **F**. Tumor growth was measured at the indicated time points using a digital vernier caliper and plotted as a graph (mean±SEM, n=5, 2-way ANOVA, ****P<0.0001). **G.** Lateral view of fluorescent transgenic [Tg(fli1:EGFP)] zebrafish embryos at Day 0 and Day 5 injected with Dil-Red stained PDC1 control and NEK9 KO cells with and without treatment of Docetaxel. The lower row indicates the Zoomed image of the distal part of the embryo (5 days post-injection) to monitor the migration of tumor cells. **H.** The tumor growth was assessed by an increase or decrease in fluorescence intensity on day 5 compared to day 0. The quantitation of fluorescence intensity was performed using ImageJ software and represented a fold change in fluorescence intensity.

### 2.4 Ectopic NEK9 expression augments Docetaxel resistance in cancer cells

To confirm the potential role of the kinase activity of NEK9 in imparting Docetaxel resistance, we generated NEK9 WT (pMRX-IPU-FLAG-NEK9) and NEK9 kinase dead domain (pMRX-IPU-FLAG-NEK9T210A) overexpressing stable H357 (OSCC cell line) cell lines using Retroviral approach (Fig. 3A). The overexpression experiments were performed in H357 cell lines because of its low NEK9 basal expression and higher Docetaxel sensitivity. Cell viability and apoptosis data suggest that overexpression of WT NEK9 but not Kinase mutant protein boosts Docetaxel-resistant phenotype (Fig. 3B and 3D). Immunoblotting of cleaved PARP and p-H2AX suggests that Docetaxel-induced cell death is significantly reduced with NEK9 WT overexpression but not with the NEK9 kinase dead domain (Fig. 3C). Ectopic WT NEK9 expression also enhanced the colony forming ability of drug-treated cells (Fig. 3E). Furthermore, the compactness of microtubule bundles was lost when WT NEK9 overexpressing H357 cells were treated with Docetaxel. On the other hand, when the H357 control and NEK9 kinase mutant cells when treated with Docetaxel, they retained the bundling morphology (Fig. 3F). We also transiently overexpressed NEK9 WT and kinase mutant genes in NEK9 KD (DU145 and SCC9) cells. NEK9 knockdown-induced cell death as evident from cleaved PARP expression was rescued with WT NEK9 but not kinase mutant overexpression, suggesting rescue of Docetaxel resistance by NEK9 kinase activity (Fig. 3G). These findings support the possible involvement of NEK9 kinase activity in Docetaxel resistance.

**Fig. 3.**
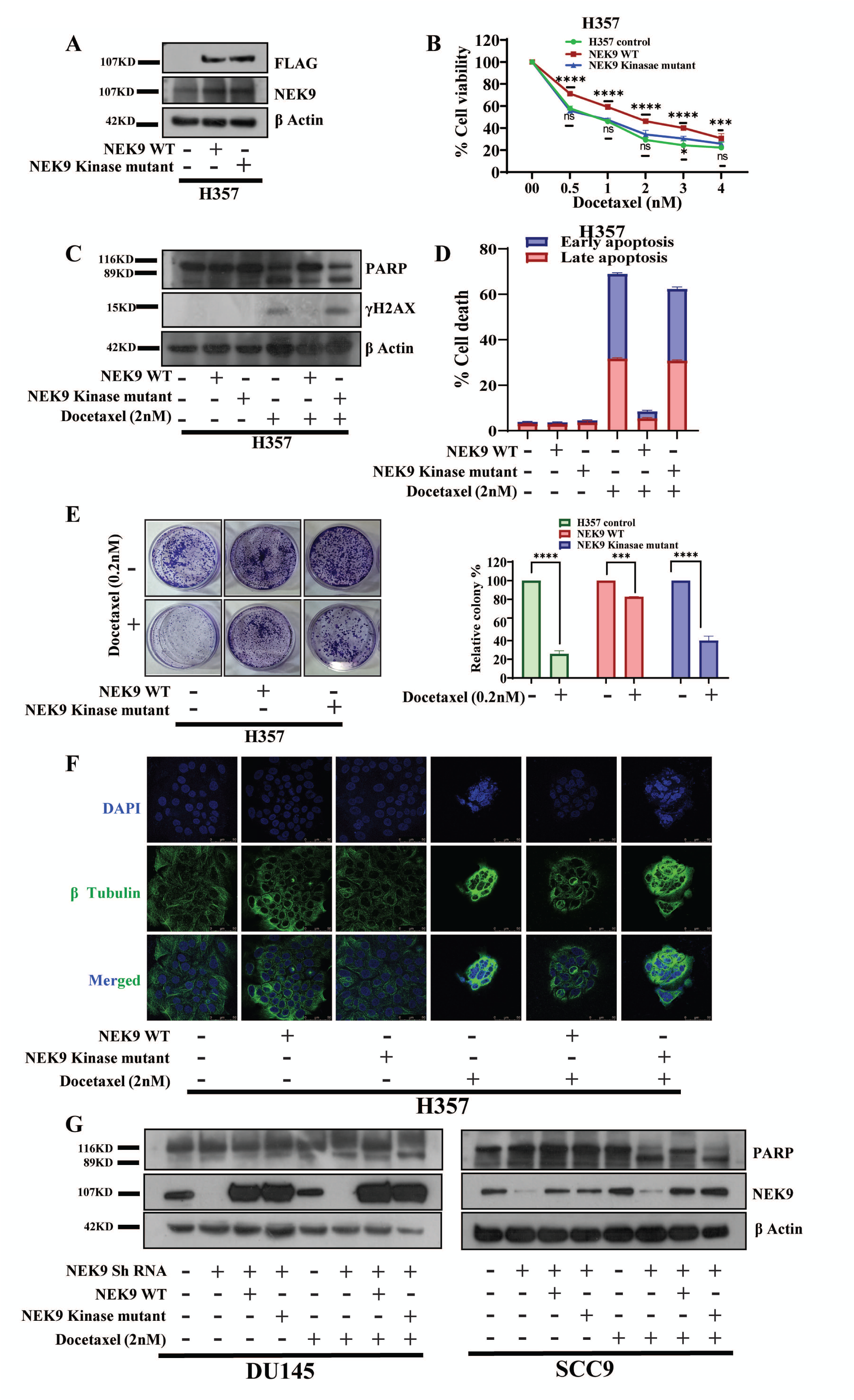
NEK9 kinase activity is the driver of Docetaxel resistance. **A.** NEK9 WT (pMRX-IPU-FLAG-NEK9) and NEK9 kinase mutant (pMRX-IPU-FLAG-NEK9T210A) constructs were stably overexpressed in H357 cell line using retroviral approach and the overexpression was confirmed by immunoblotting against indicated antibodies. **B.** Cell viability was assessed in the indicated cell lines by MTT assay after 48 hrs of Docetaxel treatment with the indicated dose (n=3 and 2-way ANOVA, ****P<0.0001). **C.** H357 cells stably expressing NEK9 WT or NEK9 kinase mutant were treated with vehicle control or Docetaxel for 48h and lysates were collected to perform immunoblotting with indicated antibodies (n=3). **D.** H357 cells stably expressing NEK9 WT or NEK9 kinase mutant were treated with vehicle control or Docetaxel for 48h followed by which cell death was monitored by annexin V/7AAD assay (n=3). **E.** Representative images of colony-forming assay for indicated cells treated with 0.2nM Docetaxel for 12-14 days, the bar diagram depicts the relative percentage of colonies (n=2 and 2-way ANOVA, ****P<0.0001). **F.** Immunofluorescence showing tubulin bundling in H357 cells stable expressing NEK WT or NEK9 kinase mutant treated with Docetaxel for 48h. **G.** NEK9 was stably knocked down using ShRNA against 3’UTR of NEK9. NEK9 WT (pMRX-IPU-FLAG-NEK9) and NEK9 kinase mutant (pMRX-IPU-FLAG-NEK9T210A) genes were transiently transfected in the indicated KD cells followed by Docetaxel treatment. Immunoblotting was performed with the indicated antibodies.

### 2.5 NEK9 depletion sensitizes cancer cells to Docetaxel via induction of pERK1/2 mediated Cancer Cell Pyroptosis

To explore the role of NEK9 as a kinase in driving Docetaxel resistance, we performed phosphorylation profiling with a Human Phospho Kinase array using whole cell lysates isolated from Docetaxel treated and untreated DU145 NEK9 WT and KO cells. This Phospho kinase array detects relative levels of phosphorylation of 37 kinase phosphorylation sites and 2 related total proteins (Fig. 4A). We discovered ERK1/2 phosphorylation was enhanced at T202/Y204 position when NEK9 depleted cells were treated with Docetaxel (Fig. S04A, 4A and 4B). We confirmed our finding in DU145 and MIA PaCa-2 cell lines through immunoblotting (Fig. 4C). During our phenotypic studies, pyroptosis-like morphology with cytoplasmic swelling were observed under the microscope when NEK9 ablated cells were exposed to Docetaxel (Fig. S05B) which was furthermore distinct from SEM imaging of the same (Fig. 4D). Existing literature suggests the association of activated ERK1/2 with Cisplatin-induced pyroptosis (Fan *et al*, 2023). On the other hand, NEK7, a downstream interacting partner of NEK9 is a very well-known regulator of NLRP3-mediated pyroptosis (Shi *et al*, 2015). We hypothesized from the phospho array data that the increased phosphorylation on ERK1/2 upon Docetaxel exposure might induce pyroptosis in NEK9-depleted cells. The status of pyroptosis was first checked by measuring the presence of LDH in cell supernatants which is one of the hallmarks of pyroptosis. Our LDH assay data suggests that LDH release was significantly increased in NEK9 KO cells upon drug exposure (Fig. 4E). Next, we checked if NEK9 depletion and exposure to Docetaxel induces Immune Cell Pyroptosis (ICP) in cancer cells. Hence, we performed immunoblotting against ICP markers like cleaved caspase-1, cleaved Gasdermin-D (GSDMD) from the lysate, and mature IL1β from the supernatant. Our data revealed no clear ICP in NEK9-depleted cancer cells treated with Docetaxel (Fig. S05A). THP1 monocytic cells stimulated with LPS in combination with Nigericin were used as a positive control. So, next, to confirm the other possible pathway that is leading to pyroptosis, we checked for Cancer Cell Pyroptosis (CCP) markers like Caspase 3 and GSDME cleavage. From the immunoblotting data, it was clear that Docetaxel acts as an external stimulus to induce pyroptosis via the Caspase 3-GSDME axis in the absence of NEK9 (Fig. 4F). Similar results were obtained for NEK9 depletion in SCC4 and PDC1 cells (Fig. S05B-D). We also found GSDME oligomerization (which is required for pore formation in the cell membrane) in NEK9-depleted cells upon drug treatment (Fig. 4G).

**Fig. 4.**
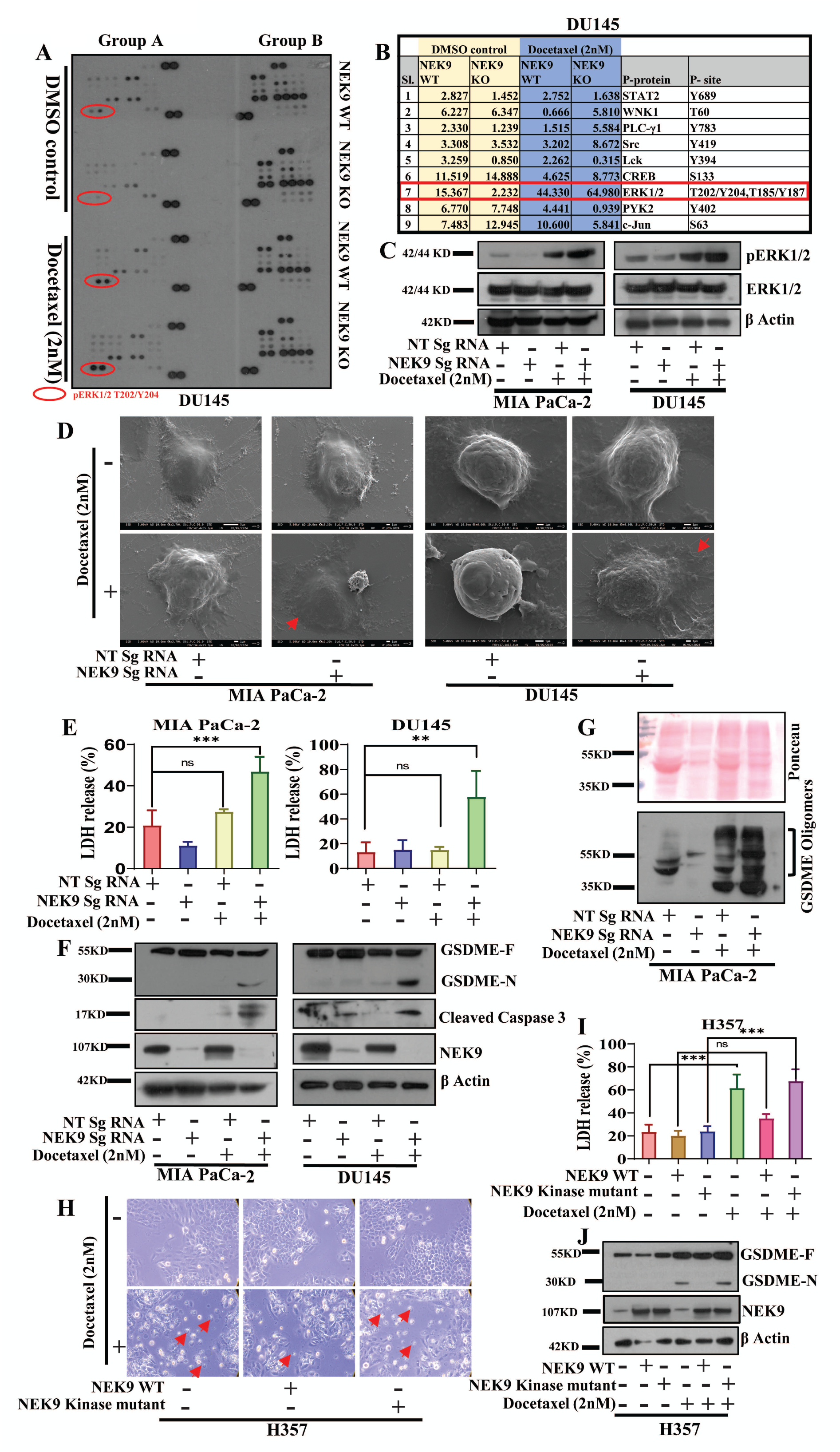
NEK9 depletion regulates pyroptosis through phosphorylation of ERK1/2 T202/Y204. **A.** Lysates from DU145 NEK9 WT and KO cells prior treated for 48 hrs with Docetaxel were subjected to Human Phospho Kinase array, to detect relative levels of phosphorylation of 37 kinase phosphorylation sites and 2 related total proteins. The change in expression pattern of ERK1/2 is indicated in red circles. **B.** Relative signals for the phospho-proteins from the indicated cells. **C.** Indicated cells were treated with vehicle control and Docetaxel for 48h and immunoblotting was performed against indicated antibodies. **D**. Morphological changes in NEK9 KO cells upon Docetaxel treatment. The arrow indicates cell swelling under SEM. **E**. LDH release % was measured in the indicated cells after 48 hours of drug treatment (n=3, Ordinary one-way ANOVA, ***p<0.001). **F.** NEK9 WT and KO cells were treated with Docetaxel for 48 hours and immunoblotting was performed with the indicated antibodies. **G.** The upper panel shows the Ponceau stained blot and the lower panel depicts the immunoblot analysis for GSDME oligomerization from insoluble fraction of Docetaxel-treated MIA PaCa-2 WT and KO cells. **H.** Morphological changes in NEK9 KO cells upon Docetaxel treatment. The arrow indicates cell swelling. **I**. LDH release % detection in H357 cells stably expressing NEK9WT or NEK9 kinase dead cells treated with Docetaxel for 48h (n=3, Ordinary one-way ANOVA, ***p<0.001). **J**. Lysates were isolated from the indicated cells treated with Docetaxel for 48h and were subjected to immunoblotting with the indicated antibodies.

### 2.6 NEK9 kinase overexpression abates the Docetaxel-induced pyroptosis

To know if NEK9 kinase activity is important for Docetaxel-induced CCP, we scored pyroptotic markers in H357 cells ectopically expressing WT NEK9 and NEK9 kinase mutant proteins. Our bright field microscopy and LDH release data suggest that there is a marked reduction of Docetaxel-induced CCP in cells stably expressing WT NEK9 but not in NEK9 kinase mutant (Fig. 4H and 4I). We also found reduced cleaved GSDME in H357 cells ectopically expressing WT NEK9 as compared to NEK9 kinase mutant (Fig. 4J). To further confirm the role of GSDME in Docetaxel-induced pyroptosis, we generated stable clones of GSDME knockouts (KO) in H357 cells. To our surprise, the depletion of GSDME did not halt the pyroptotic cell death upon drug treatment (Fig. 5A and 5C). We reassessed the ICP marker GSDMD status to dig into the reason behind pyroptosis in GSDME KO cells. From immunoblotting, it was apparent that GSDMD compensates for GSDME in the absence of later to induce pyroptosis upon drug exposure (Fig. 5B). However, the mechanism of GSDMD compensation in the absence of GSDME remains unsolved, we plan to identify the mechanistic pathway for this in our subsequent study. Since our phospho array data suggests ERK to be downstream of NEK9, we also investigated whether ERK1/2 contributes to Docetaxel response in cancer cells. For this, we used Bradykinin (ERK1/2 inducer) (Mukhin *et al*, 2003) in combination with Docetaxel in NEK9 WT cells and checked if the external activation of ERK1/2 induces pyroptotic cell death. Reduced cell viability with enhanced LDH release in the case of the combined treated group confirmed our assumption that ERK1/2 activation led to the induction of pyroptosis (Fig. 5D and 5E). We next checked the pyroptosis status in tissue sections of our *in-vivo* PDX tumors, which showed cleavage of Caspase3 and GSDME in NEK9 KO tumors upon Docetaxel treatment (Fig. 5F). It is earlier reported that drug-induced DNA damage in cancer cells could lead to activation of ERK (Tang *et al*, 2002; Yan *et al*, 2008). Hence, to access the DNA damage upon Docetaxel treatment to NEK9-depleted cells, we performed COMET assay. The COMET tail length was found to be significantly elongated in NEK9-depleted cells upon low dose of Docetaxel treatment (Fig. 5G and 5H).

**Fig. 5.**
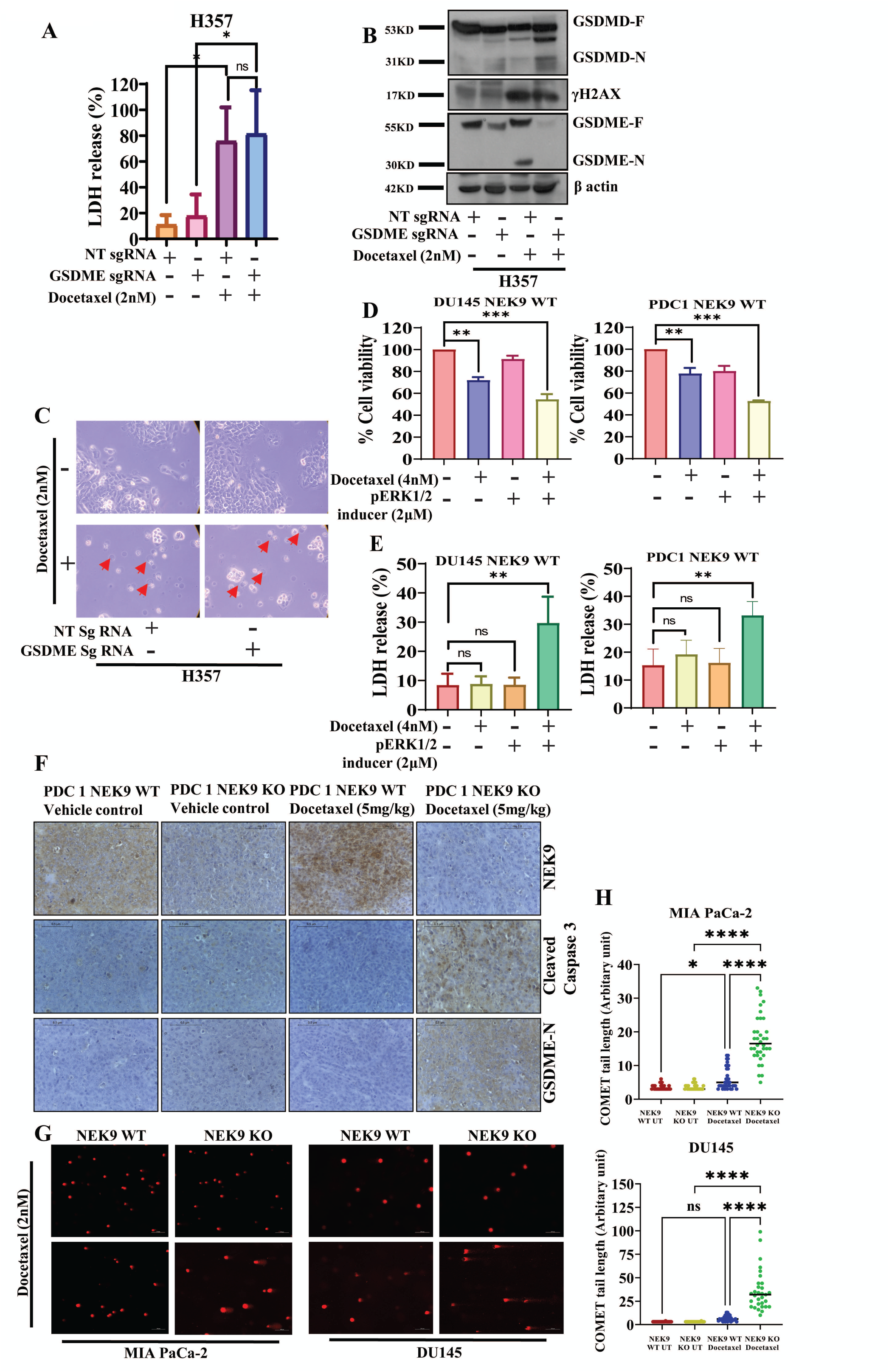
NEK9 depletion induces cancer cell pyroptosis with external stimuli Docetaxel. **A.** GSDME was stably knocked out in H357 cells and LDH release % was analyzed followed by treatment with Docetaxel for 48h (n=3, Ordinary one-way ANOVA, *p<0.05). **B.** Immunoblotting with indicated antibodies was performed in H357 GSDME WT and KO cells treated with Docetaxel for 48h. **C.** Morphological changes in H357 GSDME WT and KO cells upon Docetaxel treatment. The arrow indicates cell swelling. **D.** NEK9 WT cells were treated with pERK1/2 inducer bradykinin (2μM) and Docetaxel (4nM) for 48 hrs and cell viability was analyzed with MTT assay (n=3, Ordinary one-way ANOVA, **p<0.01, ***p<0.001). **E.** LDH release % detection in the NEK9 WT cells after treatment with indicated drugs as mentioned in panel C (n=3, Ordinary one-way ANOVA, **p<0.01). **F.** Paraffin-embedded tumor sections from experiments mentioned in panel 3D were subjected to IHC with the indicated antibodies. Scale bar: 0.5μM. **G. H.** NEK9 WT and KO cells subjected to Docetaxel treatment (2nM) were analyzed for DNA damage with comet assay (Ordinary one-way ANOVA, ****p<0.0001).

### 2.7 Identification and characterization of Fostamatinib as an NEK9 inhibitor to target Docetaxel resistant cancer

Since very limited information is available about NEK9 inhibitors in the literature, we searched for prospective NEK9 inhibitors. From the DrugBank online database (https://go.drugbank.com) (Wishart *et al*, 2006), where we found information on a single possible potential inhibitor Fostamatinib. From the literature, we also found Dabrafenib as another small molecule inhibitor targeting NEK9 (Phadke *et al*, 2018). We evaluated the kinase activity of NEK9 in the presence of dabrafenib and Fostamatinib individually. The *in-vitro* kinase assay data suggest the potency of both the molecules to have high NEK9 inhibitory activity, with both inhibiting 50% kinase activity at 500nM concentration (Fig 6A and 6B). Dose-dependent cell viability assay was performed to select the non-toxic concentration of NEK9 inhibitors in different cancer lines. Both dabrafenib and Fostamatinib were non-toxic (cell viability> 80% even at 5μM) when treated alone to cancer lines (Fig. 6C). Since Fostamatinib was less toxic as compared to dabrafenib, we further chose the former one for combination treatment with Docetaxel. In the combinatorial study, we found that Fostamatinib efficiently sensitizes cancer lines to Docetaxel (Fig. 6D). Fostamatinib failed to further sensitize NEK9 KO cells to Docetaxel, confirming its specificity via NEK9 kinase activity inhibition (Fig. 6E). In harmony with our NEK9 KO observation, we found that the novel combination of Docetaxel and Fostamatinib enhanced phosphorylation of ERK1/2 at T202/Y204 position and induced pyroptotic cell death via cleavage of GSDME (Fig. 6F and 6G). To check the *in-vivo* efficacy of this novel combination, we generated PDX tumors in nude mice using PDC1 cells and treated them with Docetaxel (5mg/kg) twice weekly and Fostamatinib (0.5mg/kg) once every two days for two weeks. The data suggest that there is a profound reduction of tumor burden in the combination group of both agents as compared to single treatments of either agent (Fig. 7A, 7B, and 7C). Immunohistochemistry data revealed the reduction in tumor burden was possibly due to induced pyroptosis in the combination treatment group (Fig. 7D). Lastly, we confirmed our *in-vivo* data using a zebrafish patient-derived xenograft model and we observed a reduction of tumor burden when Docetaxel was treated with a combination of Fostamatinib (Fig. 7E and 7F).

**Fig. 6.**
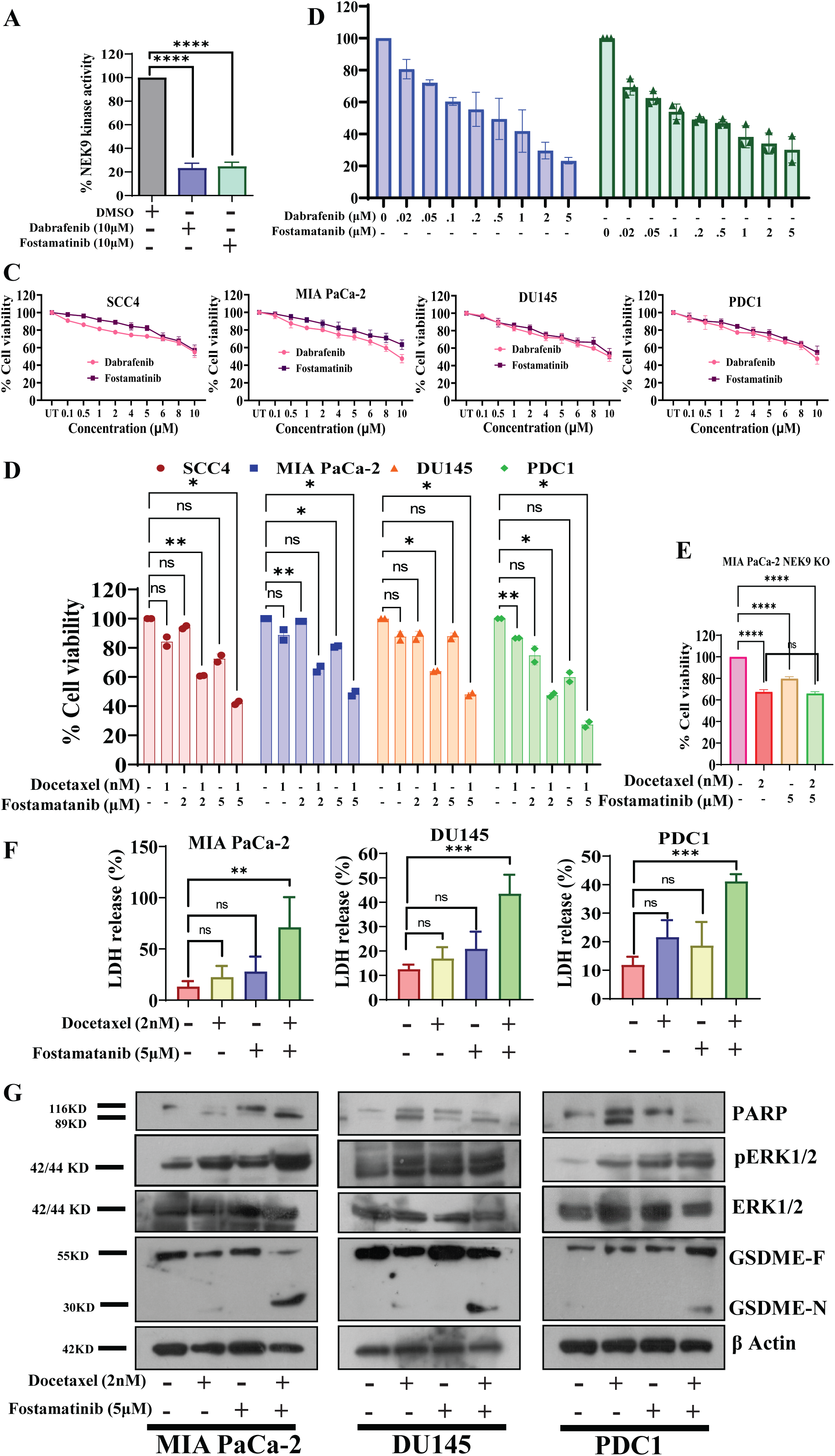
Evaluation of Fostamatinib as a potent NEK9 inhibitor to sensitize cancer cells to Docetaxel-mediated pyroptosis. **A.** In vitro NEK9 kinase activity assay was performed using two different compounds dabrafenib (10μM) and Fostamatinib (10μM) and kinase activity was analyzed by ADP-Glo™ according to manufacturer’s protocol (n=3, Ordinary one-way ANOVA, ****p<0.0001). **B.** Determination of EC50 value for kinase activity of the inhibitors (n=3 and 2-way ANOVA, **p<0.01, ****P<0.0001). **C.** Indicated cells were treated with either Fostamatinib or dabrafenib for 48h and cell viability was determined by MTT assay (n=3). **D.** MIA PaCa-2 NEK9 KO cells were treated with indicated concentrations of drugs in singlets or combination for 48 hand cell viability was accessed by MTT assay (n=3, Ordinary one-way ANOVA, ****P<0.0001). **E.** Indicated cancer cell lines were treated with Docetaxel (1nM) and Fostamatinib (2 or 5μM) in combination for 48 hours and cell viability was determined by MTT assay (n=2 and 2-way ANOVA, *p<0.05, **P<0.01). **F.** Indicated cancer cell lines were treated with Docetaxel (2nM) and Fostamatinib (5μM) in combination for 48 hours. And LDH release percentage was determined (n=3, Ordinary one-way ANOVA, **p<0.01, ***p<0.001). **G.** Lysates from indicated cells were treated with Docetaxel in combination with Fostamatinib for 48h and cell lysates were subjected to immunoblot analysis with indicated antibodies.

**Fig. 7.**
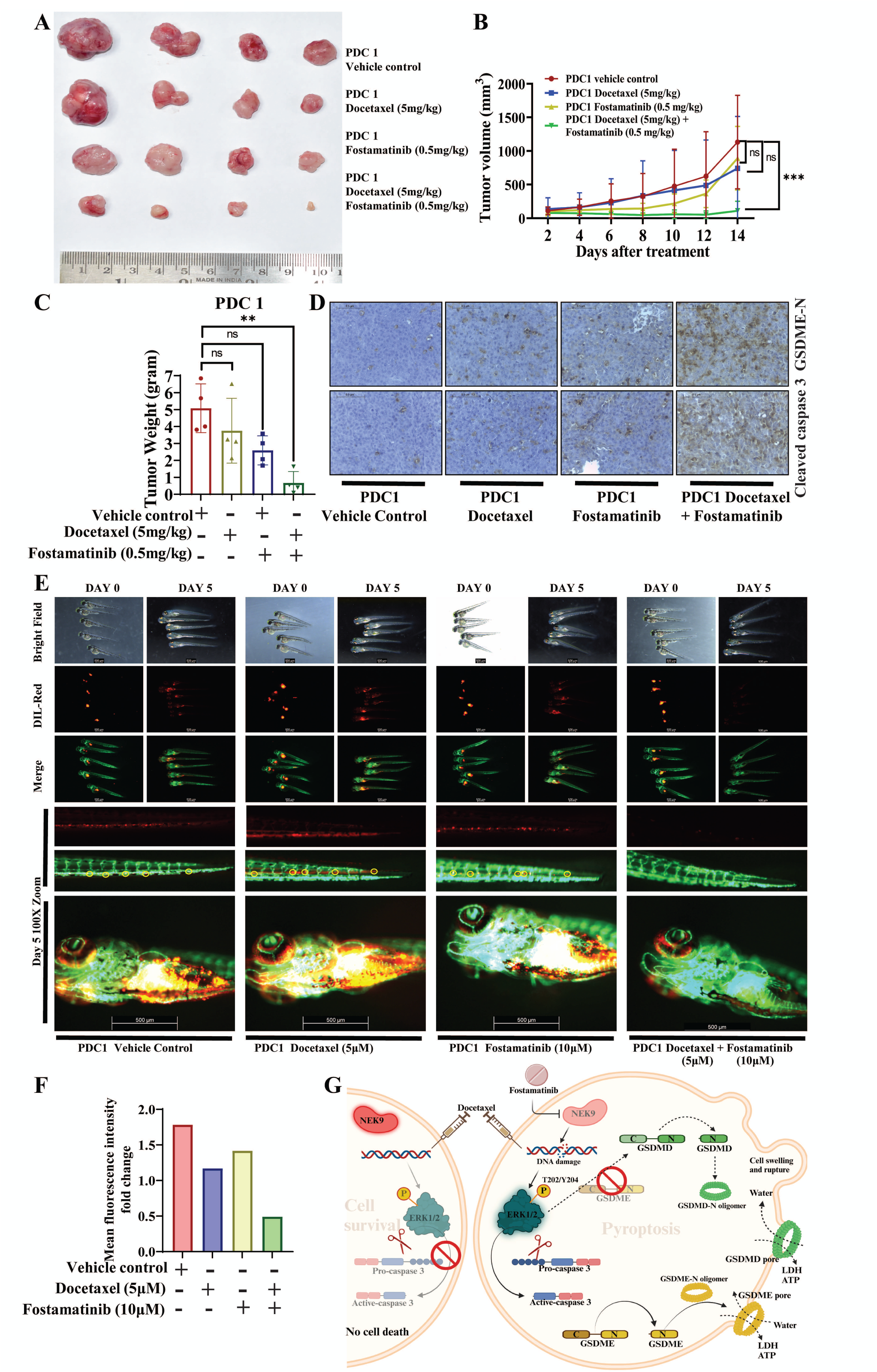
Fostamatinib and Docetaxel synergistically reduce tumor burden in in vivo drug-resistant OSCC. **A.** PDC1 cells were implanted subcutaneously in athymic male nude mice, followed by intraperitoneal Docetaxel (5mg/kg) and Fostamatinib (0.5mg/kg) injection for the indicated time period. Tumors were isolated, arranged and photographed (n=4). **B.** Tumor weight was measured at the end of the experiment and is indicated in the bar diagram (mean±SEM, n=4, Ordinary one-way ANOVA, **p<0.01). **C.** Tumor growth was monitored regularly and was measured every consecutive day using a digital vernier caliper and the growth of each group is depicted in the graph (mean±SEM, n=4, 2-way ANOVA, ***p<0.001). **D.** Paraffin-embedded tumor sections were subjected to IHC with the indicated antibodies. Scale bar: 0.5μM. **E.** Lateral view of fluorescent transgenic [Tg(fli1:EGFP)] zebrafish embryos at Day 0 and Day 5 injected with Dil-Red stained PDC1 cells with combined treatment of Docetaxel and Fostamatinib and their control groups. The bottom panel shows zoomed images of the proximal and distal parts of the embryo (5 days post-injection) to monitor the migration of tumor cells. **F.** The tumor growth was assessed by an increase or decrease in fluorescence intensity on day 5 compared to day 0. The quantitation of fluorescence intensity was performed using ImageJ software and represented a fold change in fluorescence intensity. **G.** Schematic presentation of NEK9-driven Docetaxel resistance and its sensitization via pERK1/2 mediated cancer cell pyroptosis in the absence of NEK9 (Created with BioRender.com).

## 3. Discussion

The addition of Docetaxel to Cisplatin and 5FU (PF) regimen in a phase-2 clinical trial significantly improved the overall survival and prolonged the progression-free survival as compared to PF in advanced HNSCC patients (Vermorken *et al*, 2007). Hence, TPF is the standard chemotherapy regimen both in the form of adjuvant and NACT in the case of OSCC. Docetaxel predominantly induces apoptosis in tumor cells by regulating not only the dynamics of microtubules but also expression of Bcl-2 family members(Mhaidat *et al*., 2007; Yvon *et al*, 1999). Nevertheless, the literature also suggests that Docetaxel can induce other form of cell death including senescence, mitotic catastrophe, and lytic necrosis (Hernandez-Vargas *et al*, 2007). However, until now it is not clear how Docetaxel induces non-apoptotic modes of cell death. Pyroptosis is a tightly regulated process that comprises pore formation in the cell membrane leading to the release of pro-inflammatory factors that induce prominent immune responses. Pyroptosis can be broadly divided into two different types i.e. immune cell pyroptosis (ICP) and cancer cell pyroptosis (CCP)(Hou *et al*, 2021). In the case of ICP, intracellular pathogens can be detected in inflammasome complexes to induce pyroptotic cell death and release pro-inflammatory factors to recruit phagocytes to kill these cells. The most notable example is a canonical pathway, where the inflammasome complex activates caspase-1 which cleaves the GSDMD and processes the mature IL-1β/IL-18. The cleaved GSDMD-N terminal domain oligomerizes to form pores in the cell membrane that allow the mature IL-1β/IL-18 to be released into the extracellular region (Shi *et al*., 2015). Non-canonically, activated caspase 11 by bacterial LPS and activated caspase-8 by TNFα can also cleave GSDMD to induce pyroptosis in immune cells (Demarco *et al*, 2020; Kayagaki *et al*, 2011).

In cancer cell pyroptosis, GSDME is cleaved by chemotherapy-induced activated caspase-3 which results in the release of LDH. The mutant GSDME (at D267A and D270A) which lacks a caspase-3 binding site did not show cleavage of GSDME and hence no pyroptosis was observed (Wang *et al*, 2017). Similarly, the combinatorial effect of MEK and BRAF inhibitors augmented the caspase-3-dependent cleavage of GSDME which resulted in the induction of pyroptosis. This led to the release of HMGB1 release into cell supernatant that elicited anti anti-tumor immune response (Erkes *et al*, 2020). In this study, using a CRISPR-based kinome screening, we found that NEK9 kinase is a key player for Docetaxel resistance in cancer cells. Our observation revealed that the depletion of NEK9 triggers DNA damage upon treatment with Docetaxel, which activates ERK(p-T202/Y204). This activation ultimately leads to Gasdermin-E-mediated Cancer-Cell-Pyroptosis. Remarkably, when we knocked out GSDME, we observed GSDMD-mediated pyroptosis in NEK9 exhausted cells when treated with Docetaxel (Fig. 7G). Although we did not investigate the mechanism by which GSDMD compensates for GSDME, we intend to identify a potential mechanistic pathway for this in a subsequent study.

Our subsequent inquiry was to understand how NEK9 depletion activates ERK1/2 upon drug exposure. From the literature survey, we found that NEK9 depletion induces replication stress hypersensitivity and impairment in recovery from replication arrest (Smith *et al*, 2014). Wu *et al*, showed that PLK1 kinase inhibitor enhances the chemosensitivity of Cisplatin by inducing DNA damage and pyroptosis (Wu *et al*, 2019). Intriguingly, PLK1 is a recognized upstream activator of NEK9 (Bertran *et al*, 2011). Hence, our assumption was that PLK1 inhibitor-induced pyroptosis may have been the consequence of inactivated NEK9 in the system. On the other hand, apart from its well-known microtubule stabilization effect, Docetaxel is known to induce DNA damage-mediated cell death (Chao & Goodman, 2021). Several reports suggest that drug-induced DNA damage in cancer cells leads to the activation of ERK1/2 (Tang *et al*., 2002; Yan *et al*., 2008). Hence, we anticipate that when NEK9-depleted cells were treated with Docetaxel, DNA damage is induced which activates ERK1/2 ultimately driving cancer cell pyroptosis. Since Smith *et al*., in 2014 reported that NEK9 is required for recovery from replication arrest, we adopt that inhibiting ERK or caspase-3 that comes into play after DNA damage may not impart changes to the triggered fate of cell death.

Small molecule Fostamatinib is a pro-drug that gets converted to active R406 by the enzyme alkaline phosphatase in the intestinal mucosa. R406 is a potent inhibitor of spleen tyrosine kinase (Syk (Braselmann *et al*, 2006). In addition to Syk, Fostamatinib is also known to inhibit other kinases like Flt3, aurora A kinase, and RET (Clemens *et al*, 2009). Clinical trial suggests that Fostamatinib is well tolerated in patients having mild to moderate adverse effects like diarrhea, hypertension, nausea, dizziness, and an increase in alanine transaminase ALT. These adverse effects were comfortably managed with appropriate drugs (Bussel *et al*, 2018). The *in-vitro* study suggests that Fostamatinib can induce differentiation and block the clonogenic potential of Acute myeloid leukemia (AML) cells through the MEK/ERK1/2 pathway and STAT5A transcription factor (Polak *et al*, 2020). In mice models for pancreatic ductal adenocarcinomas (PDAC), combinatorial treatment of Fostamatinib and gemcitabine rewires the tumor microenvironment by augmenting immunostimulatory macrophages and enhancing CD8+ T-cell responses. (Rohila *et al*, 2023). However, a phase 2 clinical trial with single agent Fostamatinib suggests limited anti-tumor activity in patients with advanced colorectal (CRC), thyroid, non-small-cell lung, head and neck, and renal cell carcinomas (Park *et al*, 2013). In this study, we found Fostamatinib as a potent inhibitor of NEK9. The patient-derived TP-resistant cell-based xenograft data suggest that Fostamatinib restores Docetaxel sensitivity and facilitates a significant reduction of tumor burden. This unique combination of Docetaxel and Fostamatinib needs further clinical investigation in advanced carcinomas.

## 4. Material and methods

### 4.1 High content screening

Cas9 overexpressing OSCC line SCC4 (with three replicates) were seeded in a black, flat-bottomed 96-well plate (Thermo ScientificTM Nunc) and divided into two experimental groups, one vehicle control treatment and the other with Docetaxel treatment. A kinome-wide screening was performed using a lentiviral Sg RNA library (LentiArray™ Human Kinase CRISPR Library, Thermo Fisher Scientific, Cat No# M3775) that knocks out 840 kinase and kinase-related genes individually with a total number of 3214 Sg RNA constructs. Additionally, control lentiviruses, both positive and negative, were transduced. The transduction process was conducted in the presence of polybrene (8 µg/ml). Following a 48-hour transduction period, puromycin (0.5 µg/ml) selection was performed for the next 4 days to eliminate un-transduced cells. One experimental group was treated with a sub-lethal dose of Docetaxel (2nM) and another with vehicle control for 48 hours. Finally, cells were stained with LIVE/ DEAD™ Viability/Cytotoxicity Kit (Thermo Fisher Scientific, Cat No# L3224). Live and dead cell count and photographs in individual wells were taken with the help of Cell Insight CX7 High Content Screening (HCS) Platform (20 fields per well using 10X objective lens) based on intensity. The green fluorescence indicates the living cells and the red fluorescence indicates dead cells. For acquiring green and red fluorescence, two different fluorescent channels (excitation wavelengths −488 nm and 561 nm) were used. Image analysis was performed using the HCS Studio software. A threshold value for each channel was set once and used for the entire screening. To identify the cells, segmentation was performed. Some of the clumped and poorly segmented cells were excluded from further analysis on the basis of area, shape, and intensity. For positive control, cells were transduced with lentiviruses expressing Sg RNA targeting human hypoxanthine phosphoribosyltransferase 1 (HPRT1) (Lenti Array™ CRISPR Positive Control Lentivirus, human HPRT, Thermo Fisher Scientific, Cat No# A32829). HPRT1 knockout cells showed resistance to 6-thioguanine (6TG) induced cell death. For negative control, cells were transduced with lentiviruses expressing Sg RNA with no sequence homology to any region of the human genome (LentiArray™ CRISPR Negative Control Lentivirus, Thermo Fisher Scientific, Cat No# A32327). Vehicle-treated knockouts were normalized to vehicle-treated negative control Sg RNA whereas, Docetaxel-treated knockout cells were normalized to Docetaxel-treated negative control Sg RNA to get the appropriate % cell viability of each target knockout. The quantification scores of primary screening and secondary screening are mentioned in Tables 1, 2, and 3.

**Table 1.** Primary screening score from high content analyser system in SCC4 Cas9 cells: % Cell viability of rejected (red dot 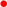) lethal knockout groups ( cell viability <65%) in absence and presence of docetaxel. This is in reference to figure 1B.

| Sl | Kinases | Cell viability of KO alone<br><= 65% |  |  | Average<br>% |  | Cell viability of KO + Doc<br>treatment |  |  | Average<br>% |
| --- | --- | --- | --- | --- | --- | --- | --- | --- | --- | --- |
| 1 | MAST2 | 70.04 | 69.91 | 54.86 | **64.94** |  | 14.70 | 21.77 | 29.10 | **21.86** |
| 2 | CAMK1D | 64.17 | 55.44 | 75.19 | **64.94** |  | 35.27 | 40.59 | 50.40 | **42.09** |
| 3 | RP2 | 60.86 | 55.43 | 78.29 | **64.86** |  | 25.91 | 30.51 | 40.75 | **32.39** |
| 4 | DGKB | 64.86 | 55.50 | 74.22 | **64.86** |  | 42.44 | 46.64 | 56.68 | **48.58** |
| 5 | TNNI3K | 61.67 | 54.55 | 78.29 | **64.84** |  | 25.40 | 28.20 | 37.05 | **30.22** |
| 6 | NEK5 | 69.88 | 68.76 | 55.72 | **64.79** |  | 21.95 | 29.42 | 43.14 | **31.50** |
| 7 | MNAT1 | 69.72 | 68.60 | 55.60 | **64.64** |  | 2.46 | 16.86 | 16.73 | **12.02** |
| 8 | GK5 | 69.29 | 68.17 | 55.26 | **64.24** |  | 27.63 | 33.80 | 25.22 | **28.88** |
| 9 | TK1 | 67.22 | 63.19 | 62.27 | **64.23** |  | 49.94 | 52.72 | 40.50 | **47.72** |
| 10 | UHMK1 | 67.11 | 63.08 | 62.25 | **64.15** |  | 43.35 | 39.99 | 38.36 | **40.57** |
| 11 | WNK4 | 69.11 | 65.08 | 58.03 | **64.07** |  | 25.20 | 24.74 | 15.62 | **21.86** |
| 12 | DCAKD | 54.90 | 50.86 | 86.22 | **63.99** |  | 68.14 | 66.89 | 57.66 | **64.23** |
| 13 | PTK2 | 54.82 | 49.70 | 87.03 | **63.85** |  | 75.94 | 72.80 | 62.39 | **70.38** |
| 14 | RIPK2 | 54.64 | 49.52 | 86.65 | **63.60** |  | 36.39 | 32.46 | 21.95 | **30.27** |
| 15 | TAOK3 | 54.54 | 49.43 | 86.36 | **63.44** |  | 37.03 | 32.31 | 21.70 | **30.35** |
| 16 | TEC | 68.43 | 63.31 | 58.59 | **63.44** |  | 63.07 | 57.54 | 46.84 | **55.82** |
| 17 | TBRG4 | 68.28 | 73.80 | 47.83 | **63.30** |  | 37.48 | 41.80 | 41.63 | **40.31** |
| 18 | UCKL1 | 68.09 | 75.18 | 46.12 | **63.13** |  | 15.05 | 20.13 | 21.43 | **18.87** |
| 19 | PAK1 | 65.90 | 72.78 | 50.24 | **62.97** |  | 31.53 | 33.28 | 37.08 | **33.96** |
| 20 | MTOR | 65.87 | 72.74 | 50.30 | **62.97** |  | 23.29 | 25.00 | 27.89 | **25.39** |
| 21 | IDNK | 67.69 | 73.58 | 47.01 | **62.76** |  | 52.15 | 54.79 | 57.75 | **54.90** |
| 22 | MAP3K9 | 53.75 | 59.63 | 74.36 | **62.58** |  | 12.76 | 15.36 | 17.40 | **15.17** |
| 23 | CDC7 | 53.73 | 57.29 | 76.48 | **62.50** |  | 55.34 | 60.22 | 63.68 | **59.75** |
| 24 | DDR2 | 53.80 | 57.68 | 76.02 | **62.50** |  | 48.06 | 52.57 | 54.79 | **51.81** |
| 25 | MINK1 | 62.40 | 60.90 | 64.10 | **62.46** |  | 33.52 | 35.3 | 44.21 | **37.67** |
| 26 | NOL9 | 53.79 | 48.67 | 84.80 | **62.42** |  | 17.64 | 20.86 | 20.93 | **19.81** |
| 27 | YES1 | 66.99 | 61.87 | 57.46 | **62.11** |  | 63.81 | 66.99 | 66.13 | **65.64** |
| 28 | ROS1 | 66.89 | 61.78 | 57.38 | **62.02** |  | 40.64 | 43.77 | 42.00 | **42.14** |
| 29 | STK3 | 66.84 | 60.54 | 58.53 | **61.97** |  | 30.95 | 33.18 | 29.64 | **31.26** |
| 30 | PRKCB | 64.79 | 71.53 | 49.43 | **61.92** |  | 59.82 | 63.23 | 54.99 | **59.35** |
| 31 | TP53RK | 64.63 | 71.82 | 48.93 | **61.79** |  | 45.51 | 48.23 | 39.05 | **44.26** |
| 32 | SKAP1 | 66.57 | 74.19 | 44.39 | **61.71** |  | 49.15 | 56.25 | 51.20 | **52.20** |
| 33 | PIK3R4 | 53.11 | 61.17 | 70.74 | **61.67** |  | 37.50 | 44.24 | 38.58 | **40.11** |
| 34 | HMGA2 | 53.07 | 47.91 | 83.74 | **61.57** |  | 45.38 | 57.14 | 56.24 | **52.92** |
| 35 | STK35 | 53.12 | 55.42 | 76.13 | **61.56** |  | 42.23 | 41.75 | 41.26 | **41.75** |
| 36 | TAF1L | 52.96 | 54.62 | 76.39 | **61.32** |  | 38.55 | 40.74 | 35.10 | **38.13** |
| 37 | DCLK1 | 66.14 | 67.18 | 50.64 | **61.32** |  | 33.29 | 29.68 | 35.85 | **32.94** |
| 38 | TTN | 65.72 | 66.13 | 50.94 | **60.93** |  | 33.27 | 42.29 | 23.39 | **32.99** |
| 39 | RPS6KA3 | 65.61 | 65.39 | 51.49 | **60.83** |  | 41.72 | 50.46 | 32.30 | **41.49** |
| 40 | SPHK2 | 63.50 | 62.65 | 55.93 | **60.69** |  | 38.13 | 29.34 | 46.45 | **37.97** |
| 41 | MPP2 | 63.31 | 67.34 | 50.95 | **60.53** |  | 18.50 | 4.60 | 15.59 | **12.89** |
| 42 | MAP2K2 | 65.21 | 70.33 | 45.83 | 60.46 |  | 39.64 | 24.43 | 33.10 | 32.39 |
| 43 | ULK4 | 52.09 | 54.82 | 74.25 | 60.39 |  | 50.43 | 37.38 | 47.20 | 45.00 |
| 44 | PRKAB1 | 52.05 | 60.56 | 68.25 | 60.29 |  | 60.29 | 41.23 | 44.03 | 48.52 |
| 45 | PKM | 51.82 | 60.76 | 67.37 | 59.98 |  | 31.08 | 24.50 | 38.76 | 31.45 |
| 46 | DLG1 | 51.81 | 61.19 | 66.72 | 59.91 |  | 45.93 | 38.91 | 51.71 | 45.52 |
| 47 | SYK | 64.55 | 74.37 | 40.61 | 59.84 |  | 20.13 | 12.33 | 24.36 | 18.94 |
| 48 | ITPKA | 64.28 | 74.55 | 39.96 | 59.59 |  | 23.53 | 15.29 | 25.87 | 21.56 |
| 49 | RYK | 63.96 | 62.84 | 51.10 | 59.30 |  | 33.91 | 34.81 | 53.53 | 40.75 |
| 50 | ERN2 | 62.02 | 60.90 | 54.91 | 59.28 |  | 24.95 | 32.48 | 36.91 | 31.45 |
| 51 | PRKD3 | 61.91 | 57.74 | 57.94 | 59.20 |  | 27.65 | 31.17 | 33.47 | 30.76 |
| 52 | SBK2 | 63.85 | 58.35 | 55.40 | 59.20 |  | 31.58 | 32.80 | 34.68 | 33.02 |
| 53 | CIT | 50.92 | 44.84 | 81.10 | 58.95 |  | 85.92 | 85.60 | 87.83 | 86.45 |
| 54 | XRCC6BP1 | 50.99 | 57.73 | 68.14 | 58.95 |  | 9.15 | 24.08 | 34.28 | 22.50 |
| 55 | RIPK3 | 50.91 | 58.09 | 67.42 | 58.81 |  | 55.94 | 56.59 | 52.56 | 55.03 |
| 56 | PDXK | 50.92 | 48.50 | 76.84 | 58.75 |  | 29.32 | 39.24 | 44.50 | 37.69 |
| 57 | TBK1 | 63.16 | 59.81 | 52.70 | 58.56 |  | 16.61 | 25.47 | 29.72 | 23.94 |
| 58 | AK7 | 63.09 | 67.36 | 45.02 | 58.49 |  | 29.45 | 37.37 | 40.72 | 35.85 |
| 59 | XYLB | 63.05 | 68.25 | 44.07 | 58.46 |  | 42.19 | 37.66 | 38.25 | 39.37 |
| 60 | MERTK | 61.02 | 67.14 | 46.85 | 58.33 |  | 31.67 | 24.91 | 25.49 | 27.36 |
| 61 | PIK3C2G | 60.86 | 67.91 | 45.86 | 58.21 |  | 34.53 | 25.55 | 26.12 | 28.73 |
| 62 | PI4K2A | 62.58 | 57.46 | 54.01 | 58.02 |  | 18.49 | 20.37 | 34.02 | 24.29 |
| 63 | DMPK | 50.17 | 45.06 | 78.59 | 57.94 |  | 59.40 | 49.73 | 54.07 | 54.40 |
| 64 | CALM1 | 50.11 | 43.81 | 79.52 | 57.81 |  | 80.54 | 68.38 | 72.45 | 73.79 |
| 65 | ZAK | 50.07 | 43.33 | 79.71 | 57.70 |  | 45.17 | 31.27 | 35.82 | 37.42 |
| 66 | PI4K2B | 50.10 | 42.91 | 79.98 | 57.67 |  | 20.97 | 5.32 | 10.35 | 12.22 |
| 67 | MAST3 | 62.07 | 54.45 | 56.12 | 57.55 |  | 41.13 | 23.73 | 29.24 | 31.37 |
| 68 | MAPKAPK3 | 61.82 | 53.75 | 56.37 | 57.31 |  | 24.76 | 5.62 | 11.60 | 13.99 |
| 69 | TK2 | 61.77 | 66.94 | 43.10 | 57.27 |  | 47.11 | 58.13 | 41.80 | 49.01 |
| 70 | FGFRL1 | 59.85 | 57.56 | 54.29 | 57.23 |  | 28.76 | 45.63 | 32.93 | 35.77 |
| 71 | SHPK | 59.82 | 58.16 | 53.72 | 57.23 |  | 30.19 | 48.10 | 34.21 | 37.50 |
| 72 | ITK | 61.72 | 60.68 | 49.26 | 57.22 |  | 41.76 | 53.23 | 53.08 | 49.36 |
| 73 | EPHA8 | 49.58 | 49.17 | 72.48 | 57.08 |  | 22.30 | 33.00 | 35.27 | 30.19 |
| 74 | CDKL1 | 49.49 | 49.71 | 71.57 | 56.92 |  | 14.86 | 24.79 | 29.49 | 23.05 |
| 75 | ERBB2 | 49.39 | 50.25 | 70.64 | 56.76 |  | 37.99 | 47.13 | 54.27 | 46.46 |
| 76 | CLK3 | 49.46 | 47.89 | 72.93 | 56.76 |  | 23.69 | 29.00 | 35.51 | 29.40 |
| 77 | PI4KA | 61.14 | 59.58 | 49.33 | 56.68 |  | 30.70 | 34.62 | 42.94 | 36.08 |
| 78 | TAOK2 | 60.97 | 59.42 | 49.19 | 56.53 |  | 37.94 | 40.47 | 50.60 | 43.00 |
| 79 | TTBK1 | 60.92 | 59.38 | 49.14 | 56.48 |  | 13.18 | 14.64 | 26.93 | 18.25 |
| 80 | PKMYT1 | 58.89 | 57.36 | 52.74 | 56.33 |  | 50.35 | 50.42 | 64.52 | 55.09 |
| 81 | TTBK2 | 58.74 | 60.44 | 49.46 | 56.21 |  | 40.94 | 41.88 | 62.93 | 48.58 |
| 82 | ADCK1 | 60.63 | 63.07 | 44.93 | 56.21 |  | 57.95 | 65.12 | 90.14 | 71.07 |
| 83 | CRIM1 | 48.58 | 51.77 | 67.10 | 55.82 |  | 23.17 | 31.84 | 56.08 | 37.03 |
| 84 | TEK | 48.57 | 52.50 | 66.14 | 55.74 |  | 34.58 | 36.45 | 51.62 | 40.88 |
| 85 | STK38L | 48.49 | 53.16 | 65.12 | 55.59 |  | 49.55 | 52.36 | 66.19 | 56.03 |
| 86 | AKAP8L | 48.39 | 53.80 | 64.08 | 55.42 |  | 46.41 | 50.15 | 62.64 | 53.07 |
| 87 | DAK | 59.78 | 65.93 | 40.56 | 55.42 |  | 36.88 | 38.59 | 55.66 | 43.71 |
| 88 | FGFR1OP | 59.48 | 66.37 | 39.58 | 55.14 |  | 52.09 | 60.07 | 65.15 | 59.10 |
| 89 | LATS1 | 59.48 | 67.11 | 38.84 | 55.14 |  | 71.94 | 80.11 | 85.34 | 79.13 |
| 90 | MARK2 | 57.49 | 65.86 | 41.63 | 55.00 |  | 36.95 | 45.30 | 50.69 | 44.31 |
| 91 | CD2 | 57.46 | 66.58 | 40.95 | 55.00 |  | 4.77 | 13.73 | 18.45 | 12.31 |
| 92 | PKN1 | 59.19 | 69.05 | 36.39 | 54.87 |  | 27.50 | 37.44 | 40.72 | 35.22 |
| 93 | RIOK2 | 47.83 | 58.43 | 58.13 | 54.80 |  | 22.21 | 33.14 | 34.98 | 30.11 |
| 94 | CSNK1G2 | 47.74 | 51.62 | 64.56 | 54.64 |  | 38.30 | 42.75 | 35.69 | 38.92 |
| 95 | STK16 | 47.73 | 50.53 | 65.42 | 54.56 |  | 36.55 | 53.31 | 56.14 | 48.66 |
| 96 | CPNE3 | 47.79 | 49.53 | 66.35 | 54.56 |  | 41.10 | 57.26 | 57.07 | 51.81 |
| 97 | MOK | 58.57 | 59.24 | 45.09 | 54.30 |  | 41.83 | 57.40 | 54.18 | 51.14 |
| 98 | SH3BP4 | 58.34 | 57.94 | 45.98 | 54.09 |  | 39.35 | 54.31 | 48.08 | 47.25 |
| 99 | PRPF4B | 58.31 | 56.84 | 47.03 | 54.06 |  | 25.71 | 40.07 | 30.81 | 32.20 |
| 100 | SMG1 | 56.44 | 53.90 | 51.68 | 54.01 |  | 40.51 | 31.30 | 64.98 | 45.60 |
| 101 | PSKH1 | 56.31 | 52.70 | 52.71 | 53.91 |  | 36.03 | 46.58 | 36.52 | 39.71 |
| 102 | ITPKC | 58.09 | 53.41 | 50.06 | 53.85 |  | 19.82 | 20.74 | 25.71 | 22.09 |
| 103 | TRIM27 | 47.15 | 41.41 | 73.00 | 53.85 |  | 49.53 | 59.63 | 42.02 | 50.39 |
| 104 | ULK2 | 47.13 | 40.31 | 73.84 | 53.76 |  | 33.44 | 38.58 | 27.69 | 33.23 |
| 105 | TEX14 | 46.97 | 39.09 | 74.55 | 53.54 |  | 38.95 | 45.64 | 27.44 | 37.34 |
| 106 | RPS6KA6 | 46.80 | 37.85 | 75.25 | 53.30 |  | 29.90 | 34.85 | 15.91 | 26.89 |
| 107 | MORN1 | 57.49 | 67.51 | 34.90 | 53.30 |  | 11.53 | 19.81 | 5.21 | 12.19 |
| 108 | ANKK1 | 57.49 | 68.58 | 33.83 | 53.30 |  | 73.69 | 80.56 | 65.57 | 73.27 |
| 109 | WEE2 | 57.49 | 69.65 | 32.76 | 53.30 |  | 76.61 | 73.77 | 50.09 | 66.82 |
| 110 | PRKD2 | 55.59 | 68.82 | 35.25 | 53.22 |  | 31.70 | 26.89 | 2.25 | 20.28 |
| 111 | NMRK1 | 55.55 | 69.85 | 34.24 | 53.21 |  | 36.99 | 17.33 | 17.50 | 23.94 |
| 112 | PIP4K2A | 57.34 | 50.47 | 51.68 | 53.17 |  | 21.33 | 22.61 | 42.71 | 28.88 |
| 113 | CLK4 | 46.63 | 47.98 | 64.58 | 53.07 |  | 29.37 | 22.21 | 32.85 | 28.14 |
| 114 | PRPSAP1 | 46.65 | 37.41 | 74.99 | 53.02 |  | 26.57 | 24.69 | 54.83 | 35.36 |
| 115 | PLK5 | 46.72 | 41.37 | 70.96 | 53.02 |  | 23.90 | 17.57 | 43.25 | 28.24 |
| 116 | MST1R | 46.75 | 45.30 | 66.91 | 52.99 |  | 18.04 | 15.55 | 20.18 | 17.92 |
| 117 | PRKG1 | 57.15 | 49.59 | 52.22 | 52.99 |  | 12.26 | 15.88 | 24.93 | 17.69 |
| 118 | LRGUK | 56.98 | 69.76 | 31.75 | 52.83 |  | 50.22 | 33.50 | 20.52 | 34.75 |
| 119 | RELN | 56.98 | 69.00 | 32.51 | 52.83 |  | 64.81 | 48.85 | 34.93 | 49.53 |
| 120 | FLT3 | 55.06 | 66.32 | 36.88 | 52.75 |  | 43.62 | 41.55 | 39.83 | 41.67 |
| 121 | MAP4K4 | 54.90 | 65.40 | 37.57 | 52.62 |  | 48.17 | 47.08 | 44.66 | 46.64 |
| 122 | CDKL4 | 56.76 | 66.50 | 34.60 | 52.62 |  | 47.65 | 47.55 | 44.42 | 46.54 |
| 123 | PDGFRA | 46.41 | 55.39 | 55.92 | 52.57 |  | 44.90 | 45.79 | 41.95 | 44.21 |
| 124 | CAB39 | 46.47 | 54.70 | 56.55 | 52.57 |  | 15.60 | 17.47 | 12.93 | 15.33 |
| 125 | GSG2 | 46.40 | 53.87 | 57.04 | 52.44 |  | 33.96 | 36.82 | 31.57 | 34.12 |
| 126 | ADCK3 | 46.23 | 52.94 | 57.43 | 52.20 |  | 33.46 | 37.30 | 31.35 | 34.04 |
| 127 | DAPK2 | 56.14 | 62.09 | 37.91 | 52.04 |  | 35.72 | 27.41 | 47.02 | 36.71 |
| 128 | MPP4 | 55.97 | 61.16 | 38.54 | 51.89 |  | 8.49 | 0.93 | 20.30 | 9.91 |
| 129 | ADPGK | 55.88 | 60.32 | 39.23 | 51.81 |  | 57.57 | 51.11 | 70.56 | 59.75 |
| 130 | CLK2 | 54.05 | 57.73 | 43.64 | 51.81 |  | 33.80 | 28.10 | 47.29 | 36.40 |
| 131 | STK32C | 53.86 | 56.78 | 44.31 | 51.65 |  | 37.62 | 32.68 | 51.62 | 40.64 |
| 132 | RPS6KC1 | 55.69 | 60.81 | 38.40 | 51.63 |  | 39.55 | 37.48 | 58.28 | 45.10 |
| 133 | ABL1 | 45.59 | 44.63 | 64.26 | 51.49 |  | 27.43 | 31.79 | 58.00 | 39.07 |
| 134 | SRMS | 45.58 | 44.60 | 64.06 | 51.42 |  | 9.70 | 6.10 | 23.35 | 13.05 |
| 135 | BMPRI1B | 45.65 | 44.66 | 63.94 | 51.42 |  | 38.08 | 34.27 | 50.29 | 40.88 |
| 136 | PTK2B | 45.64 | 44.22 | 64.15 | 51.34 |  | 41.05 | 37.10 | 52.98 | 43.71 |
| 137 | DNAJC6 | 55.10 | 52.88 | 45.28 | 51.09 |  | 49.12 | 45.75 | 61.21 | 52.03 |
| 138 | DGKE | 55.03 | 52.01 | 46.03 | 51.02 |  | 23.11 | 20.32 | 35.34 | 26.26 |
| 139 | TGFBR3 | 55.00 | 51.16 | 46.81 | 50.99 |  | 33.85 | 26.57 | 51.31 | 37.24 |
| 140 | TGFB2 | 53.17 | 48.53 | 51.27 | 50.99 |  | 38.85 | 31.81 | 56.79 | 42.48 |
| 141 | NME2 | 52.78 | 47.33 | 51.78 | 50.63 |  | 10.11 | 11.62 | 20.25 | 13.99 |
| 142 | CDK7 | 54.52 | 48.27 | 48.86 | 50.55 |  | 39.35 | 41.66 | 49.41 | 43.47 |
| 143 | WNK1 | 44.92 | 37.85 | 68.88 | 50.55 |  | 69.12 | 58.11 | 50.84 | 59.36 |
| 144 | STK36 | 44.98 | 37.10 | 69.55 | 50.54 |  | 55.05 | 43.23 | 33.46 | 43.92 |
| 145 | SIK2 | 45.04 | 59.34 | 47.24 | 50.54 |  | 5.46 | 29.41 | 19.58 | 18.15 |
| 146 | TTK | 45.06 | 38.19 | 68.23 | 50.49 |  | 16.45 | 29.05 | 10.14 | 18.55 |
| 147 | NADK2 | 54.35 | 53.01 | 43.82 | 50.39 |  | 17.69 | 37.42 | 27.91 | 27.67 |
| 148 | HIPK3 | 54.27 | 63.51 | 33.17 | 50.31 |  | 42.06 | 50.10 | 31.19 | 41.12 |
| 149 | RIOK1 | 54.25 | 59.60 | 37.04 | 50.30 |  | 18.00 | 39.53 | 36.38 | 31.31 |
| 150 | PIIP5K1 | 52.12 | 53.57 | 44.31 | 50.00 |  | 17.65 | 34.20 | 28.34 | 26.73 |
| 151 | PDIK1L | 52.09 | 59.65 | 38.26 | 50.00 |  | 10.47 | 32.03 | 33.46 | 25.32 |
| 152 | RPS6KB2 | 53.68 | 40.90 | 54.72 | 49.76 |  | 39.63 | 39.76 | 22.03 | 33.81 |
| 153 | EPHB6 | 44.32 | 32.30 | 72.44 | 49.69 |  | 47.31 | 47.10 | 31.30 | 41.90 |
| 154 | DAPK1 | 44.07 | 32.81 | 71.23 | 49.37 |  | 44.98 | 44.76 | 31.25 | 40.33 |
| 155 | TRAF3IP3 | 44.06 | 33.56 | 70.26 | 49.29 |  | 65.94 | 68.94 | 50.27 | 61.71 |
| 156 | RPS6KA2 | 44.13 | 34.38 | 69.37 | 49.29 |  | 40.68 | 43.80 | 26.13 | 36.87 |
| 157 | PRKACG | 52.83 | 43.84 | 50.26 | 48.98 |  | 17.41 | 25.75 | 14.15 | 19.10 |
| 158 | NEK4 | 52.76 | 44.53 | 49.45 | 48.91 |  | 25.00 | 31.61 | 25.73 | 27.45 |
| 159 | CDKL2 | 52.70 | 45.24 | 48.65 | 48.86 |  | 35.85 | 26.52 | 29.91 | 30.76 |
| 160 | KCNH3 | 50.52 | 43.81 | 51.06 | 48.47 |  | 56.04 | 47.58 | 49.79 | 51.14 |
| 161 | IGSF22 | 50.44 | 44.49 | 50.32 | 48.42 |  | 33.83 | 25.50 | 28.06 | 29.13 |
| 162 | RPS6KA1 | 51.90 | 46.70 | 45.74 | 48.11 |  | 39.26 | 31.04 | 33.94 | 34.75 |
| 163 | PRKCA | 42.86 | 38.42 | 62.59 | 47.96 |  | 19.65 | 11.55 | 14.80 | 15.33 |
| 164 | PTK6 | 42.92 | 35.70 | 65.24 | 47.96 |  | 27.24 | 20.82 | 10.66 | 19.58 |
| 165 | GUCY2F | 42.99 | 36.06 | 64.82 | 47.96 |  | 44.50 | 37.24 | 27.93 | 36.56 |
| 166 | TRIM28 | 42.82 | 36.18 | 64.17 | 47.73 |  | 22.16 | 14.07 | 5.61 | 13.95 |
| 167 | EPHA10 | 51.47 | 47.31 | 44.38 | 47.72 |  | 31.61 | 24.87 | 19.46 | 25.31 |
| 168 | PRKG2 | 51.47 | 47.23 | 44.45 | 47.72 |  | 47.50 | 39.56 | 34.64 | 40.57 |
| 169 | MADD | 51.42 | 47.87 | 43.73 | 47.68 |  | 20.91 | 12.52 | 8.85 | 14.09 |
| 170 | PIP5K1B | 49.67 | 46.04 | 47.32 | 47.68 |  | 20.93 | 24.47 | 34.42 | 26.61 |
| 171 | ETNK1 | 49.59 | 45.88 | 47.41 | 47.63 |  | 42.46 | 45.03 | 55.69 | 47.73 |
| 172 | RIPK1 | 51.30 | 47.52 | 43.87 | 47.56 |  | 34.22 | 35.81 | 47.19 | 39.07 |
| 173 | RPS6KL1 | 42.73 | 38.87 | 61.09 | 47.56 |  | 37.94 | 38.55 | 50.64 | 42.37 |
| 174 | SPHK1 | 42.80 | 38.10 | 61.79 | 47.56 |  | 42.49 | 55.06 | 39.71 | 45.75 |
| 175 | CCL3 | 42.83 | 38.06 | 61.70 | 47.53 |  | 35.48 | 47.30 | 32.21 | 38.33 |
| 176 | PRKD1 | 42.69 | 37.47 | 61.82 | 47.33 |  | 23.81 | 34.50 | 19.29 | 25.86 |
| 177 | MAGI1 | 50.96 | 45.36 | 45.43 | 47.25 |  | 34.51 | 44.15 | 28.89 | 35.85 |
| 178 | TNK2 | 50.89 | 46.50 | 44.15 | 47.18 |  | 29.08 | 39.27 | 25.57 | 31.31 |
| 179 | TESK1 | 50.71 | 46.24 | 44.09 | 47.01 |  | 27.41 | 21.94 | 39.33 | 29.56 |
| 180 | MAGI3 | 48.72 | 44.17 | 47.46 | 46.79 |  | 50.39 | 44.84 | 61.14 | 52.13 |
| 181 | PLXNB1 | 48.68 | 44.06 | 47.59 | 46.78 |  | 62.52 | 56.89 | 72.10 | 63.84 |
| 182 | DGKG | 50.20 | 48.01 | 41.41 | 46.54 |  | 40.92 | 42.80 | 64.50 | 49.41 |
| 183 | ARAF | 41.89 | 39.94 | 57.56 | 46.46 |  | 43.64 | 45.75 | 67.68 | 52.36 |
| 184 | SGK494 | 41.89 | 40.16 | 57.12 | 46.39 |  | 27.25 | 21.62 | 34.82 | 27.89 |
| 185 | AK5 | 41.87 | 40.38 | 56.67 | 46.31 |  | 60.30 | 54.68 | 66.87 | 60.61 |
| 186 | CSNK1D | 41.86 | 40.60 | 56.22 | 46.23 |  | 48.84 | 38.15 | 59.48 | 48.82 |
| 187 | MAP3K1 | 49.72 | 48.69 | 39.88 | 46.09 |  | 24.18 | 13.16 | 34.17 | 23.84 |
| 188 | RIPK4 | 49.61 | 48.81 | 39.56 | 45.99 |  | 46.43 | 43.38 | 47.45 | 45.75 |
| 189 | LCK | 49.52 | 48.96 | 39.26 | 45.91 |  | 24.67 | 21.85 | 24.46 | 23.66 |
| 190 | MYLK2 | 47.76 | 47.43 | 42.49 | 45.90 |  | 20.18 | 17.60 | 18.75 | 18.84 |
| 191 | ASB10 | 47.68 | 47.58 | 42.27 | 45.85 |  | 25.86 | 23.50 | 23.19 | 24.18 |
| 192 | MAP3K7CL | 49.18 | 51.45 | 36.16 | 45.60 |  | 19.52 | 30.91 | 8.53 | 19.65 |
| 193 | FGFR4 | 41.14 | 36.02 | 59.16 | 45.44 |  | 38.40 | 41.08 | 12.26 | 30.58 |
| 194 | FLT4 | 41.05 | 46.57 | 48.23 | 45.28 |  | 33.91 | 45.91 | 28.68 | 36.16 |
| 195 | JAK3 | 41.12 | 42.12 | 52.61 | 45.28 |  | 31.59 | 50.89 | 43.23 | 41.90 |
| 196 | PRPS1L1 | 41.18 | 41.83 | 52.84 | 45.28 |  | 13.73 | 31.58 | 24.74 | 23.35 |
| 197 | SRPK3 | 48.76 | 50.39 | 36.46 | 45.20 |  | 48.27 | 62.73 | 54.80 | 55.27 |
| 198 | PLXNA4 | 48.65 | 51.04 | 35.62 | 45.10 |  | 13.23 | 25.85 | 18.34 | 19.14 |
| 199 | STK11 | 48.50 | 51.65 | 34.75 | 44.97 |  | 29.55 | 40.31 | 33.21 | 34.36 |
| 200 | MFHAS1 | 46.62 | 50.53 | 37.29 | 44.81 |  | 12.79 | 21.70 | 15.03 | 16.51 |
| 201 | EEF2K | 46.51 | 51.17 | 36.52 | 44.73 |  | 33.31 | 40.36 | 34.11 | 35.93 |
| 202 | PRPSAP2 | 48.25 | 53.67 | 32.27 | 44.73 |  | 12.55 | 21.29 | 8.38 | 14.07 |
| 203 | MAP3K19 | 40.62 | 46.81 | 46.54 | 44.66 |  | 4.37 | 24.09 | 23.02 | 17.16 |
| 204 | VRK1 | 40.54 | 47.48 | 45.50 | 44.51 |  | 13.99 | 27.43 | 41.22 | 27.55 |
| 205 | UCK2 | 40.52 | 41.07 | 51.67 | 44.42 |  | 0.30 | 6.97 | 12.30 | 6.53 |
| 206 | ERBB3 | 40.35 | 41.36 | 50.84 | 44.18 |  | 28.19 | 36.15 | 41.08 | 35.14 |
| 207 | MRC2 | 47.40 | 48.36 | 36.08 | 43.95 |  | 2.13 | 15.33 | 23.82 | 13.76 |
| 208 | CSNK2A2 | 47.32 | 48.35 | 35.94 | 43.87 |  | 49.41 | 61.19 | 66.56 | 59.05 |
| 209 | FASTKD1 | 47.23 | 48.03 | 36.11 | 43.79 |  | 24.68 | 28.96 | 25.14 | 26.26 |
| 210 | BAZ1B | 45.36 | 45.93 | 39.57 | 43.62 |  | 36.39 | 43.38 | 33.14 | 37.64 |
| 211 | IP6K1 | 45.27 | 45.60 | 39.80 | 43.55 |  | 44.45 | 52.18 | 39.46 | 45.36 |
| 212 | INSR | 46.89 | 46.99 | 36.54 | 43.47 |  | 49.37 | 57.83 | 42.65 | 49.95 |
| 213 | IP6K3 | 39.63 | 37.36 | 53.20 | 43.40 |  | 35.58 | 46.93 | 31.41 | 37.97 |
| 214 | CKMT2 | 39.46 | 37.21 | 52.81 | 43.16 |  | 25.55 | 37.38 | 19.14 | 27.36 |
| 215 | HK3 | 39.53 | 37.30 | 52.66 | 43.16 |  | 50.11 | 62.43 | 41.47 | 51.34 |
| 216 | STK31 | 39.56 | 38.87 | 50.95 | 43.13 |  | 28.73 | 40.00 | 14.80 | 27.84 |
| 217 | PIP5K1A | 46.36 | 39.60 | 42.98 | 42.98 |  | 18.92 | 36.77 | 14.93 | 23.54 |
| 218 | CHKA | 46.30 | 45.44 | 37.03 | 42.92 |  | 47.83 | 60.29 | 29.85 | 45.99 |
| 219 | TRIM24 | 46.30 | 45.50 | 36.97 | 42.92 |  | 7.58 | 1.76 | 5.99 | 5.11 |
| 220 | FGFR3 | 44.53 | 43.85 | 40.16 | 42.85 |  | 40.57 | 51.90 | 59.19 | 50.55 |
| 221 | LAMTOR3 | 44.48 | 43.59 | 40.42 | 42.83 |  | 46.42 | 47.05 | 75.37 | 56.28 |
| 222 | DYRK1A | 46.04 | 44.95 | 37.07 | 42.69 |  | 9.85 | 14.23 | 34.65 | 19.58 |
| 223 | PRKCE | 39.11 | 37.80 | 50.92 | 42.61 |  | 11.70 | 19.85 | 32.36 | 21.31 |
| 224 | ERCC3 | 39.10 | 40.61 | 47.88 | 42.53 |  | 29.75 | 31.67 | 56.27 | 39.23 |
| 225 | IGF2R | 39.16 | 40.89 | 47.54 | 42.53 |  | 19.82 | 41.94 | 25.74 | 29.17 |
| 226 | BMPRI1A | 39.08 | 41.02 | 47.05 | 42.38 |  | 39.05 | 60.29 | 45.47 | 48.27 |
| 227 | PSTK | 45.62 | 47.77 | 33.50 | 42.30 |  | 17.56 | 33.29 | 29.10 | 26.65 |
| 228 | PRPS2 | 45.50 | 47.86 | 33.20 | 42.19 |  | 19.03 | 32.15 | 32.80 | 27.99 |
| 229 | DAPK3 | 45.37 | 47.93 | 32.88 | 42.06 |  | 22.51 | 27.44 | 45.10 | 31.68 |
| 230 | ACVR2A | 43.52 | 46.30 | 35.89 | 41.90 |  | 34.11 | 39.61 | 55.76 | 43.16 |
| 231 | EIF2AK1 | 43.33 | 46.31 | 35.58 | 41.74 |  | 29.50 | 34.80 | 50.98 | 38.43 |
| 232 | ICK | 45.02 | 48.22 | 31.98 | 41.74 |  | 15.72 | 19.49 | 34.38 | 23.19 |
| 233 | TGFBR2 | 38.51 | 41.91 | 44.81 | 41.74 |  | 43.08 | 46.56 | 61.40 | 50.35 |
| 234 | GSK3A | 38.34 | 39.61 | 46.58 | 41.51 |  | 33.96 | 31.21 | 40.03 | 35.06 |
| 235 | LTK | 38.25 | 39.59 | 46.22 | 41.35 |  | 33.80 | 29.58 | 39.44 | 34.28 |
| 236 | AK3 | 38.31 | 39.71 | 46.01 | 41.35 |  | 46.70 | 41.77 | 51.15 | 46.54 |
| 237 | KSR1 | 44.52 | 45.99 | 33.31 | 41.27 |  | 16.28 | 10.26 | 19.92 | 15.49 |
| 238 | MASTL | 44.44 | 42.90 | 36.26 | 41.20 |  | 43.00 | 39.05 | 51.92 | 44.66 |
| 239 | GRK7 | 44.35 | 42.74 | 36.26 | 41.12 |  | 39.32 | 36.10 | 46.04 | 40.49 |
| 240 | MPP3 | 42.59 | 40.92 | 39.60 | 41.04 |  | 1.85 | 1.95 | 3.75 | 2.52 |
| 241 | BLK | 42.48 | 40.74 | 39.65 | 40.96 |  | 57.14 | 52.77 | 62.02 | 57.31 |
| 242 | TNK1 | 44.17 | 42.36 | 36.32 | 40.95 |  | 37.37 | 32.42 | 41.34 | 37.04 |
| 243 | CKM | 37.91 | 34.70 | 50.03 | 40.88 |  | 46.31 | 45.14 | 50.54 | 47.33 |
| 244 | SKP2 | 37.95 | 36.46 | 48.15 | 40.85 |  | 19.92 | 16.19 | 18.34 | 18.15 |
| 245 | NTPCR | 37.89 | 36.01 | 48.27 | 40.72 |  | 6.55 | 2.91 | 4.46 | 4.64 |
| 246 | PRKX | 37.96 | 36.01 | 48.21 | 40.72 |  | 23.99 | 33.41 | 47.32 | 34.91 |
| 247 | PTCD2 | 43.92 | 41.91 | 36.33 | 40.72 |  | 9.24 | 18.63 | 31.80 | 19.89 |
| 248 | PRKAA1 | 43.42 | 41.33 | 36.00 | 40.25 |  | 5.19 | 14.54 | 26.97 | 15.57 |
| 249 | FASTKD3 | 43.33 | 48.18 | 29.01 | 40.17 |  | 30.14 | 32.46 | 37.17 | 33.25 |
| 250 | MAPK8 | 41.60 | 46.52 | 32.20 | 40.11 |  | 13.27 | 15.41 | 19.24 | 15.97 |
| 251 | FGR | 41.48 | 46.48 | 32.09 | 40.02 |  | 39.93 | 41.88 | 44.84 | 42.22 |
| 252 | SPEF2 | 43.16 | 42.58 | 34.30 | 40.02 |  | 23.45 | 30.87 | 38.60 | 30.97 |
| 253 | ACVR1C | 37.24 | 40.12 | 42.45 | 39.94 |  | 55.16 | 61.15 | 60.34 | 58.88 |
| 254 | MAP2K7 | 37.22 | 39.88 | 42.47 | 39.86 |  | 16.88 | 25.95 | 19.67 | 20.83 |
| 255 | TPR | 37.21 | 39.64 | 42.49 | 39.78 |  | 20.44 | 18.77 | 33.66 | 24.29 |
| 256 | CALM2 | 37.16 | 39.35 | 42.48 | 39.66 |  | 34.09 | 36.83 | 42.59 | 37.83 |
| 257 | MAPK1 | 42.65 | 44.61 | 31.37 | 39.54 |  | 12.67 | 20.74 | 30.75 | 21.38 |
| 258 | IGF1R | 42.57 | 44.30 | 31.53 | 39.47 |  | 27.11 | 34.42 | 46.63 | 36.05 |
| 259 | AXL | 42.46 | 40.97 | 34.67 | 39.37 |  | 32.11 | 30.60 | 36.26 | 32.99 |
| 260 | GSK3B | 40.74 | 39.48 | 37.71 | 39.31 |  | 44.89 | 42.29 | 49.14 | 45.44 |
| 261 | ACVR1B | 40.63 | 39.94 | 37.12 | 39.23 |  | 45.51 | 42.16 | 50.54 | 46.07 |
| 262 | HKDC1 | 42.23 | 41.83 | 33.39 | 39.15 |  | 20.47 | 16.09 | 25.71 | 20.75 |
| 263 | NME6 | 36.64 | 36.53 | 44.05 | 39.07 |  | 21.13 | 15.72 | 26.59 | 21.15 |
| 264 | ROR1 | 36.65 | 36.84 | 43.57 | 39.02 |  | 43.88 | 37.44 | 49.54 | 43.62 |
| 265 | RFK | 36.52 | 36.99 | 42.95 | 38.82 |  | 19.48 | 12.00 | 25.35 | 18.94 |
| 266 | EPHA4 | 36.44 | 37.20 | 42.38 | 38.67 |  | 58.43 | 57.06 | 57.36 | 57.62 |
| 267 | SOX9 | 41.56 | 42.61 | 31.42 | 38.53 |  | 56.23 | 54.13 | 55.07 | 55.14 |
| 268 | DLG2 | 41.04 | 42.38 | 30.73 | 38.05 |  | 23.53 | 21.21 | 21.77 | 22.17 |
| 269 | MAP3K13 | 40.97 | 42.60 | 30.38 | 37.98 |  | 19.98 | 19.38 | 15.69 | 18.35 |
| 270 | GRIP2 | 38.79 | 40.71 | 32.76 | 37.42 |  | 18.24 | 26.43 | 32.22 | 25.63 |
| 271 | PRKCH | 38.68 | 42.56 | 30.79 | 37.34 |  | 35.24 | 40.90 | 44.85 | 40.33 |
| 272 | JAK1 | 40.11 | 44.08 | 27.37 | 37.19 |  | 43.29 | 48.29 | 52.29 | 47.96 |
| 273 | DGKA | 34.87 | 38.93 | 37.33 | 37.04 |  | 21.13 | 25.47 | 29.52 | 25.37 |
| 274 | IKBKG | 34.93 | 39.06 | 37.10 | 37.03 |  | 18.42 | 19.98 | 28.34 | 22.25 |
| 275 | PTK7 | 34.99 | 39.21 | 36.88 | 37.03 |  | 18.37 | 19.53 | 27.43 | 21.78 |
| 276 | FASTKD5 | 34.82 | 34.70 | 40.86 | 36.79 |  | 14.70 | 19.62 | 31.24 | 21.86 |
| 277 | TPK1 | 39.63 | 39.51 | 31.10 | 36.75 |  | 41.55 | 47.68 | 55.88 | 48.37 |
| 278 | CSNK2A1 | 39.60 | 39.47 | 31.07 | 36.71 |  | 22.68 | 22.41 | 42.41 | 29.17 |
| 279 | TGFBR1 | 39.26 | 39.13 | 30.80 | 36.40 |  | 29.42 | 35.05 | 43.24 | 35.91 |
| 280 | MAPK12 | 37.63 | 36.51 | 34.83 | 36.32 |  | 15.52 | 21.47 | 25.03 | 20.68 |
| 281 | STRADA | 37.57 | 36.45 | 34.87 | 36.30 |  | 12.54 | 18.04 | 21.50 | 17.36 |
| 282 | VRK2 | 39.01 | 37.89 | 31.60 | 36.16 |  | 28.89 | 39.33 | 47.11 | 38.44 |
| 283 | HK2 | 34.20 | 33.08 | 41.03 | 36.10 |  | 16.21 | 26.86 | 34.22 | 25.77 |
| 284 | COASY | 34.17 | 30.14 | 43.71 | 36.01 |  | 23.59 | 23.24 | 18.97 | 21.93 |
| 285 | LTBP4 | 34.16 | 30.13 | 43.50 | 35.93 |  | 31.70 | 31.00 | 25.75 | 29.48 |
| 286 | CHUK | 34.23 | 30.19 | 43.36 | 35.93 |  | 38.32 | 37.26 | 31.03 | 35.53 |
| 287 | PPP4C | 38.67 | 42.70 | 26.18 | 35.85 |  | 28.50 | 27.09 | 19.87 | 25.16 |
| 288 | CHEK2 | 38.67 | 43.78 | 25.10 | 35.85 |  | 33.93 | 31.08 | 21.79 | 28.93 |
| 289 | ADRBK1 | 38.57 | 43.68 | 25.02 | 35.76 |  | 43.15 | 39.94 | 29.67 | 37.59 |
| 290 | MAPK9 | 36.85 | 41.97 | 28.01 | 35.61 |  | 30.41 | 26.85 | 15.59 | 24.28 |
| 291 | MAP3K11 | 36.75 | 41.86 | 27.99 | 35.53 |  | 34.36 | 30.45 | 18.21 | 27.67 |
| 292 | STK17A | 38.30 | 32.78 | 35.45 | 35.51 |  | 26.29 | 32.66 | 30.07 | 29.67 |
| 293 | IRAK2 | 33.82 | 26.74 | 45.81 | 35.46 |  | 23.81 | 31.38 | 29.38 | 28.19 |
| 294 | MAPK10 | 33.81 | 26.94 | 45.39 | 35.38 |  | 6.53 | 13.55 | 10.34 | 10.14 |
| 295 | CAMK2A | 33.66 | 26.79 | 45.04 | 35.16 |  | 13.20 | 30.65 | 44.29 | 29.38 |
| 296 | CSF1R | 33.71 | 27.82 | 43.90 | 35.14 |  | 11.25 | 27.44 | 39.62 | 26.10 |
| 297 | EFNA3 | 37.57 | 31.68 | 35.23 | 34.83 |  | 16.93 | 32.85 | 44.56 | 31.45 |
| 298 | TLK1 | 37.45 | 33.89 | 32.81 | 34.72 |  | 27.96 | 41.28 | 50.19 | 39.81 |
| 299 | EPHA3 | 37.39 | 33.52 | 33.09 | 34.67 |  | 20.95 | 34.33 | 43.08 | 32.79 |
| 300 | MAP3K15 | 35.61 | 40.73 | 26.96 | 34.43 |  | 30.28 | 32.68 | 35.39 | 32.78 |
| 301 | NME1 | 35.47 | 40.59 | 26.91 | 34.32 |  | 9.80 | 12.15 | 13.95 | 11.97 |
| 302 | MAP3K12 | 36.72 | 41.83 | 23.57 | 34.04 |  | 35.12 | 37.42 | 38.31 | 36.95 |
| 303 | ROCK1 | 32.06 | 38.37 | 29.87 | 33.43 |  | 17.00 | 18.07 | 16.85 | 17.31 |
| 304 | TPD52L3 | 32.11 | 38.85 | 29.27 | 33.41 |  | 33.86 | 34.78 | 32.54 | 33.73 |
| 305 | LY6G5B | 32.10 | 39.28 | 28.62 | 33.33 |  | 14.56 | 14.99 | 11.40 | 13.65 |
| 306 | FGFR2 | 32.09 | 39.71 | 27.97 | 33.25 |  | 33.92 | 33.86 | 28.91 | 32.23 |
| 307 | LRRK1 | 35.87 | 27.80 | 36.09 | 33.25 |  | 20.82 | 25.35 | 24.12 | 23.43 |
| 308 | PLXND1 | 35.87 | 41.04 | 22.86 | 33.25 |  | 1.30 | 18.43 | 23.91 | 14.54 |
| 309 | PKN2 | 35.79 | 33.50 | 30.26 | 33.18 |  | 25.49 | 29.54 | 21.68 | 25.57 |
| 310 | ACVR1 | 34.29 | 32.63 | 32.61 | 33.18 |  | 38.30 | 42.49 | 34.54 | 38.44 |
| 311 | AKAP12 | 34.02 | 32.99 | 31.80 | 32.94 |  | 35.30 | 31.37 | 39.86 | 35.51 |
| 312 | CSNK1G3 | 35.45 | 35.04 | 28.10 | 32.86 |  | 14.82 | 10.88 | 19.59 | 15.09 |
| 313 | KNDC1 | 31.44 | 31.67 | 34.53 | 32.55 |  | 14.52 | 15.19 | 14.87 | 14.86 |
| 314 | NEK2 | 31.51 | 32.36 | 33.76 | 32.54 |  | 30.80 | 31.31 | 31.51 | 31.21 |
| 315 | CDKL3 | 31.50 | 27.47 | 38.44 | 32.47 |  | 50.16 | 42.86 | 42.36 | 45.13 |
| 316 | KCNH1 | 31.39 | 26.28 | 39.22 | 32.29 |  | 27.16 | 25.59 | 8.67 | 20.47 |
| 317 | EPHB4 | 34.68 | 31.95 | 29.83 | 32.15 |  | 33.56 | 23.17 | 29.35 | 28.69 |
| 318 | CIB4 | 34.68 | 27.50 | 34.28 | 32.15 |  | 42.51 | 31.62 | 23.75 | 32.63 |
| 319 | SH3BP5L | 33.98 | 26.35 | 34.18 | 31.50 |  | 28.10 | 16.56 | 6.97 | 17.21 |
| 320 | ENG | 32.50 | 24.44 | 37.40 | 31.45 |  | 24.07 | 23.68 | 28.19 | 25.31 |
| 321 | VRK3 | 32.43 | 37.59 | 24.19 | 31.40 |  | 10.49 | 22.24 | 41.16 | 24.63 |
| 322 | MAPK7 | 33.83 | 31.54 | 28.73 | 31.37 |  | 2.72 | 0.47 | 7.66 | 3.62 |
| 323 | PLXNC1 | 30.53 | 28.87 | 34.70 | 31.37 |  | 12.24 | 14.96 | 29.41 | 18.87 |
| 324 | EPGN | 30.52 | 31.56 | 31.79 | 31.29 |  | 25.62 | 27.88 | 44.14 | 32.55 |
| 325 | LTBP1 | 30.55 | 30.96 | 32.26 | 31.26 |  | 4.85 | 6.64 | 24.71 | 12.07 |
| 326 | NEK10 | 30.32 | 30.10 | 32.45 | 30.96 |  | 13.96 | 15.28 | 35.16 | 21.46 |
| 327 | KALRN | 33.33 | 32.47 | 26.89 | 30.90 |  | 22.23 | 23.09 | 44.77 | 30.03 |
| 328 | PASK | 33.29 | 34.86 | 24.43 | 30.86 |  | 19.99 | 24.47 | 30.62 | 25.02 |
| 329 | CDK14 | 32.99 | 34.55 | 24.21 | 30.58 |  | 20.26 | 24.30 | 31.38 | 25.31 |
| 330 | PRKAG1 | 31.57 | 33.12 | 27.05 | 30.58 |  | 14.61 | 18.38 | 25.02 | 19.34 |
| 331 | CSNK1E | 31.46 | 33.00 | 27.04 | 30.50 |  | 6.13 | 11.56 | 13.90 | 10.53 |
| 332 | PLK1 | 32.82 | 34.34 | 24.12 | 30.42 |  | 4.69 | 14.63 | 22.19 | 13.84 |
| 333 | CDK12 | 29.70 | 27.99 | 33.11 | 30.27 |  | 15.86 | 28.80 | 40.05 | 28.24 |
| 334 | CCND3 | 29.69 | 27.24 | 33.64 | 30.19 |  | 7.13 | 12.28 | 16.44 | 11.95 |
| 335 | HK1 | 29.75 | 26.57 | 34.25 | 30.19 |  | 23.93 | 29.03 | 33.83 | 28.93 |
| 336 | HYKK | 29.74 | 25.81 | 34.78 | 30.11 |  | 13.06 | 15.97 | 25.68 | 18.24 |
| 337 | RPS6KA5 | 32.14 | 27.47 | 29.78 | 29.80 |  | 25.23 | 28.33 | 38.18 | 30.58 |
| 338 | FES | 31.97 | 26.56 | 30.39 | 29.64 |  | 22.46 | 25.76 | 35.75 | 27.99 |
| 339 | FN3K | 31.88 | 25.73 | 31.06 | 29.56 |  | 28.49 | 33.51 | 40.59 | 34.20 |
| 340 | PLK4 | 30.41 | 23.51 | 34.51 | 29.48 |  | 18.35 | 15.94 | 38.41 | 24.23 |
| 341 | IKBKE | 30.30 | 22.67 | 35.24 | 29.40 |  | 20.51 | 18.61 | 40.59 | 26.57 |
| 342 | TLK2 | 31.69 | 23.31 | 33.13 | 29.38 |  | 24.77 | 23.39 | 44.87 | 31.01 |
| 343 | KCNH8 | 28.83 | 37.94 | 20.62 | 29.13 |  | 16.13 | 29.84 | 21.68 | 22.55 |
| 344 | GCK | 28.84 | 38.70 | 19.69 | 29.08 |  | 18.23 | 31.78 | 24.47 | 24.83 |
| 345 | CLP1 | 28.76 | 39.36 | 18.67 | 28.93 |  | 0.17 | 26.34 | 32.68 | 19.73 |
| 346 | UCK1 | 28.83 | 32.71 | 25.26 | 28.93 |  | 23.94 | 43.40 | 43.03 | 36.79 |
| 347 | CDK8 | 30.99 | 33.80 | 21.41 | 28.73 |  | 10.37 | 28.08 | 26.98 | 21.81 |
| 348 | PHKG1 | 30.95 | 32.69 | 22.44 | 28.69 |  | 3.96 | 19.93 | 18.09 | 13.99 |
| 349 | DCLK3 | 30.87 | 31.54 | 23.45 | 28.62 |  | 28.01 | 47.31 | 34.59 | 36.64 |
| 350 | PLXNA1 | 29.25 | 28.85 | 27.05 | 28.38 |  | 9.21 | 27.11 | 12.98 | 16.43 |
| 351 | MAPK3 | 29.14 | 27.67 | 28.10 | 28.30 |  | 0.00 | 0.00 | 0.00 | 0.00 |
| 352 | DTYMK | 30.27 | 27.73 | 26.19 | 28.07 |  | 20.04 | 22.64 | 30.67 | 24.45 |
| 353 | FN3KRP | 27.87 | 24.27 | 31.59 | 27.91 |  | 31.35 | 33.11 | 38.61 | 34.36 |
| 354 | EPHA1 | 27.86 | 23.18 | 32.45 | 27.83 |  | 25.82 | 27.07 | 30.37 | 27.75 |
| 355 | CDKN1A | 27.77 | 22.02 | 33.22 | 27.67 |  | 13.54 | 13.95 | 14.72 | 14.07 |
| 356 | TRIB3 | 27.84 | 21.02 | 34.16 | 27.67 |  | 21.63 | 21.20 | 19.43 | 20.75 |
| 357 | CDK19 | 29.68 | 21.79 | 31.07 | 27.52 |  | 23.07 | 26.87 | 27.65 | 25.86 |
| 358 | INSRR | 29.59 | 20.64 | 32.08 | 27.44 |  | 28.78 | 32.08 | 30.65 | 30.50 |
| 359 | CNKSR3 | 29.59 | 19.57 | 33.15 | 27.44 |  | 30.05 | 24.54 | 12.62 | 22.41 |
| 360 | CDK1 | 28.20 | 17.11 | 36.88 | 27.40 |  | 31.14 | 24.56 | 9.88 | 21.86 |
| 361 | CDK9 | 27.90 | 15.73 | 37.74 | 27.12 |  | 48.34 | 40.68 | 23.24 | 37.42 |
| 362 | EFEMP1 | 28.92 | 15.69 | 35.82 | 26.81 |  | 42.74 | 20.88 | 26.95 | 30.19 |
| 363 | TYK2 | 26.80 | 12.50 | 40.41 | 26.57 |  | 49.16 | 26.01 | 29.77 | 34.98 |
| 364 | GUK1 | 26.79 | 19.92 | 32.77 | 26.49 |  | 18.18 | 15.97 | 39.67 | 24.61 |
| 365 | AK2 | 26.53 | 27.88 | 24.09 | 26.16 |  | 28.52 | 17.87 | 32.10 | 26.16 |
| 366 | PPP5K2 | 26.35 | 17.11 | 34.29 | 25.91 |  | 6.04 | 5.74 | 29.32 | 13.70 |
| 367 | IPMK | 27.90 | 22.55 | 27.14 | 25.87 |  | 33.14 | 28.73 | 47.18 | 36.35 |
| 368 | CDK5 | 27.90 | 26.44 | 23.25 | 25.86 |  | 29.63 | 21.09 | 34.42 | 28.38 |
| 369 | CCNB1 | 27.81 | 20.25 | 29.29 | 25.79 |  | 12.21 | 9.56 | 27.76 | 16.51 |
| 370 | CMPK2 | 26.21 | 38.99 | 11.21 | 25.47 |  | 29.80 | 19.72 | 29.48 | 26.34 |
| 371 | JAK2 | 25.87 | 37.89 | 11.72 | 25.16 |  | 31.11 | 29.59 | 23.49 | 28.07 |
| 372 | PRKACA | 26.97 | 38.22 | 9.81 | 25.00 |  | 23.07 | 18.74 | 19.52 | 20.44 |
| 373 | ADRBK2 | 25.61 | 36.11 | 13.21 | 24.98 |  | 34.65 | 30.00 | 32.68 | 32.44 |
| 374 | FAM20C | 25.19 | 34.94 | 14.16 | 24.76 |  | 18.84 | 13.87 | 18.46 | 17.06 |
| 375 | CMPK1 | 25.26 | 34.24 | 14.79 | 24.76 |  | 21.33 | 16.04 | 22.53 | 19.97 |
| 376 | CSNK2B | 25.46 | 33.69 | 14.74 | 24.63 |  | 24.28 | 18.67 | 27.07 | 23.34 |
| 377 | FER | 26.57 | 34.03 | 13.29 | 24.63 |  | 20.75 | 14.82 | 25.12 | 20.23 |
| 378 | HGS | 26.54 | 33.25 | 14.03 | 24.61 |  | 34.69 | 28.44 | 40.64 | 34.59 |
| 379 | ZC3HC1 | 26.54 | 32.49 | 14.79 | 24.61 |  | 22.45 | 16.22 | 30.67 | 23.11 |
| 380 | CKMT1A | 25.21 | 30.40 | 17.98 | 24.53 |  | 27.85 | 21.30 | 37.65 | 28.93 |
| 381 | TWF1 | 25.18 | 29.61 | 18.80 | 24.53 |  | 32.38 | 25.51 | 43.76 | 33.88 |
| 382 | DCK | 26.37 | 30.05 | 16.93 | 24.45 |  | 18.16 | 16.04 | 41.27 | 25.16 |
| 383 | CCNE1 | 26.29 | 29.20 | 17.62 | 24.37 |  | 3.17 | 1.07 | 28.54 | 10.93 |
| 384 | EPHA7 | 26.20 | 31.32 | 15.36 | 24.29 |  | 24.75 | 11.41 | 29.87 | 22.01 |
| 385 | EPHB1 | 26.20 | 27.17 | 19.51 | 24.29 |  | 16.80 | 15.70 | 26.23 | 19.58 |
| 386 | TTC33 | 26.20 | 27.18 | 19.50 | 24.29 |  | 29.93 | 29.35 | 37.19 | 32.15 |
| 387 | SGK3 | 26.14 | 27.13 | 19.43 | 24.23 |  | 16.63 | 16.56 | 21.71 | 18.30 |
| 388 | TSSK1B | 26.12 | 27.53 | 18.99 | 24.21 |  | 13.56 | 14.42 | 17.30 | 15.09 |
| 389 | ILK | 26.09 | 28.31 | 18.16 | 24.18 |  | 20.78 | 22.95 | 23.93 | 22.55 |
| 390 | DGKQ | 26.03 | 29.06 | 17.31 | 24.14 |  | 18.89 | 22.38 | 21.46 | 20.91 |
| 391 | HIPK4 | 13.74 | 17.57 | 41.10 | 24.14 |  | 31.07 | 35.87 | 33.06 | 33.33 |
| 392 | CSNK1G1 | 25.87 | 30.52 | 15.57 | 23.99 |  | 27.71 | 33.82 | 29.12 | 30.22 |
| 393 | FGFR1 | 25.78 | 31.23 | 14.69 | 23.90 |  | 8.88 | 16.31 | 9.71 | 11.64 |
| 394 | ITPKB | 25.61 | 31.86 | 13.75 | 23.74 |  | 16.69 | 25.43 | 16.93 | 19.68 |
| 395 | CALM3 | 25.44 | 32.51 | 12.81 | 23.58 |  | 9.10 | 32.29 | 35.03 | 25.47 |
| 396 | CDK15 | 14.27 | 22.14 | 34.35 | 23.58 |  | 5.40 | 30.13 | 31.21 | 22.25 |
| 397 | DGKZ | 25.35 | 11.05 | 34.11 | 23.51 |  | 29.49 | 32.77 | 9.20 | 23.82 |
| 398 | KCNH2 | 14.12 | 21.00 | 35.21 | 23.44 |  | 20.98 | 32.43 | 13.81 | 22.40 |
| 399 | CSK | 25.18 | 23.84 | 21.03 | 23.35 |  | 18.84 | 39.24 | 26.36 | 28.14 |
| 400 | MAPK15 | 13.86 | 23.10 | 33.09 | 23.35 |  | 7.90 | 8.45 | 2.51 | 6.29 |
| 401 | AAK1 | 25.18 | 30.53 | 14.33 | 23.35 |  | 30.92 | 46.09 | 21.57 | 32.86 |
| 402 | CAD | 25.10 | 26.55 | 18.16 | 23.27 |  | 21.23 | 41.03 | 17.92 | 26.73 |
| 403 | NRBP1 | 24.80 | 32.37 | 11.82 | 23.00 |  | 24.25 | 16.42 | 29.21 | 23.29 |
| 404 | CAMK4 | 13.48 | 0.70 | 54.45 | 22.88 |  | 24.25 | 19.43 | 10.57 | 18.08 |
| 405 | HIPK1 | 24.59 | 12.57 | 31.23 | 22.80 |  | 25.97 | 10.99 | 23.72 | 20.23 |
| 406 | MAP4K2 | 24.51 | 13.25 | 30.41 | 22.72 |  | 21.67 | 11.01 | 16.38 | 16.35 |
| 407 | KCNH5 | 24.34 | 13.84 | 29.52 | 22.56 |  | 32.37 | 31.09 | 34.17 | 32.55 |
| 408 | KDR | 24.00 | 14.25 | 28.49 | 22.25 |  | 19.04 | 12.41 | 28.46 | 19.97 |
| 409 | CSNK1A1 | 13.89 | 22.87 | 29.86 | 22.21 |  | 27.28 | 33.45 | 1.29 | 20.67 |
| 410 | PRKAG3 | 23.74 | 31.97 | 10.33 | 22.01 |  | 28.74 | 34.34 | 3.89 | 22.33 |
| 411 | TSSK3 | 13.69 | 21.16 | 31.18 | 22.01 |  | 29.73 | 34.77 | 6.03 | 23.51 |
| 412 | EPHB2 | 23.57 | 30.28 | 11.71 | 21.86 |  | 27.49 | 31.96 | 4.94 | 21.46 |
| 413 | IQCH | 13.32 | 19.27 | 32.84 | 21.81 |  | 19.33 | 12.24 | 8.93 | 13.50 |
| 414 | COL4A3BP | 23.40 | 28.60 | 13.09 | 21.70 |  | 33.47 | 36.81 | 13.21 | 27.83 |
| 415 | FAM20B | 23.40 | 27.84 | 13.85 | 21.70 |  | 29.89 | 32.68 | 10.78 | 24.45 |
| 416 | TAF9 | 23.36 | 30.59 | 11.03 | 21.66 |  | 37.82 | 36.50 | 12.77 | 29.03 |
| 417 | CDK3 | 13.14 | 20.07 | 31.41 | 21.54 |  | 37.13 | 34.78 | 12.29 | 28.07 |
| 418 | WEE1 | 23.04 | 29.69 | 11.36 | 21.36 |  | 25.42 | 22.02 | 0.78 | 16.07 |
| 419 | CKMT1B | 22.94 | 27.10 | 13.76 | 21.27 |  | 25.53 | 23.29 | 5.48 | 18.10 |
| 420 | IRAK3 | 22.78 | 27.02 | 13.56 | 21.12 |  | 20.50 | 30.00 | 26.20 | 25.57 |
| 421 | CAMK2D | 22.64 | 26.19 | 14.14 | 20.99 |  | 16.67 | 18.42 | 31.18 | 22.09 |
| 422 | HCK | 13.04 | 16.67 | 31.37 | 20.36 |  | 23.93 | 26.01 | 36.85 | 28.93 |
| 423 | TSSK4 | 21.96 | 25.67 | 13.45 | 20.36 |  | 15.38 | 14.85 | 29.68 | 19.97 |
| 424 | HMGXB3 | 12.86 | 16.64 | 31.03 | 20.18 |  | 19.56 | 24.70 | 26.95 | 23.74 |
| 425 | NADK | 21.71 | 25.57 | 13.10 | 20.13 |  | 4.41 | 9.15 | 10.96 | 8.18 |
| 426 | MAP3K5 | 12.32 | 17.02 | 30.10 | 19.81 |  | 0.96 | 0.62 | 0.31 | 0.63 |
| 427 | TRPM7 | 21.28 | 26.05 | 11.86 | 19.73 |  | 0.01 | 0.05 | 0.18 | 0.08 |
| 428 | CSNK1A1L | 20.86 | 26.09 | 11.07 | 19.34 |  | 21.88 | 25.49 | 22.44 | 23.27 |
| 429 | ERN1 | 20.86 | 26.47 | 10.69 | 19.34 |  | 26.98 | 31.02 | 25.97 | 27.99 |
| 430 | CDC42BPB | 11.86 | 16.25 | 29.67 | 19.26 |  | 23.59 | 29.31 | 23.52 | 25.47 |
| 431 | CDK18 | 20.61 | 25.07 | 11.63 | 19.10 |  | 2.93 | 9.05 | 1.23 | 4.40 |
| 432 | CDC42SE2 | 20.61 | 25.15 | 11.56 | 19.10 |  | 11.56 | 23.14 | 18.37 | 17.69 |
| 433 | CDK16 | 20.61 | 25.22 | 11.48 | 19.10 |  | 18.47 | 30.79 | 24.33 | 24.53 |
| 434 | EGFR | 20.54 | 22.73 | 13.86 | 19.04 |  | 21.68 | 28.93 | 14.98 | 21.86 |
| 435 | TSSK2 | 12.55 | 14.51 | 29.54 | 18.87 |  | 17.60 | 25.33 | 9.43 | 17.45 |
| 436 | GRK1 | 20.32 | 22.05 | 14.15 | 18.84 |  | 18.98 | 32.80 | 3.72 | 18.50 |
| 437 | CLK1 | 12.47 | 13.97 | 29.93 | 18.79 |  | 17.48 | 9.80 | 22.73 | 16.67 |
| 438 | CDK13 | 20.10 | 21.36 | 14.44 | 18.63 |  | 27.72 | 27.92 | 24.08 | 26.57 |
| 439 | KSR2 | 11.91 | 12.94 | 30.34 | 18.40 |  | 21.68 | 10.97 | 27.97 | 20.20 |
| 440 | EPHA2 | 18.49 | 19.28 | 13.65 | 17.14 |  | 19.49 | 12.34 | 21.23 | 17.69 |
| 441 | AURKC | 18.40 | 17.84 | 14.94 | 17.06 |  | 22.34 | 18.75 | 19.52 | 20.20 |
| 442 | DGUOK | 18.40 | 18.07 | 14.71 | 17.06 |  | 20.47 | 10.44 | 23.10 | 18.00 |
| 443 | ALPK3 | 10.50 | 10.40 | 29.80 | 16.90 |  | 20.02 | 29.99 | 1.64 | 17.22 |
| 444 | GK2 | 18.19 | 20.46 | 11.94 | 16.86 |  | 21.37 | 32.39 | 7.36 | 20.38 |
| 445 | HUNK | 18.15 | 13.03 | 19.30 | 16.82 |  | 20.26 | 35.71 | 17.38 | 24.45 |
| 446 | ADCK5 | 17.98 | 23.50 | 8.52 | 16.67 |  | 7.16 | 21.11 | 3.56 | 10.61 |
| 447 | AKAP8 | 17.89 | 18.90 | 12.97 | 16.59 |  | 10.22 | 27.59 | 15.73 | 17.85 |
| 448 | CCND1 | 11.27 | 11.92 | 26.58 | 16.59 |  | 2.41 | 19.05 | 8.73 | 10.06 |
| 449 | CCNK | 17.89 | 19.52 | 12.35 | 16.59 |  | 12.78 | 27.33 | 17.20 | 19.10 |
| 450 | CDC42BPA | 11.03 | 13.43 | 24.60 | 16.35 |  | 8.92 | 21.62 | 11.91 | 14.15 |
| 451 | CKB | 17.47 | 20.62 | 10.50 | 16.19 |  | 14.50 | 25.34 | 16.06 | 18.63 |
| 452 | IKBKB | 10.63 | 14.53 | 23.19 | 16.12 |  | 12.96 | 29.77 | 28.73 | 23.82 |
| 453 | NRP2 | 16.87 | 21.54 | 8.52 | 15.64 |  | 2.54 | 0.71 | 10.20 | 4.48 |
| 454 | KHK | 16.11 | 21.53 | 7.16 | 14.94 |  | 17.46 | 14.21 | 24.27 | 18.64 |
| 455 | PIK3C3 | 16.03 | 22.21 | 6.34 | 14.86 |  | 5.33 | 1.80 | 9.14 | 5.42 |
| 456 | ADK | 9.38 | 16.32 | 18.64 | 14.78 |  | 19.16 | 11.23 | 24.09 | 18.16 |
| 457 | CDK4 | 15.86 | 16.41 | 11.84 | 14.70 |  | 5.13 | 8.61 | 32.01 | 15.25 |
| 458 | FASTK | 15.69 | 16.70 | 11.24 | 14.54 |  | 10.46 | 15.96 | 33.95 | 20.13 |
| 459 | EIF2AK3 | 15.43 | 16.39 | 11.10 | 14.31 |  | 5.40 | 4.49 | 16.30 | 8.73 |
| 460 | DDR1 | 15.35 | 14.32 | 13.02 | 14.23 |  | 19.42 | 16.65 | 24.54 | 20.20 |
| 461 | PIM2 | 9.83 | 9.04 | 23.58 | 14.15 |  | 6.55 | 3.37 | 11.08 | 7.00 |
| 462 | NME3 | 15.18 | 14.61 | 12.43 | 14.07 |  | 7.82 | 4.23 | 11.76 | 7.94 |
| 463 | GRK6 | 9.68 | 9.34 | 22.96 | 13.99 |  | 19.40 | 15.40 | 22.75 | 19.18 |
| 464 | MAP4K3 | 14.84 | 14.74 | 11.70 | 13.76 |  | 20.36 | 23.67 | 15.41 | 19.81 |
| 465 | AGK | 9.11 | 11.38 | 20.31 | 13.60 |  | 18.79 | 24.74 | 16.61 | 20.05 |
| 466 | DCLK2 | 14.50 | 16.75 | 9.08 | 13.44 |  | 4.75 | 10.19 | 1.33 | 5.42 |
| 467 | PDGFRB | 14.33 | 16.56 | 8.97 | 13.29 |  | 4.78 | 1.10 | 8.74 | 4.87 |
| 468 | CCL5 | 14.08 | 14.77 | 10.31 | 13.05 |  | 16.00 | 19.60 | 6.38 | 13.99 |
| 469 | DLG3 | 8.65 | 15.41 | 15.09 | 13.05 |  | 12.07 | 19.96 | 14.67 | 15.57 |
| 470 | CDK6 | 13.91 | 14.76 | 10.01 | 12.89 |  | 6.71 | 2.92 | 0.75 | 3.46 |
| 471 | CDK2 | 13.91 | 14.70 | 10.07 | 12.89 |  | 7.37 | 21.68 | 22.36 | 17.14 |
| 472 | ALK | 13.82 | 14.50 | 10.12 | 12.81 |  | 8.46 | 22.39 | 22.46 | 17.77 |
| 473 | CARD11 | 13.74 | 14.62 | 9.85 | 12.74 |  | 6.07 | 17.44 | 22.24 | 15.25 |
| 474 | AKAP14 | 9.42 | 10.51 | 18.28 | 12.74 |  | 4.34 | 6.68 | 3.14 | 4.72 |
| 475 | CAMK1G | 13.40 | 14.70 | 9.16 | 12.42 |  | 1.70 | 8.96 | 11.04 | 7.23 |
| 476 | NTRK1 | 9.10 | 10.62 | 17.54 | 12.42 |  | 2.12 | 8.18 | 9.75 | 6.68 |
| 477 | PLK2 | 13.40 | 15.12 | 8.74 | 12.42 |  | 1.44 | 8.50 | 11.76 | 7.23 |
| 478 | ADCK4 | 8.70 | 10.63 | 17.23 | 12.19 |  | 2.69 | 12.27 | 18.76 | 11.24 |
| 479 | DYRK4 | 13.14 | 15.29 | 8.12 | 12.19 |  | 6.93 | 12.20 | 15.07 | 11.40 |
| 480 | BCKDK | 12.38 | 14.74 | 7.32 | 11.48 |  | 10.39 | 21.05 | 11.48 | 14.31 |
| 481 | PIK3CG | 12.30 | 14.86 | 7.04 | 11.40 |  | 8.58 | 19.13 | 8.14 | 11.95 |
| 482 | MUSK | 7.84 | 10.62 | 15.27 | 11.24 |  | 2.64 | 0.88 | 0.25 | 1.26 |
| 483 | CNTLN | 12.13 | 15.11 | 6.49 | 11.24 |  | 1.90 | 17.66 | 9.21 | 9.59 |
| 484 | ALPK1 | 12.04 | 15.24 | 6.21 | 11.16 |  | 14.50 | 8.51 | 18.03 | 13.68 |
| 485 | AKAP13 | 12.04 | 15.45 | 6.00 | 11.16 |  | 2.02 | 9.53 | 10.15 | 7.23 |
| 486 | CDK20 | 11.96 | 13.23 | 8.07 | 11.08 |  | 1.89 | 6.07 | 5.25 | 4.40 |
| 487 | ACVRL1 | 8.69 | 10.03 | 14.31 | 11.01 |  | 11.19 | 14.24 | 14.19 | 13.21 |
| 488 | CASS4 | 11.70 | 13.11 | 7.74 | 10.85 |  | 3.27 | 5.20 | 5.91 | 4.80 |
| 489 | AMHR2 | 8.37 | 9.85 | 13.86 | 10.69 |  | 2.20 | 3.00 | 4.47 | 3.22 |
| 490 | MVK | 11.53 | 13.07 | 7.47 | 10.69 |  | 8.43 | 8.11 | 10.35 | 8.96 |
| 491 | CAMKK2 | 8.05 | 9.65 | 13.91 | 10.53 |  | 4.21 | 8.28 | 0.25 | 4.25 |
| 492 | ALDH18A1 | 11.28 | 12.95 | 7.14 | 10.46 |  | 3.76 | 1.42 | 4.72 | 3.30 |
| 493 | PHKA2 | 11.19 | 12.94 | 7.00 | 10.38 |  | 8.74 | 10.51 | 4.10 | 7.78 |
| 494 | ADCK2 | 7.58 | 8.37 | 14.00 | 9.98 |  | 1.12 | 1.73 | 0.46 | 1.10 |
| 495 | AURKA | 10.60 | 12.09 | 6.79 | 9.83 |  | 1.04 | 2.50 | 3.77 | 2.44 |
| 496 | CDK5R1 | 10.60 | 12.48 | 6.40 | 9.83 |  | 1.42 | 3.01 | 20.10 | 8.18 |
| 497 | BCR | 10.51 | 12.46 | 6.27 | 9.75 |  | 2.31 | 1.34 | 2.49 | 2.04 |
| 498 | CHEK1 | 10.51 | 8.50 | 10.23 | 9.75 |  | 8.52 | 16.13 | 18.75 | 14.47 |
| 499 | BMX | 8.43 | 6.35 | 14.47 | 9.75 |  | 3.65 | 7.09 | 7.42 | 6.05 |
| 500 | PIP5KL1 | 10.43 | 15.28 | 3.30 | 9.67 |  | 0.70 | 3.55 | 5.18 | 3.14 |
| 501 | PAPSS1 | 8.19 | 13.12 | 7.23 | 9.51 |  | 0.41 | 0.50 | 6.40 | 2.44 |
| 502 | AURKB | 9.92 | 14.92 | 2.75 | 9.20 |  | 6.96 | 6.42 | 12.33 | 8.57 |
| 503 | PAK2 | 7.55 | 6.97 | 12.60 | 9.04 |  | 1.70 | 0.86 | 1.92 | 1.49 |
| 504 | NUAK1 | 9.67 | 12.56 | 4.66 | 8.96 |  | 1.58 | 0.76 | 3.09 | 1.81 |
| 505 | AK1 | 9.50 | 12.15 | 4.76 | 8.81 |  | 5.52 | 6.03 | 14.86 | 8.81 |
| 506 | EPHA5 | 9.50 | 11.92 | 5.00 | 8.81 |  | 0.71 | 5.98 | 20.20 | 8.96 |
| 507 | BMPR2 | 7.17 | 9.36 | 9.18 | 8.57 |  | 2.27 | 3.02 | 13.34 | 6.21 |
| 508 | MAPK14 | 8.73 | 10.69 | 4.86 | 8.10 |  | 1.94 | 1.05 | 4.09 | 2.36 |
| 509 | AURKAIP1 | 8.39 | 10.12 | 4.83 | 7.78 |  | 2.90 | 1.72 | 4.34 | 2.99 |
| 510 | ATR | 8.39 | 9.89 | 5.06 | 7.78 |  | 11.33 | 9.75 | 12.17 | 11.08 |
| 511 | AVP | 7.97 | 6.71 | 7.49 | 7.39 |  | 0.13 | 0.85 | 0.67 | 0.55 |
| 512 | AKAP7 | 6.84 | 6.15 | 8.47 | 7.15 |  | 4.65 | 2.62 | 5.00 | 4.09 |
| 513 | AK4 | 7.63 | 7.24 | 6.36 | 7.08 |  | 2.10 | 0.28 | 1.39 | 1.26 |
| 514 | DOK1 | 6.27 | 6.17 | 7.84 | 6.76 |  | 8.27 | 5.55 | 7.65 | 7.15 |
| 515 | AK8 | 6.95 | 7.14 | 5.25 | 6.45 |  | 3.12 | 0.05 | 2.02 | 1.73 |
| 516 | PIM3 | 6.61 | 7.09 | 4.69 | 6.13 |  | 8.42 | 5.02 | 6.84 | 6.76 |
| 517 | BRSK2 | 6.27 | 7.04 | 4.14 | 5.82 |  | 5.40 | 1.65 | 3.33 | 3.46 |
| 518 | C19orf35 | 5.42 | 6.47 | 5.56 | 5.82 |  | 2.93 | 7.83 | 3.39 | 4.72 |
| 519 | DSTYK | 6.02 | 7.36 | 3.36 | 5.58 |  | 2.39 | 0.95 | 1.37 | 1.57 |
| 520 | AATK | 0.34 | 1.97 | 0.00 | 0.77 |  | 0.00 | 0.00 | 0.00 | 0.00 |
| 521 | AKT1 | 0.17 | 2.09 | 0.00 | 0.75 |  | 0.16 | 0.07 | 0.24 | 0.16 |
| 522 | ACVR2B | 0.00 | 0.00 | 0.00 | 0.00 |  | 0.48 | 2.94 | 8.37 | 3.93 |
| 523 | CHKB | 0.00 | 0.00 | 0.00 | 0.00 |  | 25.41 | 23.52 | 39.28 | 29.40 |
| 524 | TSKS | 0.00 | 0.00 | 0.00 | 0.00 |  | 0.00 | 0.00 | 0.00 | 0.00 |

**Table 2.** Primary screening score from high content analyser system in SCC4 Cas9 cells: % Cell viability of accepted (green dot 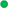) knockout groups (cell viability >=65%) in absence and presence of docetaxel. This is in reference to figure 1B.

| Sl. | Kinases | Cell viability of Ko alone<br>>= 65% |  |  | Average<br>% |  | Cell viability of Ko + Doc<br>treatment > 30% |  |  | Average<br>% |
| --- | --- | --- | --- | --- | --- | --- | --- | --- | --- | --- |
| 1 | MYO3A | 148.87 | 127.49 | 131.00 | 135.79 |  | 87.00 | 100.65 | 84.02 | 90.55 |
| 2 | MPP6 | 126.00 | 140.49 | 132.98 | 133.16 |  | 61.71 | 74.80 | 68.68 | 68.40 |
| 3 | PNKP | 132.83 | 130.84 | 128.49 | 130.72 |  | 69.48 | 82.00 | 77.01 | 76.16 |
| 4 | STK19 | 133.29 | 129.11 | 128.19 | 130.20 |  | 50.31 | 62.27 | 52.41 | 55.00 |
| 5 | PFKFB1 | 132.20 | 130.20 | 125.52 | 129.31 |  | 50.61 | 62.00 | 53.27 | 55.29 |
| 6 | FRK | 126.18 | 135.05 | 125.75 | 128.99 |  | 60.94 | 71.77 | 62.34 | 65.02 |
| 7 | TAOK1 | 127.45 | 111.05 | 144.58 | 127.70 |  | 56.41 | 66.67 | 58.37 | 60.48 |
| 8 | TRIB1 | 125.33 | 136.35 | 115.76 | 125.82 |  | 32.24 | 41.95 | 34.56 | 36.25 |
| 9 | DGKD | 121.24 | 110.57 | 142.37 | 124.73 |  | 90.80 | 99.94 | 93.68 | 94.81 |
| 10 | WNK3 | 123.34 | 101.42 | 146.00 | 123.58 |  | 41.60 | 52.64 | 41.60 | 45.28 |
| 11 | TSSK6 | 122.85 | 93.30 | 153.14 | 123.10 |  | 18.28 | 29.32 | 18.28 | 21.96 |
| 12 | CERKL | 122.51 | 108.46 | 138.02 | 123.00 |  | 61.67 | 72.35 | 61.67 | 65.23 |
| 13 | PANK2 | 117.65 | 97.51 | 151.46 | 122.21 |  | 67.21 | 77.89 | 67.21 | 70.77 |
| 14 | CNKSR1 | 121.47 | 110.61 | 133.05 | 121.71 |  | 37.00 | 46.03 | 37.00 | 40.01 |
| 15 | ZMYND8 | 121.17 | 103.60 | 140.07 | 121.61 |  | 67.12 | 76.15 | 67.12 | 70.13 |
| 16 | SH3BP5 | 116.84 | 111.31 | 135.80 | 121.32 |  | 61.38 | 70.42 | 61.38 | 64.39 |
| 17 | PFKL | 120.97 | 104.15 | 138.53 | 121.22 |  | 85.42 | 90.44 | 85.42 | 87.09 |
| 18 | PIK3C2B | 120.19 | 110.14 | 131.70 | 120.67 |  | 76.37 | 81.39 | 76.37 | 78.04 |
| 19 | CCL8 | 113.18 | 97.54 | 139.28 | 116.67 |  | 68.77 | 73.79 | 68.77 | 70.44 |
| 20 | FUK | 116.26 | 105.59 | 127.68 | 116.51 |  | 54.14 | 59.17 | 54.14 | 55.82 |
| 21 | MAP3K8 | 116.17 | 98.90 | 134.18 | 116.42 |  | 23.04 | 27.07 | 23.04 | 24.38 |
| 22 | CERK | 115.83 | 111.15 | 121.98 | 116.32 |  | 81.59 | 85.63 | 81.59 | 82.94 |
| 23 | NEK11 | 111.57 | 97.67 | 139.14 | 116.12 |  | 72.20 | 76.23 | 72.20 | 73.54 |
| 24 | MAP2K4 | 114.05 | 96.80 | 132.03 | 114.29 |  | 42.47 | 46.51 | 42.47 | 43.82 |
| 25 | PHKG2 | 113.36 | 115.94 | 112.09 | 113.80 |  | 79.30 | 88.61 | 71.11 | 79.67 |
| 26 | SBK1 | 108.57 | 105.12 | 125.46 | 113.05 |  | 74.00 | 82.97 | 66.15 | 74.37 |
| 27 | TRIB2 | 112.12 | 101.96 | 123.02 | 112.36 |  | 32.37 | 41.00 | 24.85 | 32.74 |
| 28 | MAP3K14 | 110.15 | 98.07 | 123.68 | 110.63 |  | 82.37 | 90.66 | 75.19 | 82.74 |
| 29 | PACSIN2 | 106.70 | 112.80 | 111.06 | 110.19 |  | 74.52 | 82.48 | 66.59 | 74.53 |
| 30 | MAP3K10 | 109.59 | 108.08 | 111.85 | 109.84 |  | 64.08 | 71.71 | 56.50 | 64.09 |
| 31 | CAMK2G | 109.58 | 100.22 | 119.69 | 109.83 |  | 68.78 | 76.06 | 61.53 | 68.79 |
| 32 | PFKFB2 | 109.16 | 104.08 | 115.70 | 109.64 |  | 75.61 | 82.55 | 68.69 | 75.62 |
| 33 | PLXNA2 | 104.84 | 112.06 | 111.29 | 109.40 |  | 50.79 | 57.41 | 54.86 | 54.35 |
| 34 | EIF2AK2 | 109.11 | 108.47 | 110.46 | 109.35 |  | 43.00 | 49.28 | 48.96 | 47.08 |
| 35 | STK11IP | 108.76 | 107.98 | 110.85 | 109.20 |  | 27.45 | 33.38 | 33.53 | 31.45 |
| 36 | CCL2 | 104.25 | 107.64 | 114.30 | 108.73 |  | 67.22 | 77.89 | 68.57 | 71.23 |
| 37 | MAP2K6 | 108.11 | 117.61 | 99.35 | 108.36 |  | 64.42 | 75.10 | 64.78 | 68.10 |
| 38 | CDK10 | 106.98 | 114.02 | 101.41 | 107.47 |  | 68.49 | 70.87 | 77.15 | 72.17 |
| 39 | NRP1 | 103.09 | 109.12 | 107.52 | 106.58 |  | 42.43 | 44.24 | 60.67 | 49.11 |
| 40 | MAPK6 | 105.83 | 104.80 | 107.61 | 106.08 |  | 57.56 | 58.81 | 76.37 | 64.24 |
| 41 | MAP3K7 | 105.74 | 95.06 | 117.16 | 105.98 |  | 31.22 | 31.90 | 44.59 | 35.91 |
| 42 | STK40 | 105.40 | 93.16 | 119.09 | 105.89 |  | 52.59 | 52.70 | 66.52 | 57.27 |
| 43 | GTF2H4 | 101.18 | 89.16 | 126.88 | 105.74 |  | 46.79 | 46.34 | 59.47 | 50.86 |
| 44 | CAMKK1 | 104.48 | 92.45 | 117.22 | 104.72 |  | 57.68 | 56.68 | 69.37 | 61.24 |
| 45 | PAK3 | 104.11 | 93.07 | 116.47 | 104.55 |  | 64.89 | 63.32 | 71.49 | 66.57 |
| 46 | STYK1 | 99.63 | 88.59 | 124.10 | 104.10 |  | 70.04 | 67.90 | 77.20 | 71.71 |
| 47 | SLK | 103.71 | 83.65 | 124.51 | 103.96 |  | 50.75 | 48.05 | 58.47 | 52.42 |
| 48 | GNE | 103.13 | 83.08 | 124.64 | 103.62 |  | 43.15 | 39.89 | 50.45 | 44.50 |
| 49 | CCNH | 99.58 | 85.53 | 124.09 | 103.07 |  | 71.69 | 81.00 | 66.42 | 73.03 |
| 50 | TNIK | 102.27 | 88.21 | 117.07 | 102.52 |  | 23.18 | 32.15 | 18.25 | 24.53 |
| 51 | LMTK3 | 102.22 | 89.99 | 115.21 | 102.47 |  | 68.94 | 77.58 | 70.98 | 72.50 |
| 52 | TYRO3 | 101.89 | 91.21 | 114.02 | 102.37 |  | 36.31 | 44.60 | 37.04 | 39.32 |
| 53 | EIF2AK4 | 97.02 | 91.99 | 115.71 | 101.57 |  | 33.31 | 41.27 | 34.38 | 36.32 |
| 54 | NMRK2 | 101.14 | 96.12 | 106.89 | 101.38 |  | 24.83 | 32.45 | 26.24 | 27.84 |
| 55 | PIK3CD | 100.89 | 95.87 | 107.24 | 101.34 |  | 59.01 | 66.29 | 56.75 | 60.68 |
| 56 | RET | 96.62 | 92.59 | 114.09 | 101.10 |  | 34.11 | 41.06 | 31.20 | 35.46 |
| 57 | MAP3K4 | 100.84 | 96.81 | 105.62 | 101.09 |  | 64.61 | 71.22 | 59.12 | 64.99 |
| 58 | MPP1 | 100.55 | 96.52 | 106.04 | 101.04 |  | 77.22 | 83.50 | 72.07 | 77.60 |
| 59 | PANK1 | 97.55 | 86.88 | 118.68 | 101.04 |  | 94.14 | 95.26 | 94.14 | 94.51 |
| 60 | MAP2K3 | 100.64 | 109.68 | 92.35 | 100.89 |  | 61.74 | 62.86 | 61.74 | 62.12 |
| 61 | BUB1 | 100.38 | 109.42 | 92.09 | 100.63 |  | 62.25 | 62.29 | 62.25 | 62.26 |
| 62 | PRKCD | 100.06 | 109.09 | 92.48 | 100.54 |  | 59.88 | 59.91 | 59.88 | 59.89 |
| 63 | STAP2 | 95.84 | 100.86 | 104.49 | 100.40 |  | 32.53 | 32.57 | 32.53 | 32.54 |
| 64 | PDGFRL | 100.11 | 104.14 | 96.80 | 100.35 |  | 76.74 | 76.78 | 76.74 | 76.76 |
| 65 | PAN3 | 99.41 | 100.53 | 99.61 | 99.85 |  | 86.50 | 94.81 | 86.50 | 89.27 |
| 66 | CAMK2N1 | 95.05 | 96.17 | 107.36 | 99.53 |  | 70.36 | 74.53 | 83.88 | 76.26 |
| 67 | MARK1 | 99.26 | 100.38 | 98.88 | 99.51 |  | 50.93 | 55.14 | 64.41 | 56.82 |
| 68 | CDK11B | 98.88 | 100.01 | 99.22 | 99.37 |  | 58.05 | 62.30 | 65.49 | 61.95 |
| 69 | SCYL3 | 95.72 | 95.76 | 106.14 | 99.21 |  | 38.64 | 42.92 | 46.04 | 42.53 |
| 70 | NIM1K | 98.81 | 98.85 | 99.52 | 99.06 |  | 23.37 | 27.69 | 28.91 | 26.66 |
| 71 | HIPK2 | 98.41 | 98.45 | 99.13 | 98.66 |  | 49.23 | 53.59 | 54.73 | 52.52 |
| 72 | MPP7 | 98.03 | 98.07 | 99.45 | 98.52 |  | 76.46 | 80.85 | 81.71 | 79.67 |
| 73 | MAP4K5 | 93.81 | 102.12 | 99.18 | 98.37 |  | 43.32 | 47.75 | 48.54 | 46.54 |
| 74 | MAST4 | 98.03 | 115.71 | 81.07 | 98.27 |  | 62.49 | 66.96 | 66.69 | 65.38 |
| 75 | PGK2 | 97.58 | 115.27 | 81.22 | 98.02 |  | 63.38 | 67.89 | 67.54 | 66.27 |
| 76 | RAF1 | 93.49 | 105.18 | 95.24 | 97.97 |  | 47.81 | 52.36 | 48.79 | 49.65 |
| 77 | NTSC2 | 97.38 | 109.06 | 86.44 | 97.63 |  | 79.81 | 84.40 | 80.75 | 81.65 |
| 78 | PIM1 | 96.84 | 106.71 | 88.43 | 97.33 |  | 61.12 | 65.74 | 60.38 | 62.41 |
| 79 | PLXNB3 | 93.60 | 103.46 | 94.19 | 97.08 |  | 75.71 | 80.37 | 74.93 | 77.00 |
| 80 | SHC1 | 96.64 | 106.29 | 87.73 | 96.88 |  | 61.66 | 66.36 | 60.85 | 62.96 |
| 81 | MAGI2 | 96.44 | 106.09 | 87.53 | 96.69 |  | 63.10 | 67.83 | 58.24 | 63.06 |
| 82 | PIP4K2B | 95.66 | 104.32 | 88.45 | 96.14 |  | 58.50 | 63.27 | 53.60 | 58.46 |
| 83 | MAP3K6 | 91.59 | 100.25 | 96.59 | 96.14 |  | 19.68 | 24.49 | 14.74 | 19.63 |
| 84 | MKNK2 | 95.80 | 101.32 | 91.00 | 96.04 |  | 77.14 | 81.99 | 72.17 | 77.10 |
| 85 | PDK1 | 95.35 | 100.88 | 91.16 | 95.80 |  | 73.62 | 78.50 | 67.61 | 73.24 |
| 86 | CAMK1 | 90.96 | 94.84 | 100.51 | 95.44 |  | 80.56 | 85.49 | 74.52 | 80.19 |
| 87 | LRPPRC | 95.10 | 98.99 | 91.97 | 95.35 |  | 60.22 | 65.18 | 54.13 | 59.84 |
| 88 | KIAA1804 | 94.52 | 98.40 | 92.09 | 95.00 |  | 71.84 | 76.84 | 65.72 | 71.46 |
| 89 | SCGB2A1 | 91.42 | 91.29 | 102.01 | 94.91 |  | 59.70 | 64.74 | 50.63 | 58.36 |
| 90 | SQSTM1 | 94.41 | 94.28 | 95.28 | 94.65 |  | 76.50 | 81.57 | 67.40 | 75.16 |
| 91 | MAPK13 | 94.21 | 94.08 | 95.09 | 94.46 |  | 52.43 | 57.54 | 43.29 | 51.09 |
| 92 | PIK3AP1 | 93.97 | 93.85 | 95.56 | 94.46 |  | 62.42 | 67.57 | 53.24 | 61.08 |
| 93 | FYN | 89.85 | 88.73 | 104.65 | 94.41 |  | 68.42 | 73.61 | 58.12 | 66.72 |
| 94 | MYLK4 | 94.07 | 92.95 | 95.91 | 94.31 |  | 65.45 | 70.68 | 55.11 | 63.75 |
| 95 | SGK223 | 93.66 | 92.54 | 96.11 | 94.10 |  | 62.48 | 67.74 | 52.10 | 60.77 |
| 96 | PRKCQ | 89.54 | 88.42 | 104.09 | 94.02 |  | 76.63 | 81.93 | 66.22 | 74.93 |
| 97 | STK32B | 93.72 | 89.69 | 98.49 | 93.97 |  | 44.01 | 49.34 | 44.19 | 45.85 |
| 98 | PBK | 93.43 | 89.40 | 98.92 | 93.92 |  | 59.61 | 64.98 | 61.32 | 61.97 |
| 99 | SCYL2 | 90.38 | 86.35 | 104.87 | 93.87 |  | 44.05 | 49.46 | 45.51 | 46.34 |
| 100 | BUB1B | 93.54 | 89.51 | 98.32 | 93.79 |  | 82.38 | 87.83 | 83.80 | 84.67 |
| 101 | GSAP | 93.52 | 88.40 | 99.38 | 93.77 |  | 38.44 | 43.93 | 38.84 | 40.41 |
| 102 | PRKCG | 92.74 | 87.62 | 99.31 | 93.22 |  | 56.25 | 61.77 | 56.61 | 58.21 |
| 103 | PFKP | 88.60 | 83.49 | 107.39 | 93.16 |  | 42.75 | 48.32 | 52.09 | 47.72 |
| 104 | MAP4K1 | 92.84 | 87.72 | 98.67 | 93.08 |  | 62.10 | 67.70 | 71.40 | 67.06 |
| 105 | LRRK2 | 92.24 | 97.76 | 88.04 | 92.68 |  | 73.34 | 78.98 | 76.61 | 76.31 |
| 106 | EPHB3 | 87.90 | 94.98 | 94.25 | 92.37 |  | 42.32 | 47.99 | 45.54 | 45.28 |
| 107 | FLT1 | 92.05 | 98.92 | 85.92 | 92.30 |  | 46.46 | 52.17 | 47.83 | 48.82 |
| 108 | MATK | 91.55 | 98.42 | 86.14 | 92.04 |  | 69.82 | 75.57 | 69.59 | 71.66 |
| 109 | PHKA1 | 88.50 | 94.39 | 93.08 | 91.99 |  | 44.36 | 50.14 | 38.44 | 44.31 |
| 110 | FGGY | 91.69 | 97.57 | 86.55 | 91.94 |  | 67.85 | 73.67 | 61.89 | 67.80 |
| 111 | RPS6KA4 | 91.42 | 106.32 | 77.26 | 91.67 |  | 69.38 | 75.25 | 63.39 | 69.34 |
| 112 | NEK1 | 91.11 | 106.01 | 77.66 | 91.59 |  | 50.03 | 55.93 | 43.01 | 49.65 |
| 113 | STK38 | 87.04 | 95.94 | 91.80 | 91.59 |  | 46.52 | 52.45 | 39.46 | 46.14 |
| 114 | LIMK1 | 90.86 | 99.76 | 82.68 | 91.10 |  | 69.66 | 68.54 | 69.66 | 69.29 |
| 115 | MAST1 | 90.36 | 97.44 | 84.60 | 90.80 |  | 20.96 | 26.48 | 20.96 | 22.80 |
| 116 | PRKCDBP | 86.27 | 91.80 | 94.19 | 90.75 |  | 54.12 | 55.26 | 54.12 | 54.50 |
| 117 | MYLK | 90.16 | 90.03 | 91.03 | 90.41 |  | 69.60 | 73.71 | 66.33 | 69.88 |
| 118 | GUCY2C | 89.53 | 89.40 | 91.12 | 90.02 |  | 48.92 | 54.17 | 44.79 | 49.29 |
| 119 | MAPK4 | 86.43 | 86.30 | 97.01 | 89.91 |  | 51.63 | 58.00 | 58.47 | 56.03 |
| 120 | PMVK | 89.66 | 88.54 | 91.53 | 89.91 |  | 61.20 | 68.71 | 54.51 | 61.47 |
| 121 | PLXNB2 | 89.27 | 88.15 | 91.13 | 89.52 |  | 60.40 | 61.10 | 60.40 | 60.63 |
| 122 | EFNB3 | 89.03 | 87.91 | 91.61 | 89.52 |  | 43.93 | 55.17 | 43.93 | 47.68 |
| 123 | PRPS1 | 84.91 | 90.43 | 93.06 | 89.47 |  | 52.84 | 54.27 | 52.84 | 53.31 |
| 124 | MARK4 | 88.63 | 89.77 | 88.21 | 88.87 |  | 71.77 | 72.92 | 71.77 | 72.16 |
| 125 | DOLK | 88.43 | 89.27 | 88.92 | 88.87 |  | 77.11 | 78.12 | 77.11 | 77.45 |
| 126 | PDK3 | 84.25 | 85.35 | 96.58 | 88.72 |  | 93.16 | 93.80 | 93.16 | 93.37 |
| 127 | WNK2 | 88.28 | 95.22 | 82.08 | 88.53 |  | 59.39 | 60.16 | 59.39 | 59.64 |
| 128 | MAP3K3 | 87.64 | 88.45 | 88.29 | 88.13 |  | 77.25 | 78.13 | 77.25 | 77.55 |
| 129 | DYRK2 | 84.64 | 85.34 | 94.40 | 88.13 |  | 42.08 | 42.96 | 42.08 | 42.37 |
| 130 | PANK3 | 87.59 | 94.47 | 81.45 | 87.83 |  | 70.45 | 71.26 | 70.45 | 70.72 |
| 131 | PNCK | 87.49 | 88.92 | 86.80 | 87.74 |  | 69.63 | 70.42 | 69.63 | 69.89 |
| 132 | MPP5 | 87.10 | 88.25 | 87.41 | 87.59 |  | 74.77 | 75.53 | 74.77 | 75.02 |
| 133 | PKDCC | 82.83 | 83.84 | 95.49 | 87.39 |  | 49.53 | 50.36 | 49.51 | 49.80 |
| 134 | KCNH4 | 87.10 | 87.75 | 87.17 | 87.34 |  | 43.61 | 44.46 | 53.92 | 47.33 |
| 135 | MAP2K5 | 86.90 | 87.67 | 87.45 | 87.34 |  | 60.79 | 61.65 | 60.49 | 60.98 |
| 136 | OXSRI | 82.81 | 83.69 | 95.37 | 87.29 |  | 71.92 | 72.79 | 72.06 | 72.26 |
| 137 | PRKCI | 86.69 | 87.57 | 86.57 | 86.94 |  | 62.39 | 63.27 | 62.46 | 62.71 |
| 138 | PANK4 | 86.41 | 87.21 | 87.06 | 86.89 |  | 59.32 | 60.21 | 59.11 | 59.55 |
| 139 | SGMS2 | 83.41 | 84.20 | 93.07 | 86.89 |  | 59.83 | 60.74 | 74.24 | 64.94 |
| 140 | MAPK11 | 86.62 | 87.39 | 86.60 | 86.87 |  | 48.54 | 49.45 | 48.47 | 48.82 |
| 141 | FKBP1A | 86.54 | 87.37 | 86.47 | 86.79 |  | 38.26 | 39.19 | 38.12 | 38.52 |
| 142 | PKLR | 86.23 | 93.01 | 80.91 | 86.71 |  | 35.47 | 36.41 | 35.20 | 35.69 |
| 143 | PIK3CA | 81.99 | 82.55 | 95.11 | 86.55 |  | 40.26 | 41.21 | 40.19 | 40.55 |
| 144 | N4BP2 | 86.26 | 87.27 | 85.97 | 86.50 |  | 49.36 | 50.32 | 48.39 | 49.36 |
| 145 | MAPKAPK5 | 86.06 | 87.01 | 86.43 | 86.50 |  | 57.72 | 58.69 | 56.74 | 57.72 |
| 146 | POMK | 81.87 | 82.56 | 94.62 | 86.35 |  | 66.04 | 67.03 | 66.33 | 66.47 |
| 147 | NAGK | 85.95 | 92.71 | 79.94 | 86.20 |  | 42.27 | 43.27 | 41.90 | 42.48 |
| 148 | DYRK3 | 85.67 | 86.52 | 86.27 | 86.15 |  | 22.65 | 23.66 | 21.64 | 22.65 |
| 149 | PIK3CB | 82.52 | 83.32 | 92.18 | 86.00 |  | 72.45 | 73.47 | 72.19 | 72.70 |
| 150 | MARK3 | 85.51 | 83.32 | 88.45 | 85.76 |  | 61.37 | 62.41 | 60.34 | 61.37 |
| 151 | TRRAP | 85.26 | 86.18 | 85.08 | 85.51 |  | 25.39 | 26.43 | 25.18 | 25.67 |
| 152 | SRPK1 | 84.78 | 86.15 | 84.86 | 85.26 |  | 56.49 | 57.20 | 56.49 | 56.73 |
| 153 | TIE1 | 80.51 | 84.78 | 89.91 | 85.06 |  | 46.11 | 46.94 | 46.11 | 46.38 |
| 154 | MAPKAPK2 | 84.77 | 80.51 | 89.76 | 85.01 |  | 46.09 | 46.09 | 46.09 | 46.09 |
| 155 | PACSIN1 | 84.54 | 86.05 | 84.36 | 84.98 |  | 33.30 | 34.11 | 33.30 | 33.57 |
| 156 | PLXNA3 | 80.39 | 85.18 | 89.04 | 84.87 |  | 56.10 | 57.08 | 56.10 | 56.43 |
| 157 | GAK | 84.58 | 80.39 | 89.51 | 84.83 |  | 60.86 | 61.54 | 60.86 | 61.08 |
| 158 | GRK5 | 83.95 | 85.34 | 84.01 | 84.43 |  | 57.52 | 58.30 | 57.52 | 57.78 |
| 159 | LATS2 | 80.79 | 83.95 | 88.08 | 84.27 |  | 85.36 | 86.11 | 85.36 | 85.61 |
| 160 | NPR2 | 83.79 | 81.63 | 86.70 | 84.04 |  | 37.03 | 37.03 | 37.03 | 37.03 |
| 161 | PFKM | 83.68 | 84.50 | 83.60 | 83.93 |  | 74.38 | 74.38 | 74.38 | 74.38 |
| 162 | PRKCSH | 83.34 | 84.51 | 83.64 | 83.83 |  | 22.80 | 23.38 | 22.80 | 23.00 |
| 163 | PGK1 | 79.17 | 83.34 | 88.67 | 83.73 |  | 63.73 | 64.53 | 63.73 | 64.00 |
| 164 | NLK | 83.49 | 79.98 | 87.71 | 83.73 |  | 22.56 | 23.01 | 22.11 | 22.56 |
| 165 | MAP2K1 | 83.09 | 84.46 | 83.04 | 83.53 |  | 61.75 | 62.18 | 62.42 | 62.12 |
| 166 | STK24 | 78.56 | 83.78 | 86.77 | 83.04 |  | 64.64 | 65.05 | 64.23 | 64.64 |
| 167 | TWF2 | 82.69 | 79.34 | 86.79 | 82.94 |  | 19.30 | 19.70 | 19.49 | 19.50 |
| 168 | CASK | 82.45 | 83.44 | 82.92 | 82.94 |  | 51.89 | 52.26 | 52.45 | 52.20 |
| 169 | CAMK2B | 79.38 | 82.45 | 86.76 | 82.86 |  | 54.40 | 54.76 | 54.05 | 54.40 |
| 170 | GUCY2D | 82.46 | 79.38 | 86.28 | 82.70 |  | 48.09 | 48.43 | 48.53 | 48.35 |
| 171 | SGK1 | 82.39 | 83.03 | 82.50 | 82.64 |  | 63.80 | 64.11 | 64.37 | 64.09 |
| 172 | NEK7 | 82.06 | 83.20 | 82.38 | 82.55 |  | 21.85 | 22.15 | 22.51 | 22.17 |
| 173 | ULK1 | 77.98 | 82.06 | 87.58 | 82.54 |  | 22.35 | 22.62 | 23.13 | 22.70 |
| 174 | TRPM6 | 82.07 | 79.09 | 85.78 | 82.31 |  | 74.37 | 74.63 | 74.11 | 74.37 |
| 175 | DGKI | 81.63 | 82.07 | 82.52 | 82.08 |  | 54.31 | 54.54 | 54.83 | 54.56 |
| 176 | PIKFYVE | 77.42 | 82.21 | 86.07 | 81.90 |  | 83.83 | 84.05 | 83.61 | 83.83 |
| 177 | ATM | 81.59 | 78.36 | 85.57 | 81.84 |  | 48.51 | 48.71 | 48.78 | 48.66 |
| 178 | ERBB4 | 80.88 | 81.59 | 81.63 | **81.37** |  | 62.05 | 62.24 | 62.50 | **62.26** |
| 179 | PRKAA2 | 77.82 | 81.65 | 84.44 | **81.31** |  | 52.92 | 53.08 | 53.50 | **53.17** |
| 180 | ABL2 | 80.86 | 78.71 | 83.76 | **81.11** |  | 49.95 | 50.09 | 49.81 | **49.95** |
| 181 | TAF1 | 80.73 | 81.82 | 80.38 | **80.97** |  | 30.73 | 30.86 | 31.33 | **30.97** |
| 182 | SEPHS2 | 80.28 | 81.78 | 80.23 | **80.76** |  | 54.25 | 54.36 | 54.15 | **54.25** |
| 183 | ERCC2 | 76.10 | 80.28 | 85.60 | **80.66** |  | 44.18 | 44.27 | 44.81 | **44.42** |
| 184 | PRKACB | 80.32 | 76.86 | 84.50 | **80.56** |  | 40.94 | 41.00 | 41.51 | **41.15** |
| 185 | ROR2 | 80.06 | 80.32 | 81.12 | **80.50** |  | 51.16 | 51.20 | 51.88 | **51.42** |
| 186 | TGFA | 75.71 | 80.53 | 84.33 | **80.19** |  | 33.41 | 33.44 | 34.32 | **33.73** |
| 187 | PIP4K2C | 79.94 | 76.34 | 84.29 | **80.19** |  | 44.90 | 45.58 | 44.90 | **45.13** |
| 188 | PAK4 | 79.47 | 80.68 | 79.71 | **79.95** |  | 40.36 | 41.21 | 40.36 | **40.64** |
| 189 | NTRK2 | 76.31 | 79.47 | 83.61 | **79.80** |  | 24.76 | 24.76 | 24.76 | **24.76** |
| 190 | NEK3 | 79.52 | 77.03 | 82.76 | **79.77** |  | 29.13 | 29.13 | 29.13 | **29.13** |
| 191 | PAK6 | 79.39 | 79.52 | 80.00 | **79.64** |  | 50.25 | 50.92 | 50.25 | **50.47** |
| 192 | SEPHS1 | 79.14 | 80.10 | 79.64 | **79.62** |  | 34.49 | 35.02 | 34.49 | **34.67** |
| 193 | IP6K2 | 74.92 | 79.77 | 83.73 | **79.48** |  | 76.01 | 76.01 | 76.01 | **76.01** |
| 194 | PEAK1 | 79.24 | 75.69 | 83.50 | **79.48** |  | 89.60 | 90.24 | 89.60 | **89.81** |
| 195 | NRBP2 | 78.96 | 80.17 | 79.08 | **79.40** |  | 30.02 | 30.77 | 30.02 | **30.27** |
| 196 | DGKK | 74.77 | 79.64 | 83.33 | **79.25** |  | 53.91 | 54.69 | 53.91 | **54.17** |
| 197 | STAP1 | 78.98 | 75.62 | 83.08 | **79.23** |  | 34.39 | 34.92 | 34.39 | **34.57** |
| 198 | STK32A | 78.69 | 78.98 | 79.86 | **79.18** |  | 31.95 | 31.95 | 31.95 | **31.95** |
| 199 | SGK2 | 75.68 | 78.69 | 83.13 | **79.17** |  | 43.35 | 43.96 | 43.35 | **43.55** |
| 200 | MLKL | 78.88 | 76.35 | 82.16 | **79.13** |  | 29.28 | 29.28 | 29.28 | **29.28** |
| 201 | PFKFB3 | 78.76 | 79.41 | 78.86 | **79.01** |  | 38.01 | 38.60 | 38.01 | **38.21** |
| 202 | PRKDC | 78.49 | 78.76 | 79.69 | **78.98** |  | 45.69 | 46.46 | 45.69 | **45.94** |
| 203 | CDK11A | 74.42 | 79.14 | 83.38 | **78.98** |  | 50.88 | 51.65 | 50.88 | **51.14** |
| 204 | MYLK3 | 78.38 | 75.17 | 82.30 | **78.62** |  | 26.97 | 27.91 | 26.02 | **26.97** |
| 205 | PGM2L1 | 78.05 | 79.15 | 78.26 | **78.49** |  | 69.41 | 70.50 | 69.00 | **69.63** |
| 206 | KIT | 73.81 | 78.58 | 82.48 | **78.29** |  | 78.93 | 80.16 | 77.70 | **78.93** |
| 207 | STK10 | 77.97 | 73.81 | 82.88 | **78.22** |  | 49.70 | 51.07 | 48.99 | **49.92** |
| 208 | PDK2 | 77.26 | 78.58 | 77.40 | **77.74** |  | 53.34 | 54.85 | 52.49 | **53.56** |
| 209 | NUAK2 | 74.16 | 77.26 | 81.52 | **77.65** |  | 38.48 | 40.13 | 37.56 | **38.72** |
| 210 | SNX16 | 77.25 | 74.74 | 80.50 | **77.50** |  | 59.13 | 60.92 | 57.99 | **59.35** |
| 211 | GRK4 | 77.19 | 78.02 | 77.10 | **77.44** |  | 42.85 | 44.78 | 40.92 | **42.85** |
| 212 | SRPK2 | 76.86 | 77.95 | 77.23 | **77.35** |  | 44.91 | 46.98 | 42.84 | **44.91** |
| 213 | MAK | 72.74 | 76.86 | 82.29 | **77.30** |  | 43.17 | 45.38 | 41.71 | **43.42** |
| 214 | SIK3 | 76.96 | 73.42 | 81.22 | **77.20** |  | 65.77 | 68.12 | 64.03 | **65.97** |
| 215 | ROCK2 | 76.60 | 76.96 | 77.57 | **77.04** |  | 56.70 | 59.20 | 55.09 | **57.00** |
| 216 | FASTKD2 | 72.49 | 77.25 | 81.15 | **76.97** |  | 49.47 | 52.10 | 47.49 | **49.69** |
| 217 | PIK3C2A | 76.51 | 73.14 | 80.62 | **76.76** |  | 28.24 | 31.01 | 25.47 | **28.24** |
| 218 | MOS | 76.17 | 77.23 | 76.57 | **76.66** |  | 52.97 | 55.88 | 50.05 | **52.97** |
| 219 | RIOK3 | 73.02 | 76.81 | 79.69 | **76.51** |  | 58.72 | 59.41 | 58.72 | **58.95** |
| 220 | TBCK | 76.26 | 73.02 | 80.24 | **76.51** |  | 51.29 | 52.01 | 51.29 | **51.53** |
| 221 | PLK3 | 75.93 | 76.26 | 76.35 | **76.18** |  | 46.54 | 46.54 | 46.54 | **46.54** |
| 222 | MYO3B | 75.43 | 76.67 | 75.64 | **75.91** |  | 44.46 | 44.46 | 44.46 | **44.46** |
| 223 | NEK9 | 71.07 | 76.04 | 79.77 | **75.63** |  | 21.17 | 21.81 | 21.17 | **21.38** |
| 224 | NME4 | 75.03 | 71.95 | 78.84 | **75.27** |  | 30.28 | 30.98 | 30.28 | **30.51** |
| 225 | BRAF | 74.73 | 75.69 | 75.10 | **75.17** |  | 76.11 | 76.11 | 76.11 | **76.11** |
| 226 | PFKFB4 | 70.69 | 74.73 | 80.09 | 75.17 |  | 24.63 | 25.23 | 24.63 | 24.83 |
| 227 | PDPK1 | 74.91 | 70.69 | 79.87 | 75.16 |  | 42.54 | 43.22 | 42.54 | 42.77 |
| 228 | TESK2 | 74.59 | 75.59 | 75.05 | 75.08 |  | 41.04 | 41.04 | 41.04 | 41.04 |
| 229 | MAP3K2 | 71.51 | 75.31 | 78.18 | 75.00 |  | 79.01 | 79.48 | 79.01 | 79.17 |
| 230 | NPR1 | 74.75 | 71.51 | 78.73 | 75.00 |  | 15.96 | 15.96 | 15.96 | 15.96 |
| 231 | IRAK1 | 74.58 | 74.75 | 75.15 | 74.83 |  | 69.93 | 69.93 | 69.93 | 69.93 |
| 232 | NTRK3 | 74.28 | 75.22 | 74.79 | 74.76 |  | 28.18 | 29.01 | 28.18 | 28.46 |
| 233 | SRC | 70.12 | 74.98 | 78.94 | 74.68 |  | 51.58 | 52.18 | 51.58 | 51.78 |
| 234 | MELK | 74.39 | 70.12 | 79.38 | 74.63 |  | 20.32 | 20.94 | 20.32 | 20.52 |
| 235 | PIK3R2 | 73.85 | 74.99 | 74.04 | 74.29 |  | 41.75 | 41.75 | 41.75 | 41.75 |
| 236 | MET | 69.80 | 74.53 | 78.52 | 74.28 |  | 83.18 | 83.18 | 83.18 | 83.18 |
| 237 | NME8 | 73.98 | 69.80 | 78.91 | 74.23 |  | 44.38 | 45.06 | 44.38 | 44.61 |
| 238 | NME7 | 73.55 | 74.46 | 74.10 | 74.04 |  | 25.47 | 26.36 | 25.47 | 25.77 |
| 239 | GCKR | 70.49 | 73.55 | 77.89 | 73.98 |  | 40.89 | 41.57 | 40.89 | 41.12 |
| 240 | LMTK2 | 73.59 | 70.49 | 77.43 | 73.84 |  | 84.22 | 84.22 | 84.22 | 84.22 |
| 241 | BRSK1 | 73.57 | 74.42 | 73.47 | 73.82 |  | 59.90 | 60.86 | 59.90 | 60.22 |
| 242 | NME9 | 73.30 | 74.17 | 73.90 | 73.79 |  | 29.57 | 30.33 | 29.57 | 29.82 |
| 243 | DGKH | 69.23 | 73.93 | 78.20 | 73.79 |  | 81.15 | 82.22 | 81.15 | 81.50 |
| 244 | IGFN1 | 73.40 | 69.23 | 78.29 | 73.64 |  | 58.55 | 59.90 | 58.55 | 59.00 |
| 245 | IPPK | 73.20 | 73.40 | 74.32 | 73.64 |  | 71.44 | 72.25 | 71.44 | 71.71 |
| 246 | TXK | 68.87 | 73.87 | 77.30 | 73.35 |  | 21.82 | 22.63 | 21.82 | 22.09 |
| 247 | GLYCTK | 73.10 | 69.76 | 77.19 | 73.35 |  | 32.74 | 41.12 | 32.74 | 35.53 |
| 248 | PRKAB2 | 72.71 | 73.79 | 73.09 | 73.19 |  | 37.67 | 38.16 | 37.67 | 37.83 |
| 249 | STK25 | 69.63 | 72.71 | 77.00 | 73.11 |  | 13.55 | 14.68 | 15.88 | 14.70 |
| 250 | SIK1 | 72.86 | 70.59 | 75.89 | 73.11 |  | 51.95 | 53.16 | 51.26 | 52.12 |
| 251 | PKN3 | 72.80 | 72.03 | 74.31 | 73.05 |  | 58.77 | 60.04 | 58.04 | 58.95 |
| 252 | CIB1 | 72.55 | 71.48 | 75.08 | 73.03 |  | 32.60 | 33.93 | 39.60 | 35.38 |
| 253 | PICK1 | 68.40 | 67.05 | 83.42 | 72.96 |  | 38.03 | 39.43 | 46.36 | 41.27 |
| 254 | PXK | 72.51 | 71.71 | 74.03 | 72.75 |  | 49.21 | 50.68 | 48.32 | 49.41 |
| 255 | PCM1 | 72.26 | 71.45 | 74.39 | 72.70 |  | 69.37 | 70.91 | 68.32 | 69.54 |
| 256 | EFNA4 | 68.17 | 62.81 | 86.97 | 72.65 |  | 55.41 | 57.02 | 67.99 | 60.14 |
| 257 | SCYL1 | 72.30 | 71.81 | 73.54 | 72.55 |  | 43.24 | 44.92 | 52.64 | 46.93 |
| 258 | ITPK1 | 71.77 | 66.13 | 78.87 | 72.26 |  | 50.86 | 52.60 | 49.96 | 51.14 |
| 259 | PIK3R6 | 68.68 | 68.18 | 79.64 | 72.17 |  | 46.21 | 48.02 | 56.23 | 50.16 |
| 260 | NEK8 | 71.76 | 71.22 | 73.05 | 72.01 |  | 27.43 | 34.45 | 27.43 | 29.77 |
| 261 | PIP5K1C | 71.69 | 66.05 | 78.07 | 71.93 |  | 31.73 | 33.22 | 38.35 | 34.43 |
| 262 | STRADB | 71.37 | 65.76 | 78.44 | 71.86 |  | 46.36 | 48.24 | 56.35 | 50.31 |
| 263 | DYRK1B | 67.22 | 66.64 | 81.47 | 71.78 |  | 35.35 | 37.29 | 42.45 | 38.36 |
| 264 | CDKL5 | 71.52 | 71.03 | 72.73 | 71.76 |  | 40.60 | 42.61 | 48.98 | 44.07 |
| 265 | SPEG | 71.26 | 65.65 | 78.18 | 71.70 |  | 60.56 | 62.64 | 73.98 | 65.72 |
| 266 | LIMK2 | 67.18 | 61.90 | 85.90 | 71.66 |  | 75.46 | 77.61 | 92.63 | 81.90 |
| 267 | PINK1 | 71.37 | 70.53 | 72.96 | 71.62 |  | 47.59 | 49.80 | 46.24 | 47.88 |
| 268 | LYN | 70.88 | 65.31 | 77.91 | 71.36 |  | 48.66 | 50.95 | 47.12 | 48.91 |
| 269 | NPRL2 | 67.58 | 62.27 | 83.35 | 71.07 |  | 20.33 | 22.68 | 18.56 | 20.52 |
| 270 | PIK3R3 | 70.67 | 65.12 | 76.97 | 70.92 |  | 61.76 | 64.18 | 60.42 | 62.12 |
| 271 | GTF2H1 | 70.58 | 65.04 | 76.88 | 70.83 |  | 31.64 | 34.13 | 29.75 | 31.84 |
| 272 | PAPSS2 | 70.35 | 64.82 | 77.34 | 70.83 |  | 52.52 | 55.07 | 63.40 | 57.00 |
| 273 | TRIO | 66.26 | 61.05 | 85.14 | 70.82 |  | 66.13 | 81.77 | 51.07 | 66.32 |
| 274 | ALPK2 | 70.51 | 64.97 | 76.78 | **70.75** |  | 49.69 | 52.18 | 59.92 | **53.93** |
| 275 | NEK6 | 70.23 | 64.71 | 77.08 | **70.68** |  | 17.42 | 20.04 | 15.60 | **17.69** |
| 276 | PRKCZ | 66.14 | 65.27 | 80.45 | **70.62** |  | 46.21 | 47.34 | 56.90 | **50.15** |
| 277 | RPS6KB1 | 70.32 | 69.58 | 71.82 | **70.57** |  | 38.67 | 39.87 | 38.96 | **39.17** |
| 278 | ULK3 | 70.03 | 69.45 | 72.07 | **70.52** |  | 30.28 | 31.55 | 36.76 | **32.86** |
| 279 | ETNK2 | 66.88 | 65.79 | 78.42 | **70.36** |  | 17.34 | 18.67 | 16.58 | **17.53** |
| 280 | PI4KB | 70.03 | 69.43 | 71.38 | **70.28** |  | 35.01 | 36.42 | 34.23 | **35.22** |
| 281 | PACSIN3 | 70.03 | 64.53 | 76.29 | **70.28** |  | 28.25 | 29.72 | 34.01 | **30.66** |
| 282 | PDK4 | 69.72 | 69.13 | 71.76 | **70.20** |  | 28.61 | 30.15 | 34.40 | **31.05** |
| 283 | GTF2H3 | 65.49 | 60.34 | 84.31 | **70.05** |  | 45.31 | 46.92 | 44.10 | **45.44** |
| 284 | TJP2 | 69.02 | 68.21 | 70.55 | **69.26** |  | 54.77 | 56.45 | 53.40 | **54.87** |
| 285 | AKT2 | 68.82 | 63.41 | 75.55 | **69.26** |  | 58.53 | 60.27 | 71.77 | **63.52** |
| 286 | BTK | 64.70 | 63.21 | 79.63 | **69.18** |  | 64.69 | 66.50 | 79.43 | **70.20** |
| 287 | GK | 68.84 | 63.43 | 75.00 | **69.09** |  | 33.71 | 35.59 | 33.67 | **34.32** |
| 288 | BMP2K | 68.46 | 67.88 | 70.50 | **68.95** |  | 46.50 | 48.45 | 56.46 | **50.47** |
| 289 | NME5 | 65.46 | 64.84 | 76.53 | **68.94** |  | 31.31 | 33.32 | 37.30 | **33.98** |
| 290 | STK4 | 68.64 | 63.25 | 74.79 | **68.89** |  | 48.44 | 50.52 | 58.76 | **52.57** |
| 291 | ZAP70 | 68.62 | 63.22 | 74.76 | **68.87** |  | 35.57 | 37.72 | 42.52 | **38.60** |
| 292 | RNASEL | 68.36 | 67.96 | 70.21 | **68.84** |  | 43.52 | 47.16 | 51.02 | **47.23** |
| 293 | MKNK1 | 64.19 | 63.89 | 78.16 | **68.74** |  | 72.77 | 76.48 | 87.69 | **78.98** |
| 294 | STK17B | 68.45 | 63.07 | 74.56 | **68.69** |  | 48.67 | 52.44 | 57.35 | **52.82** |
| 295 | AK9 | 67.95 | 62.61 | 74.62 | **68.40** |  | 47.01 | 50.85 | 55.20 | **51.02** |
| 296 | STK33 | 63.72 | 61.88 | 79.00 | **68.20** |  | 40.71 | 44.62 | 37.52 | **40.95** |
| 297 | PAK7 | 67.91 | 62.57 | 74.00 | **68.16** |  | 52.83 | 56.81 | 49.55 | **53.07** |
| 298 | STK39 | 67.37 | 62.07 | 74.12 | **67.85** |  | 31.33 | 35.37 | 27.96 | **31.55** |
| 299 | GALK2 | 64.36 | 59.30 | 79.88 | **67.85** |  | 41.61 | 51.47 | 32.40 | **41.82** |
| 300 | AKT3 | 67.21 | 61.93 | 73.24 | **67.46** |  | 28.62 | 33.12 | 24.76 | **28.83** |
| 301 | SLAMF6 | 67.16 | 61.88 | 73.19 | **67.41** |  | 39.28 | 43.39 | 45.22 | **42.63** |
| 302 | OBSCN | 66.57 | 61.34 | 73.27 | **67.06** |  | 22.35 | 26.53 | 19.75 | **22.88** |
| 303 | PSKH2 | 62.31 | 57.41 | 80.88 | **66.86** |  | 54.91 | 59.16 | 64.72 | **59.59** |
| 304 | SGMS1 | 66.58 | 61.35 | 72.54 | **66.82** |  | 40.28 | 44.59 | 37.77 | **40.88** |
| 305 | CDK17 | 66.38 | 65.67 | 68.42 | **66.82** |  | 48.75 | 49.36 | 60.61 | **52.91** |
| 306 | RBKS | 61.95 | 61.26 | 76.08 | **66.43** |  | 40.68 | 41.55 | 42.30 | **41.51** |
| 307 | SNRK | 66.17 | 65.50 | 67.59 | **66.42** |  | 49.85 | 50.23 | 62.23 | **54.10** |
| 308 | IRAK4 | 65.49 | 64.83 | 67.60 | **65.97** |  | 50.70 | 63.49 | 48.57 | **54.25** |
| 309 | NRGN | 62.14 | 61.51 | 73.23 | **65.63** |  | 66.59 | 82.97 | 58.75 | **69.44** |
| 310 | NRK | 65.08 | 59.97 | 70.95 | **65.33** |  | 54.85 | 55.97 | 61.29 | **57.37** |
| 311 | EPHA6 | 65.08 | 63.50 | 67.41 | **65.33** |  | 40.81 | 50.95 | 37.25 | **43.00** |
| 312 | ATMIN | 64.77 | 59.67 | 71.32 | **65.25** |  | 65.74 | 66.13 | 70.96 | **67.61** |
| 313 | CDC42BPG | 60.63 | 58.82 | 76.10 | **65.18** |  | 45.34 | 45.61 | 49.70 | **46.88** |
| 314 | CAMKV | 64.93 | 59.83 | 70.76 | **65.17** |  | 59.08 | 59.24 | 62.58 | **60.30** |
| 315 | GTF2H2 | 64.73 | 62.24 | 68.55 | **65.17** |  | 38.80 | 48.17 | 32.13 | **39.70** |
| 316 | GALK1 | 60.69 | 55.92 | 78.90 | **65.17** |  | 36.77 | 41.76 | 33.49 | **37.34** |

| Sl. | Kinases | Average live cell count |  |  |
| --- | --- | --- | --- | --- |
| 1 | -ve SgRNA | 1499.43 | 1537.19 | 1911.43 |
| 2 | -ve SgRNA + 2nM Docetaxel | 1104.02 | 1269.44 | 1443.94 |

Each count of control group is mean of live cells from 6 individual wells

**Table 3.**
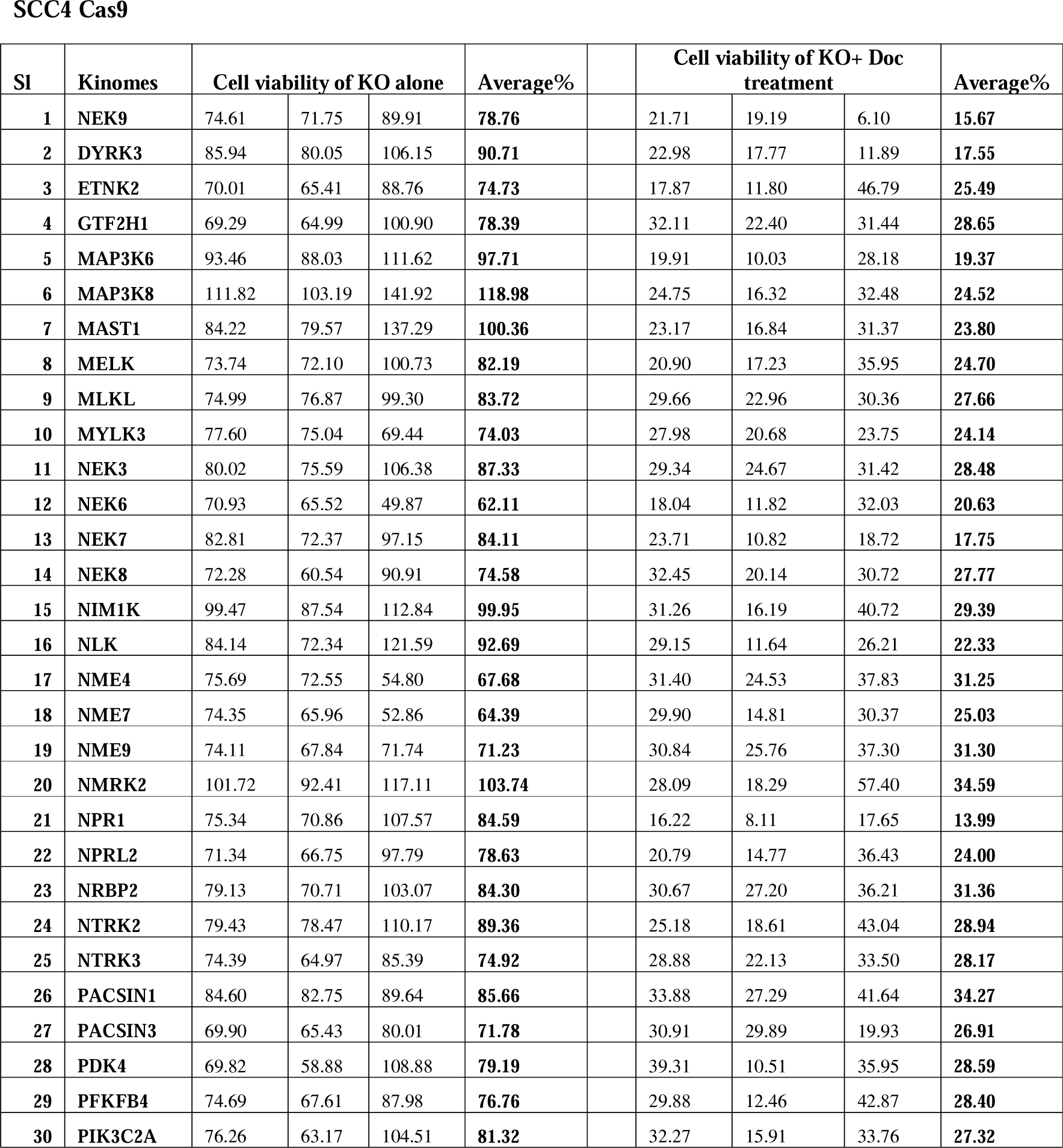
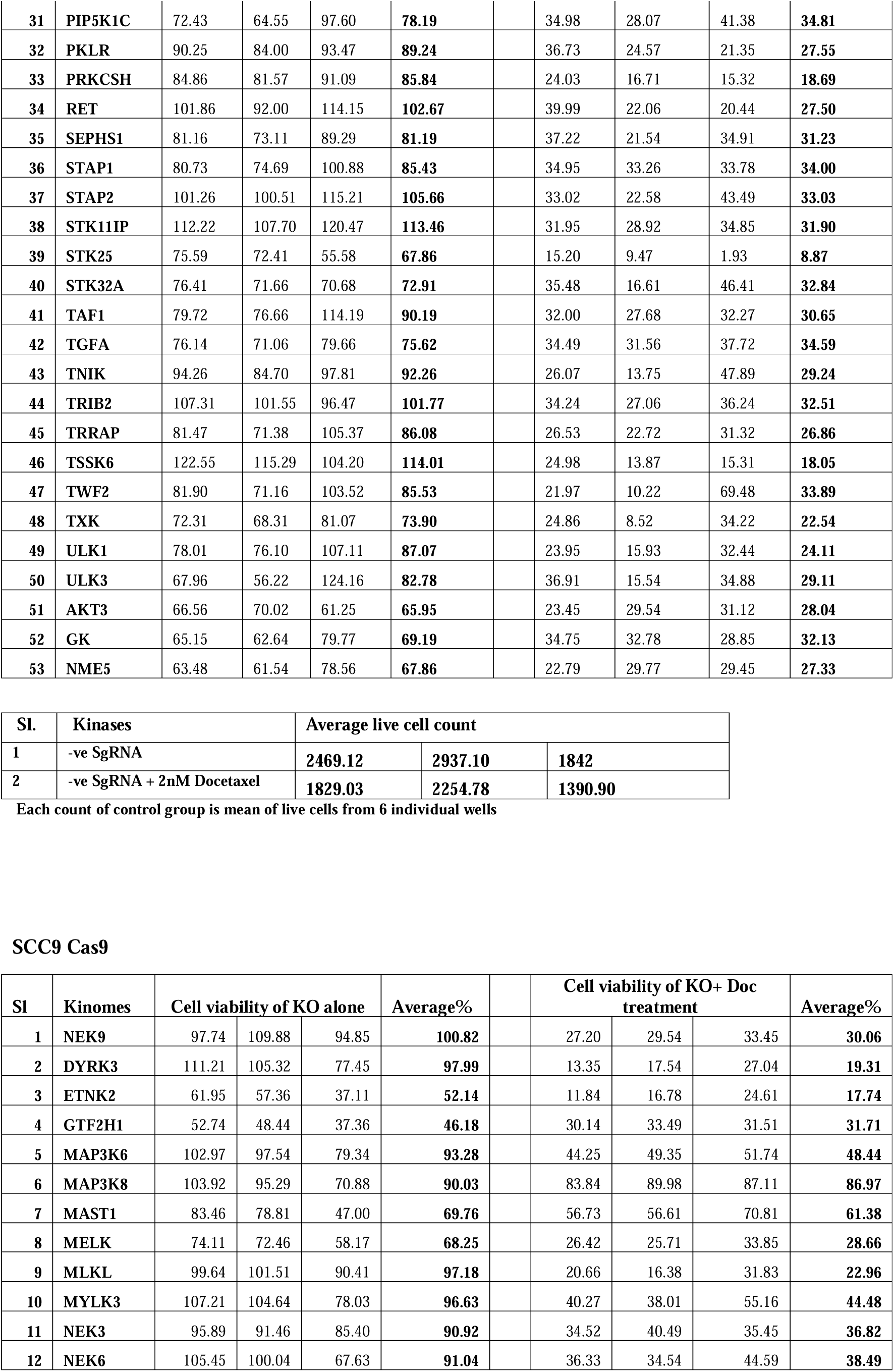
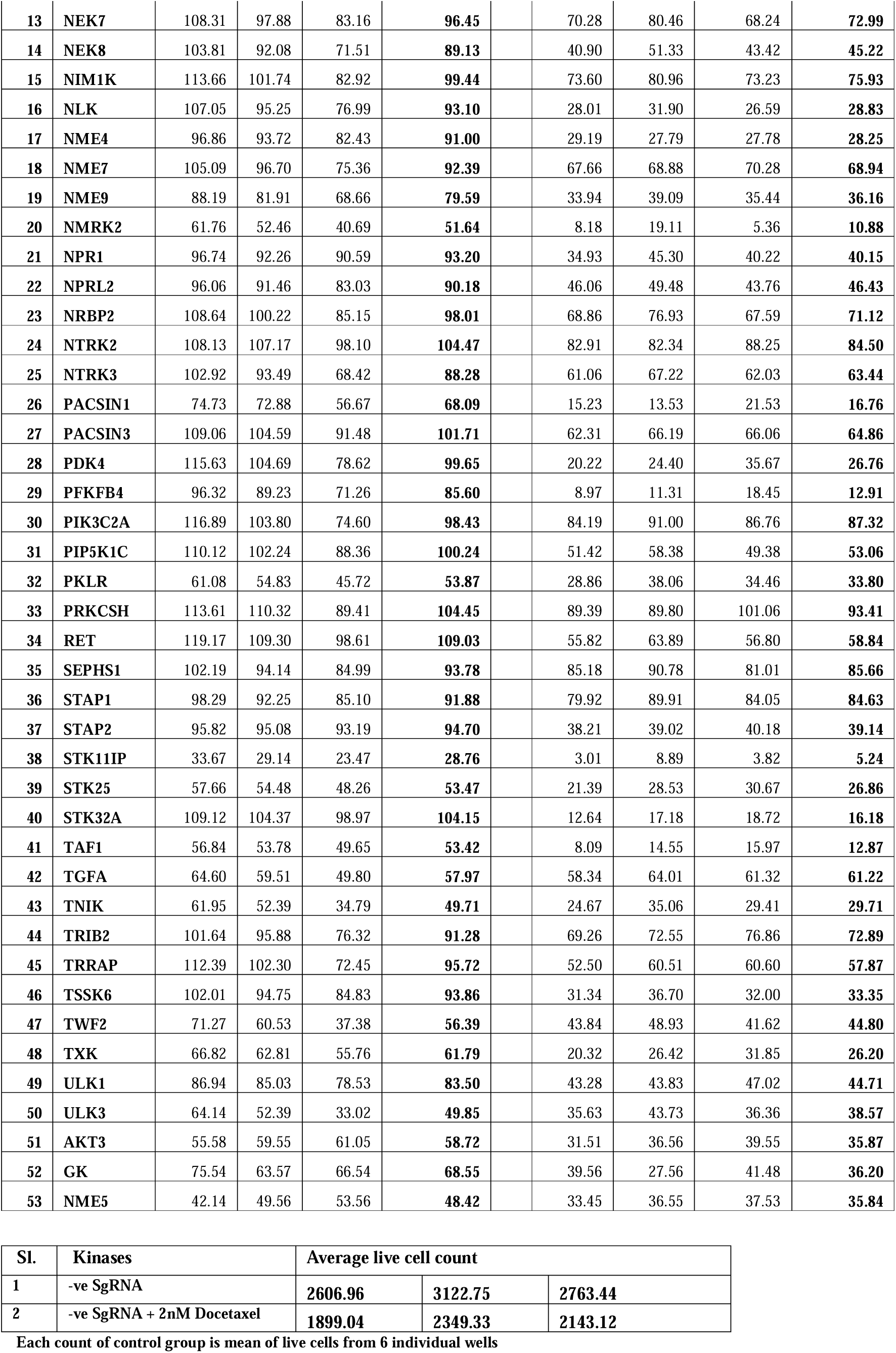
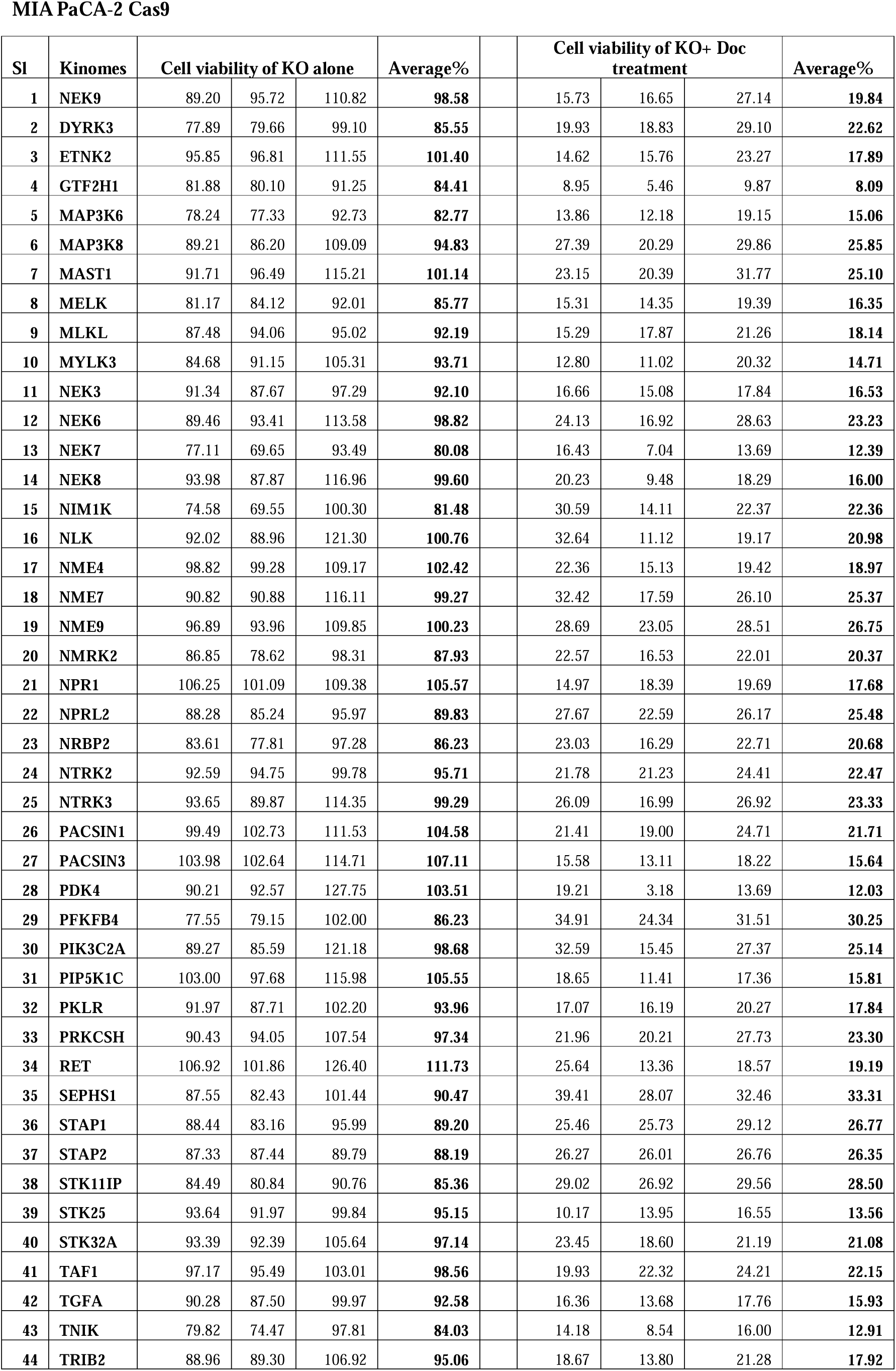
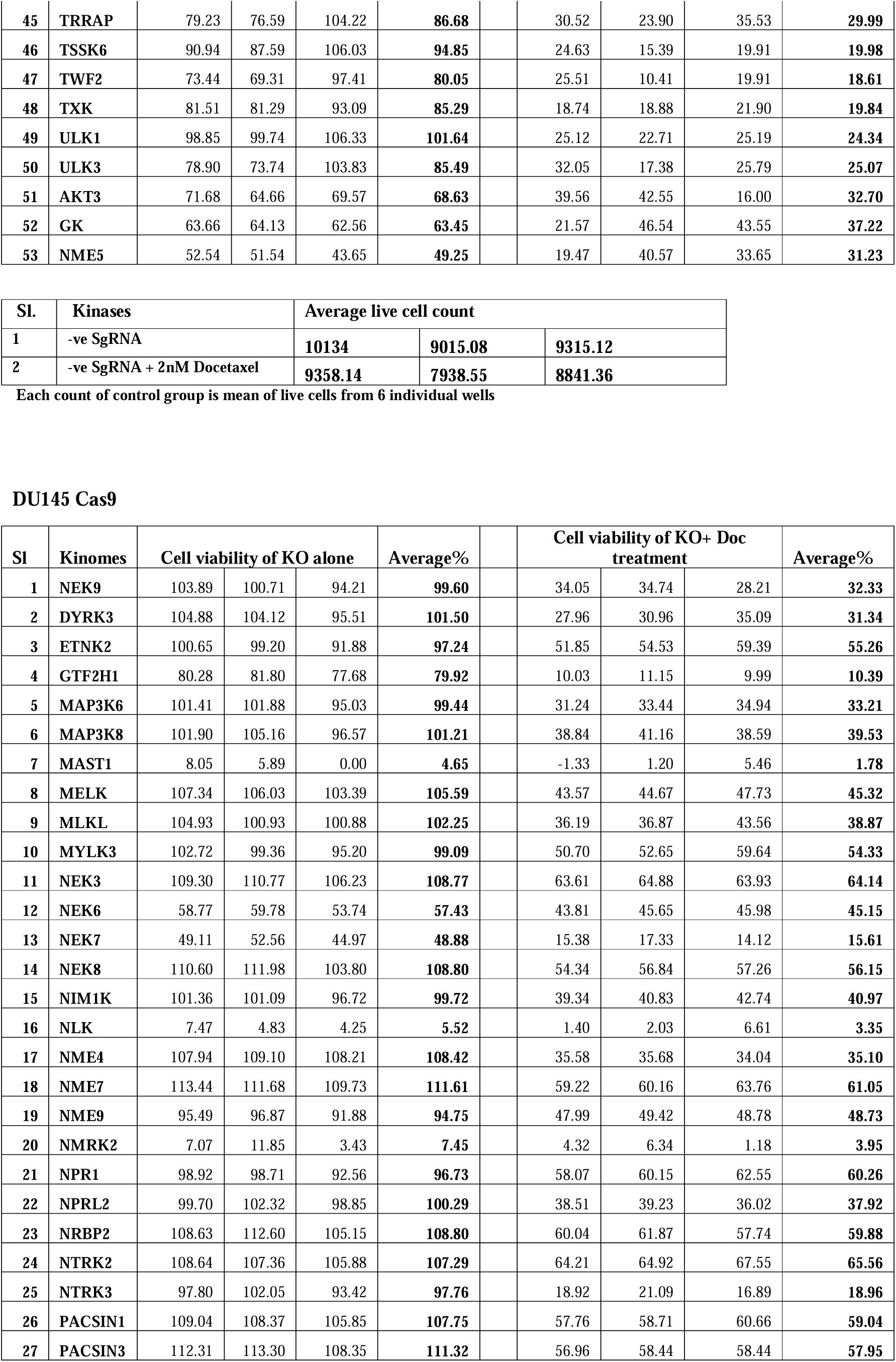
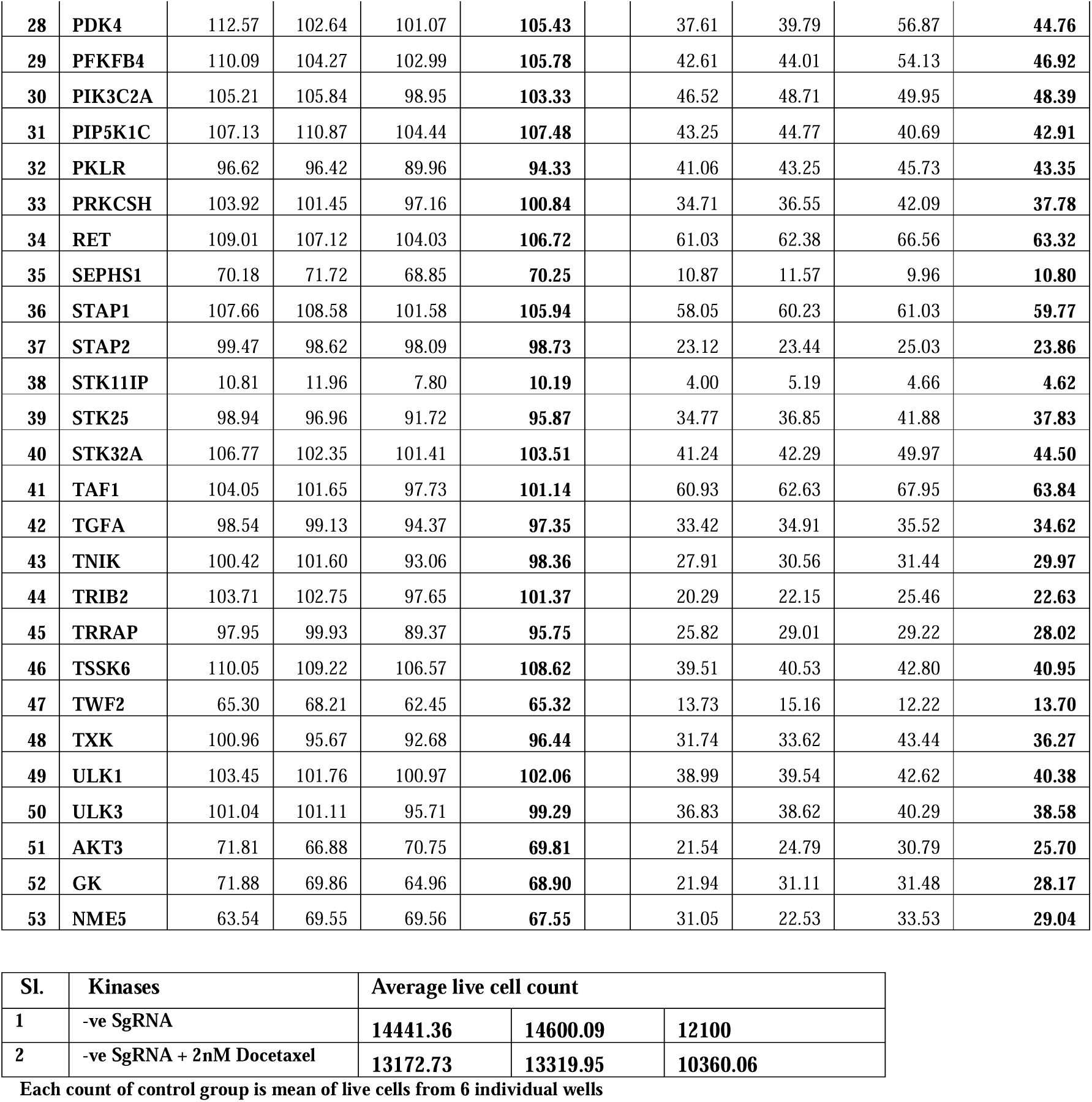
Secondary screening score from high content analyser system: % Cell viability of selected 50 targets from primary screening in multiple cell lines in absence and presence of Docetaxel. This is in reference to figure 1D.

**SCC4 Cas9**
| Sl | Kinomes | Cell viability of KO alone |  |  | Average% | Cell viability of KO+ Doc treatment |  |  | Average% |
| --- | --- | --- | --- | --- | --- | --- | --- | --- | --- |
| 1 | NEK9 | 74.61 | 71.75 | 89.91 | **78.76** | 21.71 | 19.19 | 6.10 | **15.67** |
| 2 | DYRK3 | 85.94 | 80.05 | 106.15 | **90.71** | 22.98 | 17.77 | 11.89 | **17.55** |
| 3 | ETNK2 | 70.01 | 65.41 | 88.76 | **74.73** | 17.87 | 11.80 | 46.79 | **25.49** |
| 4 | GTF2H1 | 69.29 | 64.99 | 100.90 | **78.39** | 32.11 | 22.40 | 31.44 | **28.65** |
| 5 | MAP3K6 | 93.46 | 88.03 | 111.62 | **97.71** | 19.91 | 10.03 | 28.18 | **19.37** |
| 6 | MAP3K8 | 111.82 | 103.19 | 141.92 | **118.98** | 24.75 | 16.32 | 32.48 | **24.52** |
| 7 | MAST1 | 84.22 | 79.57 | 137.29 | **100.36** | 23.17 | 16.84 | 31.37 | **23.80** |
| 8 | MELK | 73.74 | 72.10 | 100.73 | **82.19** | 20.90 | 17.23 | 35.95 | **24.70** |
| 9 | MLKL | 74.99 | 76.87 | 99.30 | **83.72** | 29.66 | 22.96 | 30.36 | **27.66** |
| 10 | MYLK3 | 77.60 | 75.04 | 69.44 | **74.03** | 27.98 | 20.68 | 23.75 | **24.14** |
| 11 | NEK3 | 80.02 | 75.59 | 106.38 | **87.33** | 29.34 | 24.67 | 31.42 | **28.48** |
| 12 | NEK6 | 70.93 | 65.52 | 49.87 | **62.11** | 18.04 | 11.82 | 32.03 | **20.63** |
| 13 | NEK7 | 82.81 | 72.37 | 97.15 | **84.11** | 23.71 | 10.82 | 18.72 | **17.75** |
| 14 | NEK8 | 72.28 | 60.54 | 90.91 | **74.58** | 32.45 | 20.14 | 30.72 | **27.77** |
| 15 | NIM1K | 99.47 | 87.54 | 112.84 | **99.95** | 31.26 | 16.19 | 40.72 | **29.39** |
| 16 | NLK | 84.14 | 72.34 | 121.59 | **92.69** | 29.15 | 11.64 | 26.21 | **22.33** |
| 17 | NME4 | 75.69 | 72.55 | 54.80 | **67.68** | 31.40 | 24.53 | 37.83 | **31.25** |
| 18 | NME7 | 74.35 | 65.96 | 52.86 | **64.39** | 29.90 | 14.81 | 30.37 | **25.03** |
| 19 | NME9 | 74.11 | 67.84 | 71.74 | **71.23** | 30.84 | 25.76 | 37.30 | **31.30** |
| 20 | NMRK2 | 101.72 | 92.41 | 117.11 | **103.74** | 28.09 | 18.29 | 57.40 | **34.59** |
| 21 | NPR1 | 75.34 | 70.86 | 107.57 | **84.59** | 16.22 | 8.11 | 17.65 | **13.99** |
| 22 | NPRL2 | 71.34 | 66.75 | 97.79 | **78.63** | 20.79 | 14.77 | 36.43 | **24.00** |
| 23 | NRBP2 | 79.13 | 70.71 | 103.07 | **84.30** | 30.67 | 27.20 | 36.21 | **31.36** |
| 24 | NTRK2 | 79.43 | 78.47 | 110.17 | **89.36** | 25.18 | 18.61 | 43.04 | **28.94** |
| 25 | NTRK3 | 74.39 | 64.97 | 85.39 | **74.92** | 28.88 | 22.13 | 33.50 | **28.17** |
| 26 | PACSIN1 | 84.60 | 82.75 | 89.64 | **85.66** | 33.88 | 27.29 | 41.64 | **34.27** |
| 27 | PACSIN3 | 69.90 | 65.43 | 80.01 | **71.78** | 30.91 | 29.89 | 19.93 | **26.91** |
| 28 | PDK4 | 69.82 | 58.88 | 108.88 | **79.19** | 39.31 | 10.51 | 35.95 | **28.59** |
| 29 | PFKFB4 | 74.69 | 67.61 | 87.98 | **76.76** | 29.88 | 12.46 | 42.87 | **28.40** |
| 30 | PIK3C2A | 76.26 | 63.17 | 104.51 | **81.32** | 32.27 | 15.91 | 33.76 | **27.32** |

|  |  |  |  |  |  |  |  |  |  |  |
| --- | --- | --- | --- | --- | --- | --- | --- | --- | --- | --- |
| 31 | PIP5K1C | 72.43 | 64.55 | 97.60 | 78.19 |  | 34.98 | 28.07 | 41.38 | 34.81 |
| 32 | PKLR | 90.25 | 84.00 | 93.47 | 89.24 |  | 36.73 | 24.57 | 21.35 | 27.55 |
| 33 | PRKCSH | 84.86 | 81.57 | 91.09 | 85.84 |  | 24.03 | 16.71 | 15.32 | 18.69 |
| 34 | RET | 101.86 | 92.00 | 114.15 | 102.67 |  | 39.99 | 22.06 | 20.44 | 27.50 |
| 35 | SEPHS1 | 81.16 | 73.11 | 89.29 | 81.19 |  | 37.22 | 21.54 | 34.91 | 31.23 |
| 36 | STAP1 | 80.73 | 74.69 | 100.88 | 85.43 |  | 34.95 | 33.26 | 33.78 | 34.00 |
| 37 | STAP2 | 101.26 | 100.51 | 115.21 | 105.66 |  | 33.02 | 22.58 | 43.49 | 33.03 |
| 38 | STK11IP | 112.22 | 107.70 | 120.47 | 113.46 |  | 31.95 | 28.92 | 34.85 | 31.90 |
| 39 | STK25 | 75.59 | 72.41 | 55.58 | 67.86 |  | 15.20 | 9.47 | 1.93 | 8.87 |
| 40 | STK32A | 76.41 | 71.66 | 70.68 | 72.91 |  | 35.48 | 16.61 | 46.41 | 32.84 |
| 41 | TAF1 | 79.72 | 76.66 | 114.19 | 90.19 |  | 32.00 | 27.68 | 32.27 | 30.65 |
| 42 | TGFA | 76.14 | 71.06 | 79.66 | 75.62 |  | 34.49 | 31.56 | 37.72 | 34.59 |
| 43 | TNIK | 94.26 | 84.70 | 97.81 | 92.26 |  | 26.07 | 13.75 | 47.89 | 29.24 |
| 44 | TRIB2 | 107.31 | 101.55 | 96.47 | 101.77 |  | 34.24 | 27.06 | 36.24 | 32.51 |
| 45 | TRRAP | 81.47 | 71.38 | 105.37 | 86.08 |  | 26.53 | 22.72 | 31.32 | 26.86 |
| 46 | TSSK6 | 122.55 | 115.29 | 104.20 | 114.01 |  | 24.98 | 13.87 | 15.31 | 18.05 |
| 47 | TWF2 | 81.90 | 71.16 | 103.52 | 85.53 |  | 21.97 | 10.22 | 69.48 | 33.89 |
| 48 | TXK | 72.31 | 68.31 | 81.07 | 73.90 |  | 24.86 | 8.52 | 34.22 | 22.54 |
| 49 | ULK1 | 78.01 | 76.10 | 107.11 | 87.07 |  | 23.95 | 15.93 | 32.44 | 24.11 |
| 50 | ULK3 | 67.96 | 56.22 | 124.16 | 82.78 |  | 36.91 | 15.54 | 34.88 | 29.11 |
| 51 | AKT3 | 66.56 | 70.02 | 61.25 | 65.95 |  | 23.45 | 29.54 | 31.12 | 28.04 |
| 52 | GK | 65.15 | 62.64 | 79.77 | 69.19 |  | 34.75 | 32.78 | 28.85 | 32.13 |
| 53 | NME5 | 63.48 | 61.54 | 78.56 | 67.86 |  | 22.79 | 29.77 | 29.45 | 27.33 |

| Sl. | Kinases | Average live cell count |  |  |
| --- | --- | --- | --- | --- |
| 1 | -ve SgRNA | 2469.12 | 2937.10 | 1842 |
| 2 | -ve SgRNA + 2nM Docetaxel | 1829.03 | 2254.78 | 1390.90 |

SCC9 Cas9
| Sl | Kinomes | Cell viability of KO alone |  |  | Average% | Cell viability of KO+ Doc treatment |  |  | Average% |
| --- | --- | --- | --- | --- | --- | --- | --- | --- | --- |
| 1 | NEK9 | 97.74 | 109.88 | 94.85 | 100.82 | 27.20 | 29.54 | 33.45 | 30.06 |
| 2 | DYRK3 | 111.21 | 105.32 | 77.45 | 97.99 | 13.35 | 17.54 | 27.04 | 19.31 |
| 3 | ETNK2 | 61.95 | 57.36 | 37.11 | 52.14 | 11.84 | 16.78 | 24.61 | 17.74 |
| 4 | GTF2H1 | 52.74 | 48.44 | 37.36 | 46.18 | 30.14 | 33.49 | 31.51 | 31.71 |
| 5 | MAP3K6 | 102.97 | 97.54 | 79.34 | 93.28 | 44.25 | 49.35 | 51.74 | 48.44 |
| 6 | MAP3K8 | 103.92 | 95.29 | 70.88 | 90.03 | 83.84 | 89.98 | 87.11 | 86.97 |
| 7 | MAST1 | 83.46 | 78.81 | 47.00 | 69.76 | 56.73 | 56.61 | 70.81 | 61.38 |
| 8 | MELK | 74.11 | 72.46 | 58.17 | 68.25 | 26.42 | 25.71 | 33.85 | 28.66 |
| 9 | MLKL | 99.64 | 101.51 | 90.41 | 97.18 | 20.66 | 16.38 | 31.83 | 22.96 |
| 10 | MYLK3 | 107.21 | 104.64 | 78.03 | 96.63 | 40.27 | 38.01 | 55.16 | 44.48 |
| 11 | NEK3 | 95.89 | 91.46 | 85.40 | 90.92 | 34.52 | 40.49 | 35.45 | 36.82 |
| 12 | NEK6 | 105.45 | 100.04 | 67.63 | 91.04 | 36.33 | 34.54 | 44.59 | 38.49 |

|  |  |  |  |  |  |  |  |  |  |  |
| --- | --- | --- | --- | --- | --- | --- | --- | --- | --- | --- |
| 13 | NEK7 | 108.31 | 97.88 | 83.16 | 96.45 |  | 70.28 | 80.46 | 68.24 | 72.99 |
| 14 | NEK8 | 103.81 | 92.08 | 71.51 | 89.13 |  | 40.90 | 51.33 | 43.42 | 45.22 |
| 15 | NIM1K | 113.66 | 101.74 | 82.92 | 99.44 |  | 73.60 | 80.96 | 73.23 | 75.93 |
| 16 | NLK | 107.05 | 95.25 | 76.99 | 93.10 |  | 28.01 | 31.90 | 26.59 | 28.83 |
| 17 | NME4 | 96.86 | 93.72 | 82.43 | 91.00 |  | 29.19 | 27.79 | 27.78 | 28.25 |
| 18 | NME7 | 105.09 | 96.70 | 75.36 | 92.39 |  | 67.66 | 68.88 | 70.28 | 68.94 |
| 19 | NME9 | 88.19 | 81.91 | 68.66 | 79.59 |  | 33.94 | 39.09 | 35.44 | 36.16 |
| 20 | NMRK2 | 61.76 | 52.46 | 40.69 | 51.64 |  | 8.18 | 19.11 | 5.36 | 10.88 |
| 21 | NPR1 | 96.74 | 92.26 | 90.59 | 93.20 |  | 34.93 | 45.30 | 40.22 | 40.15 |
| 22 | NPRL2 | 96.06 | 91.46 | 83.03 | 90.18 |  | 46.06 | 49.48 | 43.76 | 46.43 |
| 23 | NRBP2 | 108.64 | 100.22 | 85.15 | 98.01 |  | 68.86 | 76.93 | 67.59 | 71.12 |
| 24 | NTRK2 | 108.13 | 107.17 | 98.10 | 104.47 |  | 82.91 | 82.34 | 88.25 | 84.50 |
| 25 | NTRK3 | 102.92 | 93.49 | 68.42 | 88.28 |  | 61.06 | 67.22 | 62.03 | 63.44 |
| 26 | PACSIN1 | 74.73 | 72.88 | 56.67 | 68.09 |  | 15.23 | 13.53 | 21.53 | 16.76 |
| 27 | PACSIN3 | 109.06 | 104.59 | 91.48 | 101.71 |  | 62.31 | 66.19 | 66.06 | 64.86 |
| 28 | PDK4 | 115.63 | 104.69 | 78.62 | 99.65 |  | 20.22 | 24.40 | 35.67 | 26.76 |
| 29 | PFKFB4 | 96.32 | 89.23 | 71.26 | 85.60 |  | 8.97 | 11.31 | 18.45 | 12.91 |
| 30 | PIK3C2A | 116.89 | 103.80 | 74.60 | 98.43 |  | 84.19 | 91.00 | 86.76 | 87.32 |
| 31 | PIP5K1C | 110.12 | 102.24 | 88.36 | 100.24 |  | 51.42 | 58.38 | 49.38 | 53.06 |
| 32 | PKLR | 61.08 | 54.83 | 45.72 | 53.87 |  | 28.86 | 38.06 | 34.46 | 33.80 |
| 33 | PRKCSH | 113.61 | 110.32 | 89.41 | 104.45 |  | 89.39 | 89.80 | 101.06 | 93.41 |
| 34 | RET | 119.17 | 109.30 | 98.61 | 109.03 |  | 55.82 | 63.89 | 56.80 | 58.84 |
| 35 | SEPHS1 | 102.19 | 94.14 | 84.99 | 93.78 |  | 85.18 | 90.78 | 81.01 | 85.66 |
| 36 | STAP1 | 98.29 | 92.25 | 85.10 | 91.88 |  | 79.92 | 89.91 | 84.05 | 84.63 |
| 37 | STAP2 | 95.82 | 95.08 | 93.19 | 94.70 |  | 38.21 | 39.02 | 40.18 | 39.14 |
| 38 | STK11IP | 33.67 | 29.14 | 23.47 | 28.76 |  | 3.01 | 8.89 | 3.82 | 5.24 |
| 39 | STK25 | 57.66 | 54.48 | 48.26 | 53.47 |  | 21.39 | 28.53 | 30.67 | 26.86 |
| 40 | STK32A | 109.12 | 104.37 | 98.97 | 104.15 |  | 12.64 | 17.18 | 18.72 | 16.18 |
| 41 | TAF1 | 56.84 | 53.78 | 49.65 | 53.42 |  | 8.09 | 14.55 | 15.97 | 12.87 |
| 42 | TGFA | 64.60 | 59.51 | 49.80 | 57.97 |  | 58.34 | 64.01 | 61.32 | 61.22 |
| 43 | TNIK | 61.95 | 52.39 | 34.79 | 49.71 |  | 24.67 | 35.06 | 29.41 | 29.71 |
| 44 | TRIB2 | 101.64 | 95.88 | 76.32 | 91.28 |  | 69.26 | 72.55 | 76.86 | 72.89 |
| 45 | TRRAP | 112.39 | 102.30 | 72.45 | 95.72 |  | 52.50 | 60.51 | 60.60 | 57.87 |
| 46 | TSSK6 | 102.01 | 94.75 | 84.83 | 93.86 |  | 31.34 | 36.70 | 32.00 | 33.35 |
| 47 | TWF2 | 71.27 | 60.53 | 37.38 | 56.39 |  | 43.84 | 48.93 | 41.62 | 44.80 |
| 48 | TXK | 66.82 | 62.81 | 55.76 | 61.79 |  | 20.32 | 26.42 | 31.85 | 26.20 |
| 49 | ULK1 | 86.94 | 85.03 | 78.53 | 83.50 |  | 43.28 | 43.83 | 47.02 | 44.71 |
| 50 | ULK3 | 64.14 | 52.39 | 33.02 | 49.85 |  | 35.63 | 43.73 | 36.36 | 38.57 |
| 51 | AKT3 | 55.58 | 59.55 | 61.05 | 58.72 |  | 31.51 | 36.56 | 39.55 | 35.87 |
| 52 | GK | 75.54 | 63.57 | 66.54 | 68.55 |  | 39.56 | 27.56 | 41.48 | 36.20 |
| 53 | NME5 | 42.14 | 49.56 | 53.56 | 48.42 |  | 33.45 | 36.55 | 37.53 | 35.84 |

| Sl. | Kinases | Average live cell count |  |  |
| --- | --- | --- | --- | --- |
| 1 | -ve SgRNA | 2606.96 | 3122.75 | 2763.44 |
| 2 | -ve SgRNA + 2nM Docetaxel | 1899.04 | 2349.33 | 2143.12 |

MIA PaCA-2 Cas9
| Sl | Kinomes | Cell viability of KO alone |  |  | Average% |  | Cell viability of KO+ Doc treatment |  |  | Average% |
| --- | --- | --- | --- | --- | --- | --- | --- | --- | --- | --- |
| 1 | NEK9 | 89.20 | 95.72 | 110.82 | 98.58 |  | 15.73 | 16.65 | 27.14 | 19.84 |
| 2 | DYRK3 | 77.89 | 79.66 | 99.10 | 85.55 |  | 19.93 | 18.83 | 29.10 | 22.62 |
| 3 | ETNK2 | 95.85 | 96.81 | 111.55 | 101.40 |  | 14.62 | 15.76 | 23.27 | 17.89 |
| 4 | GTF2H1 | 81.88 | 80.10 | 91.25 | 84.41 |  | 8.95 | 5.46 | 9.87 | 8.09 |
| 5 | MAP3K6 | 78.24 | 77.33 | 92.73 | 82.77 |  | 13.86 | 12.18 | 19.15 | 15.06 |
| 6 | MAP3K8 | 89.21 | 86.20 | 109.09 | 94.83 |  | 27.39 | 20.29 | 29.86 | 25.85 |
| 7 | MAST1 | 91.71 | 96.49 | 115.21 | 101.14 |  | 23.15 | 20.39 | 31.77 | 25.10 |
| 8 | MELK | 81.17 | 84.12 | 92.01 | 85.77 |  | 15.31 | 14.35 | 19.39 | 16.35 |
| 9 | MLKL | 87.48 | 94.06 | 95.02 | 92.19 |  | 15.29 | 17.87 | 21.26 | 18.14 |
| 10 | MYLK3 | 84.68 | 91.15 | 105.31 | 93.71 |  | 12.80 | 11.02 | 20.32 | 14.71 |
| 11 | NEK3 | 91.34 | 87.67 | 97.29 | 92.10 |  | 16.66 | 15.08 | 17.84 | 16.53 |
| 12 | NEK6 | 89.46 | 93.41 | 113.58 | 98.82 |  | 24.13 | 16.92 | 28.63 | 23.23 |
| 13 | NEK7 | 77.11 | 69.65 | 93.49 | 80.08 |  | 16.43 | 7.04 | 13.69 | 12.39 |
| 14 | NEK8 | 93.98 | 87.87 | 116.96 | 99.60 |  | 20.23 | 9.48 | 18.29 | 16.00 |
| 15 | NIM1K | 74.58 | 69.55 | 100.30 | 81.48 |  | 30.59 | 14.11 | 22.37 | 22.36 |
| 16 | NLK | 92.02 | 88.96 | 121.30 | 100.76 |  | 32.64 | 11.12 | 19.17 | 20.98 |
| 17 | NME4 | 98.82 | 99.28 | 109.17 | 102.42 |  | 22.36 | 15.13 | 19.42 | 18.97 |
| 18 | NME7 | 90.82 | 90.88 | 116.11 | 99.27 |  | 32.42 | 17.59 | 26.10 | 25.37 |
| 19 | NME9 | 96.89 | 93.96 | 109.85 | 100.23 |  | 28.69 | 23.05 | 28.51 | 26.75 |
| 20 | NMRK2 | 86.85 | 78.62 | 98.31 | 87.93 |  | 22.57 | 16.53 | 22.01 | 20.37 |
| 21 | NPR1 | 106.25 | 101.09 | 109.38 | 105.57 |  | 14.97 | 18.39 | 19.69 | 17.68 |
| 22 | NPRL2 | 88.28 | 85.24 | 95.97 | 89.83 |  | 27.67 | 22.59 | 26.17 | 25.48 |
| 23 | NRBP2 | 83.61 | 77.81 | 97.28 | 86.23 |  | 23.03 | 16.29 | 22.71 | 20.68 |
| 24 | NTRK2 | 92.59 | 94.75 | 99.78 | 95.71 |  | 21.78 | 21.23 | 24.41 | 22.47 |
| 25 | NTRK3 | 93.65 | 89.87 | 114.35 | 99.29 |  | 26.09 | 16.99 | 26.92 | 23.33 |
| 26 | PACSIN1 | 99.49 | 102.73 | 111.53 | 104.58 |  | 21.41 | 19.00 | 24.71 | 21.71 |
| 27 | PACSIN3 | 103.98 | 102.64 | 114.71 | 107.11 |  | 15.58 | 13.11 | 18.22 | 15.64 |
| 28 | PDK4 | 90.21 | 92.57 | 127.75 | 103.51 |  | 19.21 | 3.18 | 13.69 | 12.03 |
| 29 | PFKFB4 | 77.55 | 79.15 | 102.00 | 86.23 |  | 34.91 | 24.34 | 31.51 | 30.25 |
| 30 | PIK3C2A | 89.27 | 85.59 | 121.18 | 98.68 |  | 32.59 | 15.45 | 27.37 | 25.14 |
| 31 | PIP5K1C | 103.00 | 97.68 | 115.98 | 105.55 |  | 18.65 | 11.41 | 17.36 | 15.81 |
| 32 | PKLR | 91.97 | 87.71 | 102.20 | 93.96 |  | 17.07 | 16.19 | 20.27 | 17.84 |
| 33 | PRKCSH | 90.43 | 94.05 | 107.54 | 97.34 |  | 21.96 | 20.21 | 27.73 | 23.30 |
| 34 | RET | 106.92 | 101.86 | 126.40 | 111.73 |  | 25.64 | 13.36 | 18.57 | 19.19 |
| 35 | SEPHS1 | 87.55 | 82.43 | 101.44 | 90.47 |  | 39.41 | 28.07 | 32.46 | 33.31 |
| 36 | STAP1 | 88.44 | 83.16 | 95.99 | 89.20 |  | 25.46 | 25.73 | 29.12 | 26.77 |
| 37 | STAP2 | 87.33 | 87.44 | 89.79 | 88.19 |  | 26.27 | 26.01 | 26.76 | 26.35 |
| 38 | STK11IP | 84.49 | 80.84 | 90.76 | 85.36 |  | 29.02 | 26.92 | 29.56 | 28.50 |
| 39 | STK25 | 93.64 | 91.97 | 99.84 | 95.15 |  | 10.17 | 13.95 | 16.55 | 13.56 |
| 40 | STK32A | 93.39 | 92.39 | 105.64 | 97.14 |  | 23.45 | 18.60 | 21.19 | 21.08 |
| 41 | TAF1 | 97.17 | 95.49 | 103.01 | 98.56 |  | 19.93 | 22.32 | 24.21 | 22.15 |
| 42 | TGFA | 90.28 | 87.50 | 99.97 | 92.58 |  | 16.36 | 13.68 | 17.76 | 15.93 |
| 43 | TNIK | 79.82 | 74.47 | 97.81 | 84.03 |  | 14.18 | 8.54 | 16.00 | 12.91 |
| 44 | TRIB2 | 88.96 | 89.30 | 106.92 | 95.06 |  | 18.67 | 13.80 | 21.28 | 17.92 |
| 45 | TRRAP | 79.23 | 76.59 | 104.22 | 86.68 |  | 30.52 | 23.90 | 35.53 | 29.99 |
| 46 | TSSK6 | 90.94 | 87.59 | 106.03 | 94.85 |  | 24.63 | 15.39 | 19.91 | 19.98 |
| 47 | TWF2 | 73.44 | 69.31 | 97.41 | 80.05 |  | 25.51 | 10.41 | 19.91 | 18.61 |
| 48 | TXK | 81.51 | 81.29 | 93.09 | 85.29 |  | 18.74 | 18.88 | 21.90 | 19.84 |
| 49 | ULK1 | 98.85 | 99.74 | 106.33 | 101.64 |  | 25.12 | 22.71 | 25.19 | 24.34 |
| 50 | ULK3 | 78.90 | 73.74 | 103.83 | 85.49 |  | 32.05 | 17.38 | 25.79 | 25.07 |
| 51 | AKT3 | 71.68 | 64.66 | 69.57 | 68.63 |  | 39.56 | 42.55 | 16.00 | 32.70 |
| 52 | GK | 63.66 | 64.13 | 62.56 | 63.45 |  | 21.57 | 46.54 | 43.55 | 37.22 |
| 53 | NME5 | 52.54 | 51.54 | 43.65 | 49.25 |  | 19.47 | 40.57 | 33.65 | 31.23 |

| Sl. | Kinases | Average live cell count |  |  |
| --- | --- | --- | --- | --- |
| 1 | -ve SgRNA | 10134 | 9015.08 | 9315.12 |
| 2 | -ve SgRNA + 2nM Docetaxel | 9358.14 | 7938.55 | 8841.36 |

DU145 Cas9
| Sl | Kinomes | Cell viability of KO alone |  |  | Average% |  | Cell viability of KO+ Doc treatment |  |  | Average% |
| --- | --- | --- | --- | --- | --- | --- | --- | --- | --- | --- |
| 1 | NEK9 | 103.89 | 100.71 | 94.21 | 99.60 |  | 34.05 | 34.74 | 28.21 | 32.33 |
| 2 | DYRK3 | 104.88 | 104.12 | 95.51 | 101.50 |  | 27.96 | 30.96 | 35.09 | 31.34 |
| 3 | ETNK2 | 100.65 | 99.20 | 91.88 | 97.24 |  | 51.85 | 54.53 | 59.39 | 55.26 |
| 4 | GTF2H1 | 80.28 | 81.80 | 77.68 | 79.92 |  | 10.03 | 11.15 | 9.99 | 10.39 |
| 5 | MAP3K6 | 101.41 | 101.88 | 95.03 | 99.44 |  | 31.24 | 33.44 | 34.94 | 33.21 |
| 6 | MAP3K8 | 101.90 | 105.16 | 96.57 | 101.21 |  | 38.84 | 41.16 | 38.59 | 39.53 |
| 7 | MAST1 | 8.05 | 5.89 | 0.00 | 4.65 |  | -1.33 | 1.20 | 5.46 | 1.78 |
| 8 | MELK | 107.34 | 106.03 | 103.39 | 105.59 |  | 43.57 | 44.67 | 47.73 | 45.32 |
| 9 | MLKL | 104.93 | 100.93 | 100.88 | 102.25 |  | 36.19 | 36.87 | 43.56 | 38.87 |
| 10 | MYLK3 | 102.72 | 99.36 | 95.20 | 99.09 |  | 50.70 | 52.65 | 59.64 | 54.33 |
| 11 | NEK3 | 109.30 | 110.77 | 106.23 | 108.77 |  | 63.61 | 64.88 | 63.93 | 64.14 |
| 12 | NEK6 | 58.77 | 59.78 | 53.74 | 57.43 |  | 43.81 | 45.65 | 45.98 | 45.15 |
| 13 | NEK7 | 49.11 | 52.56 | 44.97 | 48.88 |  | 15.38 | 17.33 | 14.12 | 15.61 |
| 14 | NEK8 | 110.60 | 111.98 | 103.80 | 108.80 |  | 54.34 | 56.84 | 57.26 | 56.15 |
| 15 | NIM1K | 101.36 | 101.09 | 96.72 | 99.72 |  | 39.34 | 40.83 | 42.74 | 40.97 |
| 16 | NLK | 7.47 | 4.83 | 4.25 | 5.52 |  | 1.40 | 2.03 | 6.61 | 3.35 |
| 17 | NME4 | 107.94 | 109.10 | 108.21 | 108.42 |  | 35.58 | 35.68 | 34.04 | 35.10 |
| 18 | NME7 | 113.44 | 111.68 | 109.73 | 111.61 |  | 59.22 | 60.16 | 63.76 | 61.05 |
| 19 | NME9 | 95.49 | 96.87 | 91.88 | 94.75 |  | 47.99 | 49.42 | 48.78 | 48.73 |
| 20 | NMRK2 | 7.07 | 11.85 | 3.43 | 7.45 |  | 4.32 | 6.34 | 1.18 | 3.95 |
| 21 | NPR1 | 98.92 | 98.71 | 92.56 | 96.73 |  | 58.07 | 60.15 | 62.55 | 60.26 |
| 22 | NPRL2 | 99.70 | 102.32 | 98.85 | 100.29 |  | 38.51 | 39.23 | 36.02 | 37.92 |
| 23 | NRBP2 | 108.63 | 112.60 | 105.15 | 108.80 |  | 60.04 | 61.87 | 57.74 | 59.88 |
| 24 | NTRK2 | 108.64 | 107.36 | 105.88 | 107.29 |  | 64.21 | 64.92 | 67.55 | 65.56 |
| 25 | NTRK3 | 97.80 | 102.05 | 93.42 | 97.76 |  | 18.92 | 21.09 | 16.89 | 18.96 |
| 26 | PACSIN1 | 109.04 | 108.37 | 105.85 | 107.75 |  | 57.76 | 58.71 | 60.66 | 59.04 |
| 27 | PACSIN3 | 112.31 | 113.30 | 108.35 | 111.32 |  | 56.96 | 58.44 | 58.44 | 57.95 |
| 28 | PDK4 | 112.57 | 102.64 | 101.07 | 105.43 |  | 37.61 | 39.79 | 56.87 | 44.76 |
| 29 | PFKFB4 | 110.09 | 104.27 | 102.99 | 105.78 |  | 42.61 | 44.01 | 54.13 | 46.92 |
| 30 | PIK3C2A | 105.21 | 105.84 | 98.95 | 103.33 |  | 46.52 | 48.71 | 49.95 | 48.39 |
| 31 | PIP5K1C | 107.13 | 110.87 | 104.44 | 107.48 |  | 43.25 | 44.77 | 40.69 | 42.91 |
| 32 | PKLR | 96.62 | 96.42 | 89.96 | 94.33 |  | 41.06 | 43.25 | 45.73 | 43.35 |
| 33 | PRKCSH | 103.92 | 101.45 | 97.16 | 100.84 |  | 34.71 | 36.55 | 42.09 | 37.78 |
| 34 | RET | 109.01 | 107.12 | 104.03 | 106.72 |  | 61.03 | 62.38 | 66.56 | 63.32 |
| 35 | SEPHS1 | 70.18 | 71.72 | 68.85 | 70.25 |  | 10.87 | 11.57 | 9.96 | 10.80 |
| 36 | STAP1 | 107.66 | 108.58 | 101.58 | 105.94 |  | 58.05 | 60.23 | 61.03 | 59.77 |
| 37 | STAP2 | 99.47 | 98.62 | 98.09 | 98.73 |  | 23.12 | 23.44 | 25.03 | 23.86 |
| 38 | STK11IP | 10.81 | 11.96 | 7.80 | 10.19 |  | 4.00 | 5.19 | 4.66 | 4.62 |
| 39 | STK25 | 98.94 | 96.96 | 91.72 | 95.87 |  | 34.77 | 36.85 | 41.88 | 37.83 |
| 40 | STK32A | 106.77 | 102.35 | 101.41 | 103.51 |  | 41.24 | 42.29 | 49.97 | 44.50 |
| 41 | TAF1 | 104.05 | 101.65 | 97.73 | 101.14 |  | 60.93 | 62.63 | 67.95 | 63.84 |
| 42 | TGFA | 98.54 | 99.13 | 94.37 | 97.35 |  | 33.42 | 34.91 | 35.52 | 34.62 |
| 43 | TNIK | 100.42 | 101.60 | 93.06 | 98.36 |  | 27.91 | 30.56 | 31.44 | 29.97 |
| 44 | TRIB2 | 103.71 | 102.75 | 97.65 | 101.37 |  | 20.29 | 22.15 | 25.46 | 22.63 |
| 45 | TRRAP | 97.95 | 99.93 | 89.37 | 95.75 |  | 25.82 | 29.01 | 29.22 | 28.02 |
| 46 | TSSK6 | 110.05 | 109.22 | 106.57 | 108.62 |  | 39.51 | 40.53 | 42.80 | 40.95 |
| 47 | TWF2 | 65.30 | 68.21 | 62.45 | 65.32 |  | 13.73 | 15.16 | 12.22 | 13.70 |
| 48 | TXK | 100.96 | 95.67 | 92.68 | 96.44 |  | 31.74 | 33.62 | 43.44 | 36.27 |
| 49 | ULK1 | 103.45 | 101.76 | 100.97 | 102.06 |  | 38.99 | 39.54 | 42.62 | 40.38 |
| 50 | ULK3 | 101.04 | 101.11 | 95.71 | 99.29 |  | 36.83 | 38.62 | 40.29 | 38.58 |
| 51 | AKT3 | 71.81 | 66.88 | 70.75 | 69.81 |  | 21.54 | 24.79 | 30.79 | 25.70 |
| 52 | GK | 71.88 | 69.86 | 64.96 | 68.90 |  | 21.94 | 31.11 | 31.48 | 28.17 |
| 53 | NME5 | 63.54 | 69.55 | 69.56 | 67.55 |  | 31.05 | 22.53 | 33.53 | 29.04 |

| Sl. | Kinases | Average live cell count |  |  |
| --- | --- | --- | --- | --- |
| 1 | -ve SgRNA | 14441.36 | 14600.09 | 12100 |
| 2 | -ve SgRNA + 2nM Docetaxel | 13172.73 | 13319.95 | 10360.06 |

Each count of control group is mean of live cells from 6 individual wells

### 4.2 In vitro NEK9 kinase activity assays

The *in-vitro* kinase assay was performed using the NEK9 Kinase Enzyme System (Promega, Cat No# VA7516) and ADP-Glo™ Kinase Assay (Promega, Cat No# V9101). Compounds namely Dabrafenib (Adooq Bioscience, Cat No# A11381) and Fostamatinib (Adooq Bioscience, Cat No# A11926) (10 μM) were incubated with recombinant human NEK9 kinase, substrate MBP, and ATP mixed in 1X kinase buffer for 1 hour. ADP-Glo™ Reagent was added for 40 min to stop the reaction and deplete the unconsumed ATP, leaving only ADP. Kinase detection reagent was added and incubated for an additional 30 min to convert ADP to ATP and introduce luciferase and luciferin to detect the ATP. The luminescence was measured using VICTOR® Nivo™ Multimode Plate Reader (Perkin Elmer). All the reaction steps were carried out at room temperature.

### 4.3 Cell culture

OSCC cell lines were procured from a European collection of authenticated cell cultures (ECACC). The human tongue OSCC lines (H357, SCC4, and SCC9) and lung carcinoma epithelial cell line A549 were cultured and maintained in DMEM F12 (0.6 μg/ml sodium hydrocortisone succinate). PDC1 patient-derived cells were established from OSCC chemotherapy non-responder patient’s tumor tissue (treated with Docetaxel and Cisplatin without having any response) and were maintained in DMEM F12 (0.6 μg/ml sodium hydrocortisone succinate). HEK293T, MIA PaCa-2, DU145, MDAMB231, and MCF7 were maintained in DMEM media. Both the media were supplemented with 10% FBS (Thermo Fisher Scientific), and penicillin-streptomycin (Pan Biotech).

### 4.4 Generation of Docetaxel acquired resistant cell line

Docetaxel-acquired resistant cell lines were generated by consistent treatment of Docetaxel starting with a very low dose of 0.1nM and gradually rising to the IC50 values (1.5nM), over a longer period of time up to eight months. Parental cells were classified as sensitive namely H357 Doc S, and resistant ones as H357 Doc R. The generated resistant cells were maintained in media supplemented with minimal Docetaxel concentration (0.05nM).

### 4.5 Lentivirus production and generation of stable NEK9 and GSDME KO cell lines

LentiCas9-Blast was obtained from Addgene (Addgene, Cat No# 52962), which was generously donated by the Feng Zhang lab (Sanjana *et al*, 2014). NEK9 or GSDME targeting Sg RNAs were cloned into pKLV2-U6gRNA5(BbsI)-PGKpuro2ABFP-W (Addgene, Cat No# 67974) vector according to the methods described in Addgene, which was graciously donated by the Kosuke Yusa lab (Tzelepis *et al*, 2016). Lentiviruses were generated by transfecting HEK293T cells with the LentiCas9-Blast or pKLV2-U6gRNA5(BbsI)-PGKpuro2ABFP-W plasmids along with the packaging plasmid psPAX2 and the envelop plasmid pMD2G as described in Shriwas et al (Shriwas *et al*, 2021). Cas9 lentivirus and NEK9 Sg RNA lentivirus were then transduced in cancer cells using polybrene (8μg/ml), followed by puromycin (up to 5μg/ml) selection. After a week, cells were harvested and seeded in 96 well plates at a dilution of one cell per well, and individual colonies were picked up. Immunoblotting was performed to confirm the positive clones. The details of primer sequences are mentioned in Table 6.

### 4.6 Lentivirus production and generation of stable NEK9 KD cell lines

pLKO.1 vector was obtained from Addgene (Addgene, Cat No# 10878), deposited graciously by the David Root lab(Moffat *et al*, 2006). NEK9 Sh RNA was cloned into pLKO.1 according to the protocol specified in Addgene. The lentiviruses were produced as described by Shriwas et al (Shriwas *et al*., 2021), The NEK9 Sh RNA lentiviral particles were transduced in SCC9 and DU145 cells in the presence of polybrene (8µg/ml) followed by puromycin (up to 5µg/ml) selection for three weeks. To obtain the clones stably expressing NEK9 Sh RNA, distant colonies were picked up and NEK9 knockdown was confirmed by immunoblotting. The details of primer sequences are mentioned in Table 6.

### 4.7 Immunoblotting

Immunoblotting was performed by the method described in Maji et al (Maji *et al*, 2015). In this study, primary antibodies used were against β-actin (Sigma, Cat No# A2066), Cas9 (CST, Cat No# 14697S), NEK9(Abcam, Cat No# ab138488), PARP (CST, Cat No# 9542L), γH2AX (CST, Cat No# 9718S), cleaved caspase 3 (CST, Cat No# 9661S), GSDME (Abcam, Cat No# ab215191), GSDMD (CST, Cat No# 97558S), mature IL1β (Asp116) (CST, Cat No# 83186S), ERK1/2 (CST, Cat No# 4695), pERK1/2 T202/Y204 (CST, Cat No# 9101S), secondary mouse antibody (Novus, Cat No# NB7539), secondary rabbit antibody (Novus, Cat No# NB7160). The quantification of band intensity when required in immunoblots (n = 3) was performed using ImageJ software (NIH). The mean value of band intensity is indicated under the immunoblots. The intensity for each blot was calculated by normalizing with the intensity of the β-Actin blot (internal control).

### 4.8 Assessment of cell viability

Cell viability was measured by 3-(4, 5-dimethylthiazol-2-yl)-2, 5-diphenyltetrazolium bromide (MTT; Sigma-Aldrich) assay as per manufacturer’s instruction.

### 4.9 Colony formation assay

Colony formation assay was performed as described in Shriwas et al (Shriwas *et al*., 2021).

### 4.10 Annexin-V PE/7-AAD Assay

Apoptosis and cell death assay was performed by using Annexin V Apoptosis Detection Kit PE (eBioscience™, USA, Cat No# 88-8102-74). Cells were trypsinized and resuspended in a staining solution prepared in the binding buffer as mentioned in the manufacturer’s protocol, and cell death was monitored using a flow cytometer (BD FACS Fortessa, USA).

### 4.11 Immunofluorescence

The cells were seeded on a lysine-coated coverslip and cultured overnight. The next day, cells were treated with Docetaxel for 48 hr followed by 4% formaldehyde fixation for 15 min. Next, cells were permeabilized with 0.1% Triton X-100 prepared in PBS for 10-15 min, followed by blocking with 1% BSA for 30 min at room temperature. The cells were then incubated with primary antibody overnight at 4 °C. After PBS wash, cells were incubated with Goat anti–Rabbit IgG (H+L) secondary Antibody, Alexa Fluor® 488 conjugate (Invitrogen, Cat No# A −11008) for 1 hr at RT. After washing three times with PBS, cells were mounted in DAPI (Slow Fade ® GOLD Antifade, Thermo Fisher Scientific, Cat No# S36938). Images were captured using confocal microscopy (LEICA TCS-SP8). Anti β tubulin antibody (Sigma, Cat No# T4026) was used in this study.

### 4.12 Immunohistochemistry

Formalin-fixed paraffin embedded 4-5μm thick tumor sections from tumor xenografts were deparaffinized with xylene followed by rehydration, antigen unmasking, and quenching in 3% H_2_O_2_ for 20□min. After PBS wash, sections were again blocked with 2.5% horse serum for 1hr at RT. The tissues were incubated with the required dilution of primary antibodies overnight at 4°C. After PBS wash, sections were incubated with biotinylated antibody (Vector Laboratories, Cat No# 30082) for 45 min, followed by incubation with ABC reagent (Vector Laboratories, Cat No# PK-7200) for 30 min at RT. Immunoreactivity was determined using diaminobenzidine as the final chromogen. Lastly, sections were counterstained with Delafield’s hematoxylin (Himedia, Cat No# S014) and then mounted with DPX mounting medium. In this study, primary antibodies used were against NEK9 (Santa Cruz, Cat No# sc-100401), cleaved GSDME (Asp270) (CST, Cat No# 55879) and cleaved caspase-3 (CST, Cat No# 9661S). Images were obtained using a Leica DM500 microscope (Leica, Germany). Q-score was calculated by multiplying the percentage of positively stained cells and intensity of staining which was determined as strong (value=3), intermediate (value=2), weak (value=1), and negative (value=0) intensity.

### 4.13 Transient transfection and overexpression of NEK9 WT and NEK9 T210A genes in NEK9 KD cells

pMRX-IPU-FLAG-NEK9 (Addgene, Cat No# 168269) and pMRX-IPU-FLAG-NEK9T210A (Addgene, Cat No# 168270) plasmids were procured from Addgene those were graciously gifted by Noboru Mizushima (Yamamoto *et al*, 2021). SCC9 NEK9 KD and DU145 NEK9 KD cells were transiently transfected with NEK9 WT (pMRX-IPU-FLAG-NEK9) and NEK9 kinase mutant (pMRX-IPU-FLAG-NEK9T210A) plasmids using the ViaFect transfection reagent (Promega, Cat No# E4982).

### 4.14 NEK9 WT and NEK9 kinase mutant retrovirus generation

HEK293T cells were seeded the previous day in a 10cm culture plate. The next day, 70-80% confluent cells were transfected with pMRX-IPU-FLAG-NEK9 or pMRX-IPU-FLAG-NEK9T210A along with pCL-Eco plasmid (Addgene, Cat No# 12371) that was graciously donated by Inder Verma (Naviaux *et al*, 1996) and incubated in 32° incubator, 5% CO_2_ for 48 hr. The soup was collected and stored at −80° until further use.

### 4.15 Generation of stable H357 cell lines overexpressing NEK9 WT and NEK9 T210A kinase

H357 cells were transduced with NEK9 WT and NEK9 T210A retroviruses with polybrene (8g/ml), followed by puromycin (up to 5μg/ml) selection. Individual distant colonies were picked up and overexpression was confirmed by immunoblot analysis.

### 4.16 OSCC patient sample

Human OSCC samples were collected and categorized as responders and non-responders as described in Mohanty et al 2022(Mohanty *et al*., 2022). The Human Ethics Committee (HEC) of the Institution of Life Sciences approved all patient-related studies, and informed consent was obtained from all patients. Study subject details with treatment modalities are presented in Tables 4 and 5.

**Table 4.**
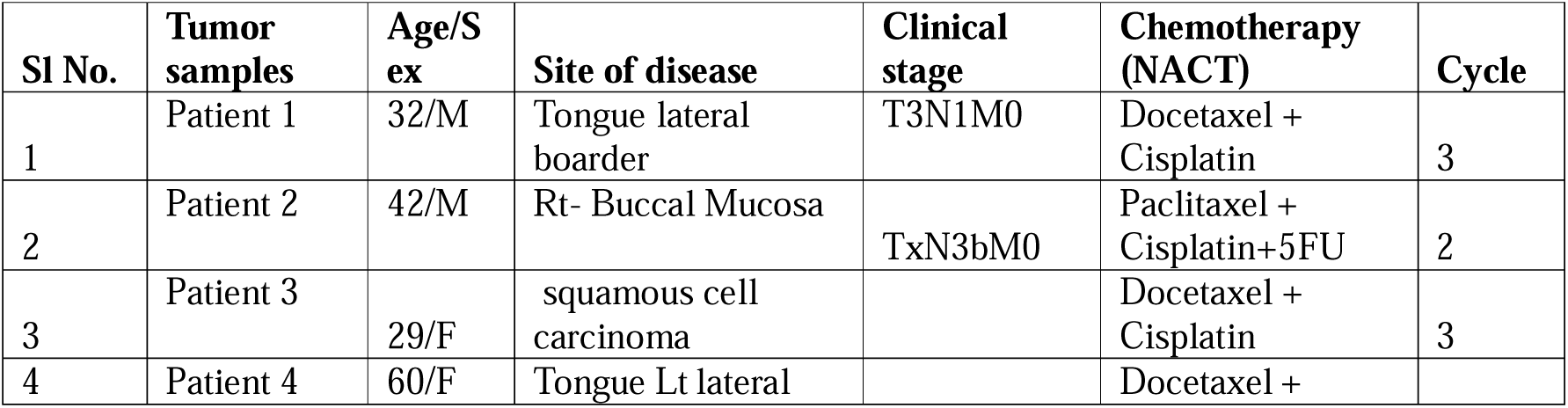
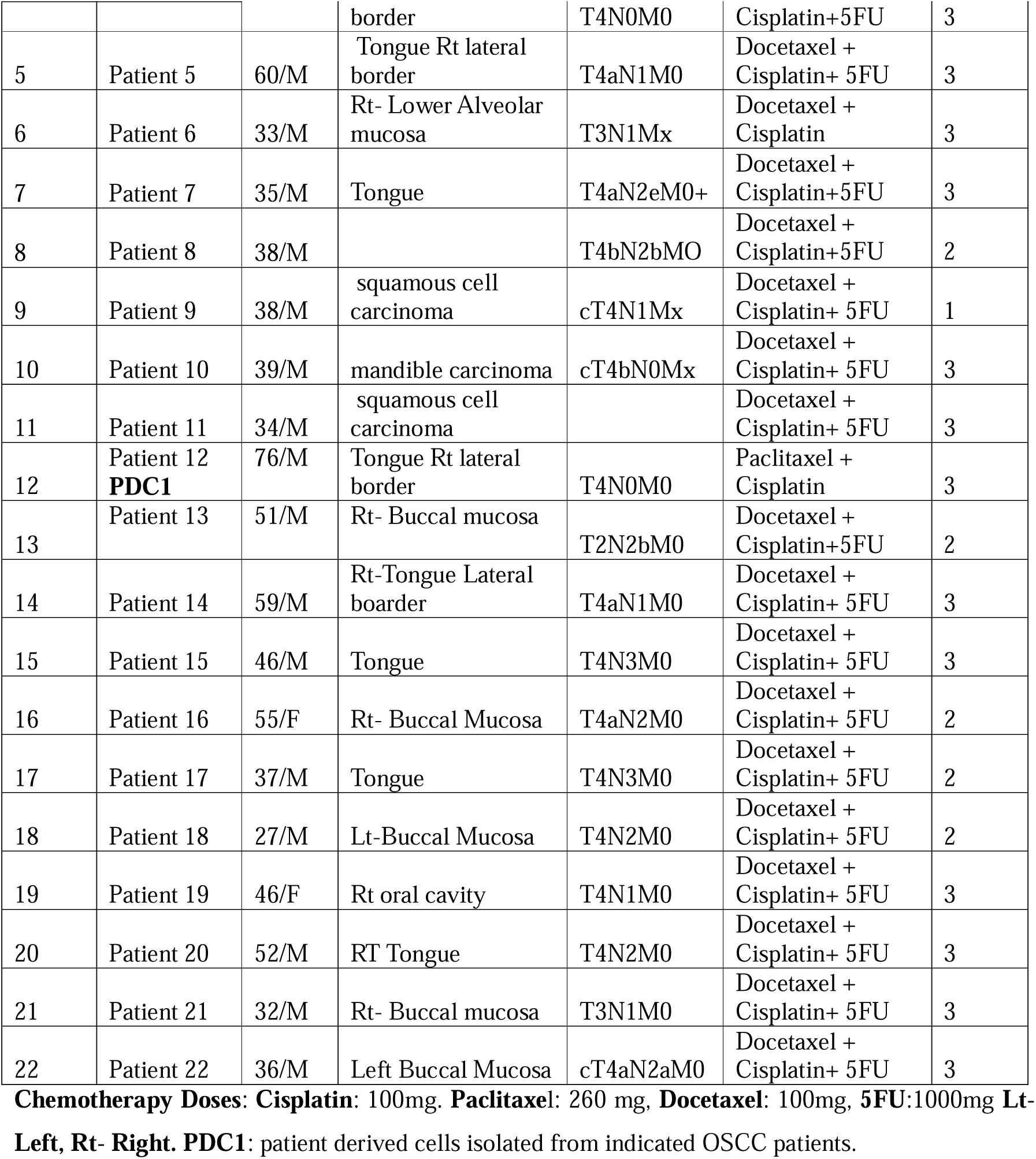
Chemotherapy non-responder patient details.

**Table 5.**
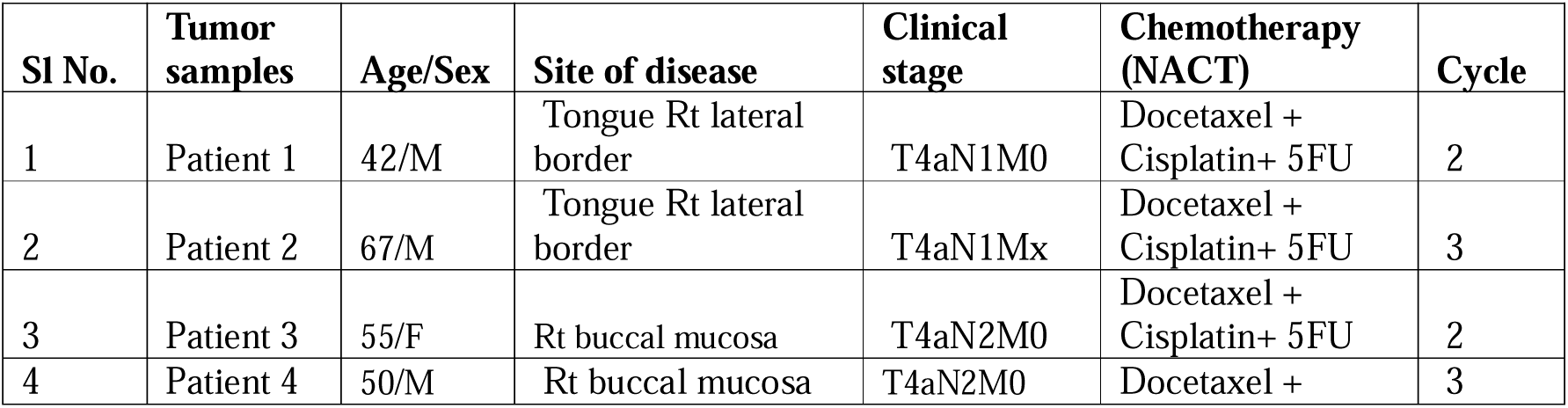
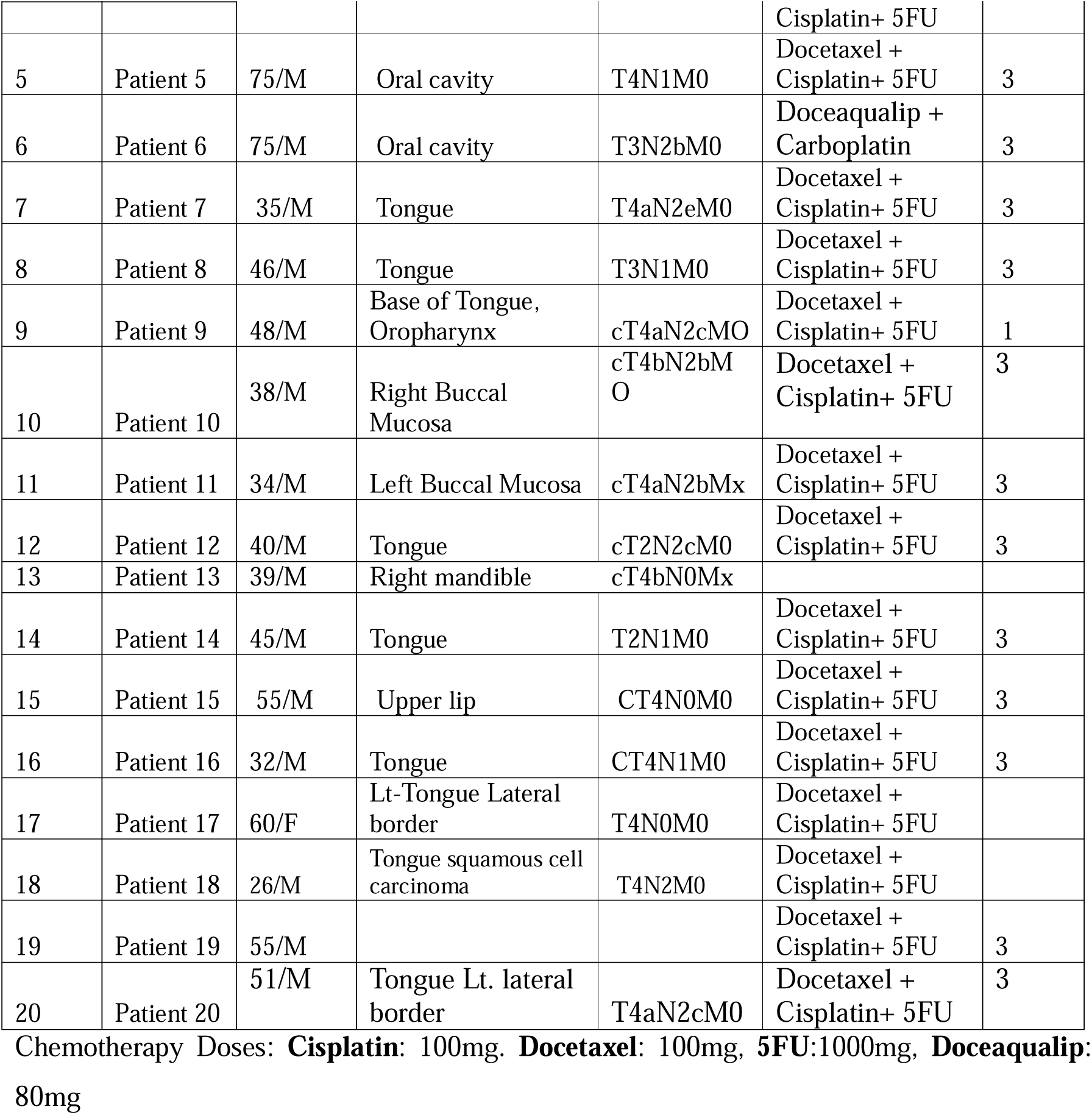
Chemotherapy responder patient details.

**Table 6.**
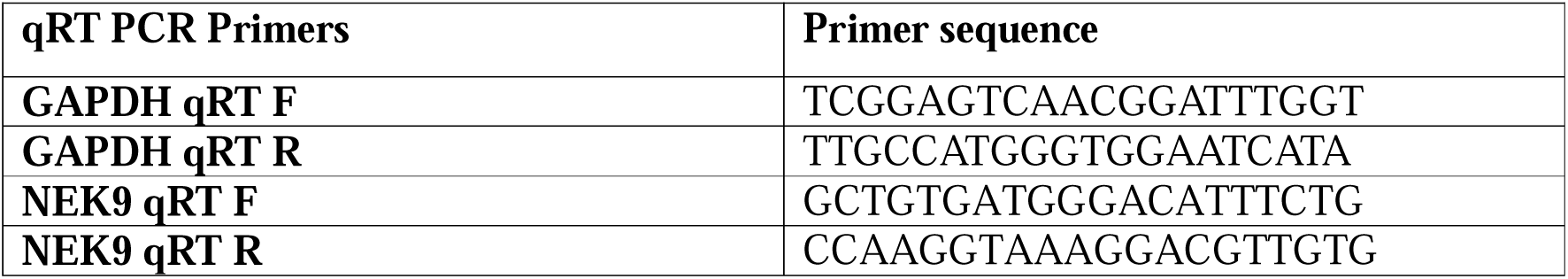
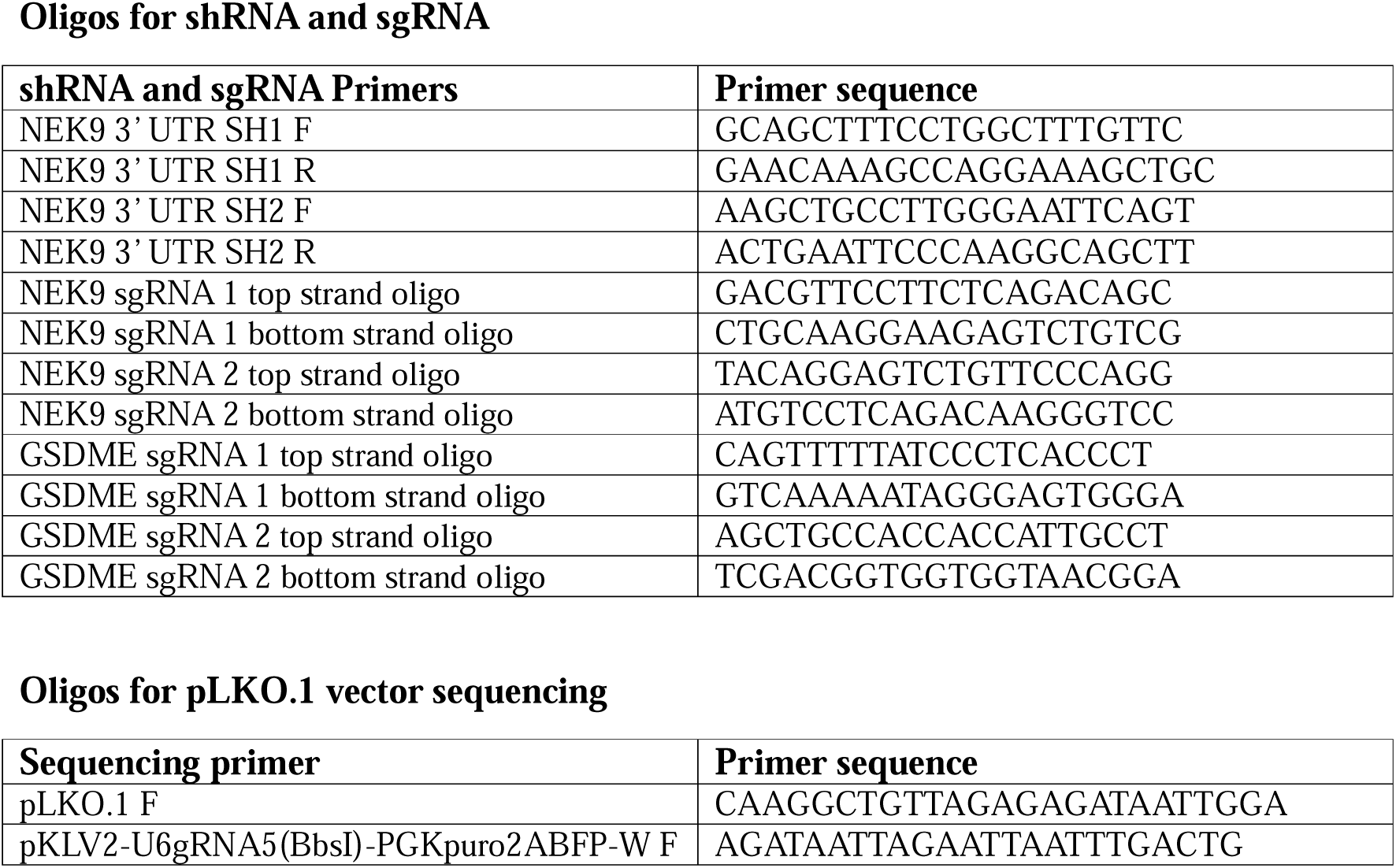
Oligos for qRT PCR.

### 4.17 Patient Derived Xenograft

BALB/C-nude mice (6-8 weeks, male, NCr-Foxn1nu athymic) were purchased from Vivo Bio-Tech. Ltd. For the xenograft model, the early passage of patient-derived cells (PDC1) established from chemo non-responder patients (treated with TP without having any response) was considered. Two million cells were suspended in phosphate-buffered solution-Matrigel (1:1, 100 μl) and transplanted into the upper flank of mice. The PDC1 NEK9 WT cells were injected in the right upper flank and PDC1 NEK9 KO cells were injected in the left upper flank of the same mice. These mice were randomly divided into 2 groups (n=5) once the tumors reached a volume of ∼50 mm^3^ and injected with vehicle control or Docetaxel (5mg/kg) intraperitoneally twice a week. In another experimental setup, PDC1 WT cells were injected into the right upper flank of mice. These mice were randomly divided into 4 groups (n=4) after the tumors had reached a volume of ∼50 mm^3^ and injected with vehicle control, Docetaxel (5mg/kg) alone, Fostamatinib (0.5mg/kg) alone or Docetaxel (5mg/kg) and Fostamatinib (0.5mg/kg) combined respectively in each group, intraperitoneally. Fostamatinib (0.5mg/kg) was injected on a one-day interval. Tumor size was measured using a digital Vernier caliper every two days until the completion of experiments. Tumor volume was determined using the following formula: Tumor volume (mm^3^) = (minimum diameter)^2^ × (maximum diameter).

### 4.18 Zebrafish xenograft

The experimental protocols used for this work were approved by the institutional animal ethical review committee (ILS/IAEC-214-AH/APR-21). The study involved the use of PDC1 control and NEK9 stable knockout cells. The cells were suspended in normal media after trypsinization and labeled using a cell-labeling solution (Vybrant™ DiI Cell-Labeling Solution, Thermo Fisher Scientific, Cat No# V22885). The stained cells were resuspended in warm media with a final density of 200 cells/nl. Approximately 400 cells were then microinjected (Femtojet microinjector) into perivitelline space of 48 hours post fertilization (hpf) embryos of zebrafish (*Danio rerio*) [Tg(fli1: EGFP)] for the development of tumor. The images of zebrafish embryos were captured using a fluorescence stereomicroscope (Leica MZ16) on the day of injection (Day 0), followed by Docetaxel treatment (5μM) 3 days post injection (Day 3) and then final imaging was obtained 5 days after injection (Day 5). The tumor growth was assessed by an increase or decrease in fluorescence intensity on the 5^th^ day compared to the day of injection. The fluorescence intensity was quantified using ImageJ software and represented as mean fluorescent intensity where day 0 readings were taken as baseline. Similar experiments were performed with PDC1 tumor-bearing fishes treated with Docetaxel (5μM) and Fostamatinib (10μM) or the combination of both for accessing the combinatorial effect of both drugs.

### 4.19 Chloroform-methanol protein precipitation

Proteins from cultured media supernatants were isolated by methanol/chloroform precipitation assay. Briefly, 500μl of supernatant, 500μl of methanol, and 125μl of chloroform were mixed together by for 30 seconds (vortex) to get a milky white solution.

After centrifugation, the aqueous phase was removed without disturbing the protein layer. An additional 500μl of methanol was added to the sample and vortexed well to break the protein layer followed by centrifugation. The protein pellet was air-dried and re-suspended in sample loading dye. The sample was boiled at 95°C for 5 min and separated on a 12% SDS-polyacrylamide gel and transferred to the PVDF membrane.

### 4.20 Soluble and Insoluble proteins fractionation

The insoluble pellet fraction after initial homogenization in lysis buffer was washed 3 times in PBS. The pellet was then dissolved in a buffer containing 2% Triton X-100, 50mM Tris HCl (pH 8), 150mM NaCl, 1mM EDTA, 10% glycerol, protease inhibitor, and 1mM PMSF. The dissolved contents were boiled at 95°C for 5-8 min and then allowed to cool at RT. After quantification, an equal concentration of proteins was boiled again with a sample loading buffer without a reducing agent. Proteins were resolved on 6% of SDS PAGE and immunoblotting was carried out as mentioned earlier with GSDME primary antibody.

### 4.21 LDH release assay

The release of LDH cancer cells was determined using the CyQUANT™ LDH Cytotoxicity Assay (Invitrogen, Cat No# C20300). Briefly, 50μl of each sample medium was mixed with 50μl of LDH reaction mixture (prepared as per manufacturer’s protocol) in 96 well plates and incubated for 30 min at RT. 50μl of stop solution was added to each sample and mixed well by gentle tapping followed by measuring absorbance at 490 nm/ 680 nm. To determine the LDH activity, the 680 nm absorbance value was subtracted from the 490 nm absorbance value.

### 4.22 RT-PCR and Real-Time Quantitative PCR

RNA mini kit (HiMedia, Cat No# MB602) was used to isolate total RNA as per the manufacturer’s instruction and quantified by Nanodrop. Verso cDNA synthesis kit (Thermo Fisher Scientific, Cat No# AB1453A) was used to synthesize cDNA by reverse transcription PCR using 300ng of RNA. qRT-PCR was carried out using SYBR Green master mix (Thermo Fisher Scientific, Cat No# 4367659). GAPDH was used as an internal control. The details of primer sequences are mentioned in Table 6.

### 4.23 Scanning Electron Microscopy (SEM)

NEK9 WT and KO cells were cultured on 10mm glass coverslips coated with 1% poly-l-lysine. The next day cells were treated with vehicle control and Docetaxel (2nM). After 36 hr of treatment, cells were fixed for 1–2 hr in 2% paraformaldehyde and 3% glutaraldehyde prepared in PBS buffer. Post fixation, cells were washed with PBS and then dehydrated through ethanol series (40%, 60%, 80%, 100%), dried at RT for 1 hr and then at 37°C for 1-2 days, mounted on stubs, sputter coated with a thin layer of conductive metal, gold and palladium, and viewed under SEM (JSM-IT800 system).

### 4.24 Alkaline comet assay

1.0 × 10^5^/ml trypsinized and PBS-washed cells were resuspended in 400μl of PBS. Slides were pre-coated with 1% of low melting agarose (Sigma, Cat No# A9414-25G) and air dried. The cell suspension was mixed with 1.2ml of 1% molten agarose (acclimatized at 37°C) and 1.2ml of the solution was spread over the precoated slide followed by covering with the glass coverslip. The slides were kept at 4°C for 10 min and the cover slips were slid and removed gently. All the procedures hereafter were done at 4°C. The prepared slides were dipped with chilled lysis buffer (2.5M NaCl, 100mM Na2EDTA, 10mM Tris-HCl, 200mM NaOH, pH 10) for 2hr. 1% of Trito X-100 and 1% of DMSO were added just prior to use in the lysis buffer. After lysis, slides were then dipped in chilled electrophoresis buffer (300mM NaOH, 1mM Na2EDTA, 1% DMSO, pH>13) for 20 min, and then the gel was run at 300mAmp for 45 min in the chilled electrophoresis buffer. After the run, slides were again dipped in chilled neutralization buffer (500mM Tris-HCl, pH 7.5) for 20 min followed by staining with 2.5μg/ml Propadium iodide in water for 20 min and slides were accessed for COMET tail. The images were captured in an Apotome microscope (Zeiss) and tail lengths were analyzed by *CASP* Version *1.2*.*3b1* software (Sourceforge, Diceholdings Inc., New York, NY, USA).

### 4.25 Statistical analysis

The data are presented as mean and standard deviation using Graph Pad Prism 9.0, and statistical significance was determined using 2-tailed Student’s t-test, one-way variance (one-way ANOVA), and Two-Way ANOVA, with P ≤ 0.05.

### 4.26 Study approval

The study was approved by the Institute Review Board and Human Ethics Committees (HEC) of the Institute of Life Sciences, Bhubaneswar (120/HEC/23) and All India Institute of Medical Sciences, Bhubaneswar (T/EMF/ Surg.Onco/19/03). Animal-related experiments were conducted in accordance with the protocol approved by the Institutional Animal Ethics Committee of Institute of Life Sciences, Bhubaneswar (ILS/IAEC-289-AH/ NOV-22). The Institutional Biosafety Committee (IBSC) approved all related experiments.

## 5. Acknowledgements

SAA is a UGC-SRF, SM is UGC-SRF and PM is CSIR-SRF. We acknowledge Institute of Life Sciences, Bhubaneswar intramural support and DBT BT/ INF/22/SP28293/2018 (for imaging facility). This work is funded by DST-SERB (CRG/2022/003427).

## 6. Author contributions

RD: Project administration, Supervision, SAA, RD: Conceptualisation, SAA, RKS, SM, PM: Methodology, RR, DM, SKDM: Validation, Visualization, SAA, RD: Writing-original draft, Writing-review and editing, SAA, RD: Data curation, Formal analysis.

## 8. References

Belham C, Roig J, Caldwell JA, Aoyama Y, Kemp BE, Comb M, Avruch J (2003) A mitotic cascade of NIMA family kinases. Nercc1/Nek9 activates the Nek6 and Nek7 kinases. J Biol Chem 278: 34897–34909

Bertran MT, Sdelci S, Regue L, Avruch J, Caelles C, Roig J (2011) Nek9 is a Plk1-activated kinase that controls early centrosome separation through Nek6/7 and Eg5. EMBO J 30: 2634–2647

Braselmann S, Taylor V, Zhao H, Wang S, Sylvain C, Baluom M, Qu K, Herlaar E, Lau A, Young C et al (2006) R406, an orally available spleen tyrosine kinase inhibitor blocks fc receptor signaling and reduces immune complex-mediated inflammation. J Pharmacol Exp Ther 319: 998–1008

Bussel J, Arnold DM, Grossbard E, Mayer J, Trelinski J, Homenda W, Hellmann A, Windyga J, Sivcheva L, Khalafallah AA et al (2018) Fostamatinib for the treatment of adult persistent and chronic immune thrombocytopenia: Results of two phase 3, randomized, placebo-controlled trials. Am J Hematol 93: 921–930

Chao OS, Goodman OB, Jr. (2021) DNA-PKc inhibition overcomes taxane resistance by promoting taxane-induced DNA damage in prostate cancer cells. Prostate 81: 1032–1048

Clemens GR, Schroeder RE, Magness SH, Weaver EV, Lech JW, Taylor VC, Masuda ES, Baluom M, Grossbard EB (2009) Developmental toxicity associated with receptor tyrosine kinase Ret inhibition in reproductive toxicity testing. Birth Defects Res A Clin Mol Teratol 85: 130–136

Demarco B, Grayczyk JP, Bjanes E, Le Roy D, Tonnus W, Assenmacher CA, Radaelli E, Fettrelet T, Mack V, Linkermann A et al (2020) Caspase-8-dependent gasdermin D cleavage promotes antimicrobial defense but confers susceptibility to TNF-induced lethality. Sci Adv 6

Dumontet C, Jordan MA (2010) Microtubule-binding agents: a dynamic field of cancer therapeutics. Nat Rev Drug Discov 9: 790–803

Erkes DA, Cai W, Sanchez IM, Purwin TJ, Rogers C, Field CO, Berger AC, Hartsough EJ, Rodeck U, Alnemri ES et al (2020) Mutant BRAF and MEK Inhibitors Regulate the Tumor Immune Microenvironment via Pyroptosis. Cancer Discov 10: 254–269

Fan CY, Ye FH, Peng M, Dong JJ, Chai WW, Deng WJ, Zhang H, Yang LC (2023) Endogenous HMGB1 regulates GSDME-mediated pyroptosis via ROS/ERK1/2/caspase-3/GSDME signaling in neuroblastoma. Am J Cancer Res 13: 436–451

Fry AM, O’Regan L, Sabir SR, Bayliss R (2012) Cell cycle regulation by the NEK family of protein kinases. J Cell Sci 125: 4423–4433

Gau M, Karabajakian A, Reverdy T, Neidhardt EM, Fayette J (2019) Induction chemotherapy in head and neck cancers: Results and controversies. Oral Oncol 95: 164–169

Hernandez-Vargas H, Palacios J, Moreno-Bueno G (2007) Molecular profiling of docetaxel cytotoxicity in breast cancer cells: uncoupling of aberrant mitosis and apoptosis. Oncogene 26: 2902–2913

Hou J, Hsu JM, Hung MC (2021) Molecular mechanisms and functions of pyroptosis in inflammation and antitumor immunity. Mol Cell 81: 4579–4590

Hsieh CY, Lin CC, Chang WC (2023) Taxanes in the Treatment of Head and Neck Squamous Cell Carcinoma. Biomedicines 11

Jin L, Chun J, Pan C, Li D, Lin R, Alesi GN, Wang X, Kang HB, Song L, Wang D et al (2018) MAST1 Drives Cisplatin Resistance in Human Cancers by Rewiring cRaf-Independent MEK Activation. Cancer Cell 34: 315–330 e317

Kayagaki N, Warming S, Lamkanfi M, Vande Walle L, Louie S, Dong J, Newton K, Qu Y, Liu J, Heldens S et al (2011) Non-canonical inflammasome activation targets caspase-11. Nature 479: 117–121

Lu G, Du R, Dong J, Sun Y, Zhou F, Feng F, Feng B, Han Y, Shang Y (2023) Cancer associated fibroblast derived SLIT2 drives gastric cancer cell metastasis by activating NEK9. Cell Death Dis 14: 421

Lu G, Tian S, Sun Y, Dong J, Wang N, Zeng J, Nie Y, Wu K, Han Y, Feng B et al (2021) NEK9, a novel effector of IL-6/STAT3, regulates metastasis of gastric cancer by targeting ARHGEF2 phosphorylation. Theranostics 11: 2460–2474

Maji S, Samal SK, Pattanaik L, Panda S, Quinn BA, Das SK, Sarkar D, Pellecchia M, Fisher PB, Dash R (2015) Mcl-1 is an important therapeutic target for oral squamous cell carcinomas. Oncotarget 6: 16623–16637

Maji S, Shriwas O, Samal SK, Priyadarshini M, Rath R, Panda S, Das Majumdar SK, Muduly DK, Dash R (2019) STAT3- and GSK3beta-mediated Mcl-1 regulation modulates TPF resistance in oral squamous cell carcinoma. Carcinogenesis 40: 173–183

Mhaidat NM, Zhang XD, Jiang CC, Hersey P (2007) Docetaxel-induced apoptosis of human melanoma is mediated by activation of c-Jun NH2-terminal kinase and inhibited by the mitogen-activated protein kinase extracellular signal-regulated kinase 1/2 pathway. Clin Cancer Res 13: 1308–1314

Moffat J, Grueneberg DA, Yang X, Kim SY, Kloepfer AM, Hinkle G, Piqani B, Eisenhaure TM, Luo B, Grenier JK et al (2006) A lentiviral RNAi library for human and mouse genes applied to an arrayed viral high-content screen. Cell 124: 1283–1298

Mohanty S, Mohapatra P, Shriwas O, Ansari SA, Priyadarshini M, Priyadarsini S, Rath R, Sultania M, Das Majumdar SK, Swain RK et al (2022) CRISPR-based kinome-screening revealed MINK1 as a druggable player to rewire 5FU-resistance in OSCC through AKT/MDM2/p53 axis. Oncogene 41: 4929–4940

Mukhin YV, Garnovsky EA, Ullian ME, Garnovskaya MN (2003) Bradykinin B2 receptor activates extracellular signal-regulated protein kinase in mIMCD-3 cells via epidermal growth factor receptor transactivation. J Pharmacol Exp Ther 304: 968–977

Naviaux RK, Costanzi E, Haas M, Verma IM (1996) The pCL vector system: rapid production of helper-free, high-titer, recombinant retroviruses. J Virol 70: 5701–5705

Panchal NK, Evan Prince S (2023) The NEK family of serine/threonine kinases as a biomarker for cancer. Clin Exp Med 23: 17–30

Park SR, Speranza G, Piekarz R, Wright JJ, Kinders RJ, Wang L, Pfister T, Trepel JB, Lee MJ, Alarcon S et al (2013) A multi-histology trial of fostamatinib in patients with advanced colorectal, non-small cell lung, head and neck, thyroid, and renal cell carcinomas, and pheochromocytomas. Cancer Chemother Pharmacol 71: 981–990

Phadke M, Remsing Rix LL, Smalley I, Bryant AT, Luo Y, Lawrence HR, Schaible BJ, Chen YA, Rix U, Smalley KSM (2018) Dabrafenib inhibits the growth of BRAF-WT cancers through CDK16 and NEK9 inhibition. Mol Oncol 12: 74–88

Polak A, Bialopiotrowicz E, Krzymieniewska B, Wozniak J, Stojak M, Cybulska M, Kaniuga E, Mikula M, Jablonska E, Gorniak P et al (2020) SYK inhibition targets acute myeloid leukemia stem cells by blocking their oxidative metabolism. Cell Death Dis 11: 956

Rohila D, Park IH, Pham TV, Weitz J, Hurtado de Mendoza T, Madheswaran S, Ishfaq M, Beaman C, Tapia E, Sun S et al (2023) Syk Inhibition Reprograms Tumor-Associated Macrophages and Overcomes Gemcitabine-Induced Immunosuppression in Pancreatic Ductal Adenocarcinoma. Cancer Res 83: 2675–2689

Sanjana NE, Shalem O, Zhang F (2014) Improved vectors and genome-wide libraries for CRISPR screening. Nat Methods 11: 783–784

Schiff PB, Fant J, Horwitz SB (1979) Promotion of microtubule assembly in vitro by taxol. Nature 277: 665–667

Shi J, Zhao Y, Wang K, Shi X, Wang Y, Huang H, Zhuang Y, Cai T, Wang F, Shao F (2015) Cleavage of GSDMD by inflammatory caspases determines pyroptotic cell death. Nature 526: 660–665

Shriwas O, Arya R, Mohanty S, Mohapatra P, Kumar S, Rath R, Kaushik SR, Pahwa F, Murmu KC, Majumdar SKD et al (2021) RRBP1 rewires cisplatin resistance in oral squamous cell carcinoma by regulating Hippo pathway. Br J Cancer 124: 2004–2016

Smith SC, Petrova AV, Madden MZ, Wang H, Pan Y, Warren MD, Hardy CW, Liang D, Liu EA, Robinson MH et al (2014) A gemcitabine sensitivity screen identifies a role for NEK9 in the replication stress response. Nucleic Acids Res 42: 11517–11527

Tan Q, Liu X, Fu X, Li Q, Dou J, Zhai G (2012) Current development in nanoformulations of docetaxel. Expert Opin Drug Deliv 9: 975–990

Tang D, Wu D, Hirao A, Lahti JM, Liu L, Mazza B, Kidd VJ, Mak TW, Ingram AJ (2002) ERK activation mediates cell cycle arrest and apoptosis after DNA damage independently of p53. J Biol Chem 277: 12710–12717

Tzelepis K, Koike-Yusa H, De Braekeleer E, Li Y, Metzakopian E, Dovey OM, Mupo A, Grinkevich V, Li M, Mazan M et al (2016) A CRISPR Dropout Screen Identifies Genetic Vulnerabilities and Therapeutic Targets in Acute Myeloid Leukemia. Cell Rep 17: 1193–1205

Vermorken JB, Remenar E, van Herpen C, Gorlia T, Mesia R, Degardin M, Stewart JS, Jelic S, Betka J, Preiss JH et al (2007) Cisplatin, fluorouracil, and docetaxel in unresectable head and neck cancer. N Engl J Med 357: 1695–1704

Wang Y, Gao W, Shi X, Ding J, Liu W, He H, Wang K, Shao F (2017) Chemotherapy drugs induce pyroptosis through caspase-3 cleavage of a gasdermin. Nature 547: 99–103

Wani MC, Taylor HL, Wall ME, Coggon P, McPhail AT (1971) Plant antitumor agents. VI. The isolation and structure of taxol, a novel antileukemic and antitumor agent from Taxus brevifolia. J Am Chem Soc 93: 2325–2327

Wishart DS, Knox C, Guo AC, Shrivastava S, Hassanali M, Stothard P, Chang Z, Woolsey J (2006) DrugBank: a comprehensive resource for in silico drug discovery and exploration. Nucleic Acids Res 34: D668–672

Wu M, Wang Y, Yang D, Gong Y, Rao F, Liu R, Danna Y, Li J, Fan J, Chen J et al (2019) A PLK1 kinase inhibitor enhances the chemosensitivity of cisplatin by inducing pyroptosis in oesophageal squamous cell carcinoma. EBioMedicine 41: 244–255

Yamamoto Y, Chino H, Tsukamoto S, Ode KL, Ueda HR, Mizushima N (2021) NEK9 regulates primary cilia formation by acting as a selective autophagy adaptor for MYH9/myosin IIA. Nat Commun 12: 3292

Yan Y, Black CP, Cao PT, Haferbier JL, Kolb RH, Spieker RS, Ristow AM, Cowan KH (2008) Gamma-irradiation-induced DNA damage checkpoint activation involves feedback regulation between extracellular signal-regulated kinase 1/2 and BRCA1. Cancer Res 68: 5113–5121

Yvon AM, Wadsworth P, Jordan MA (1999) Taxol suppresses dynamics of individual microtubules in living human tumor cells. Mol Biol Cell 10: 947–959

