## Supplemental figure for "NEK9 ablation rewires docetaxel resistance through induction of ERK-mediated cancer cell pyroptosis"

### Supplementary figure 01

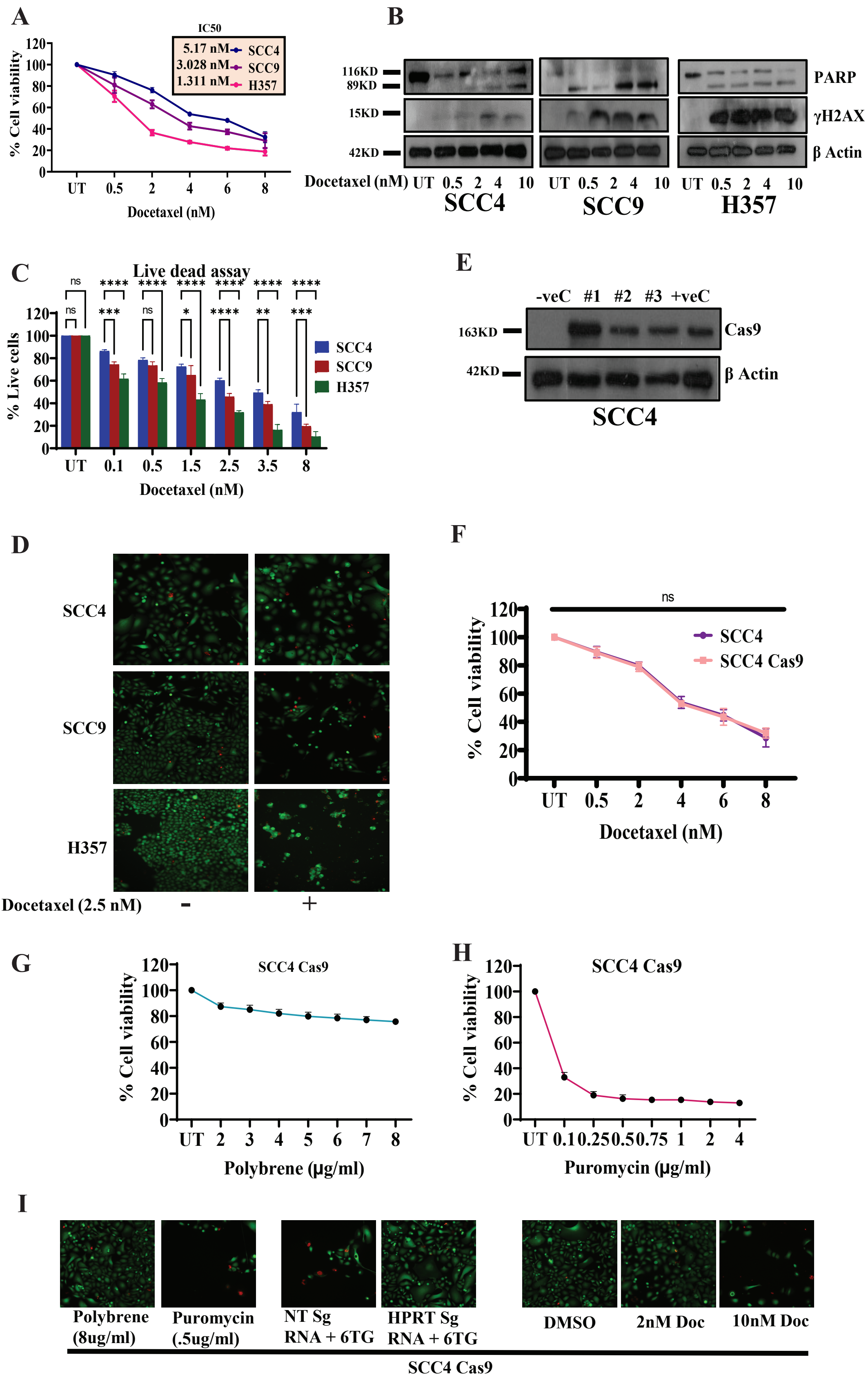

### Supplementary figure 02

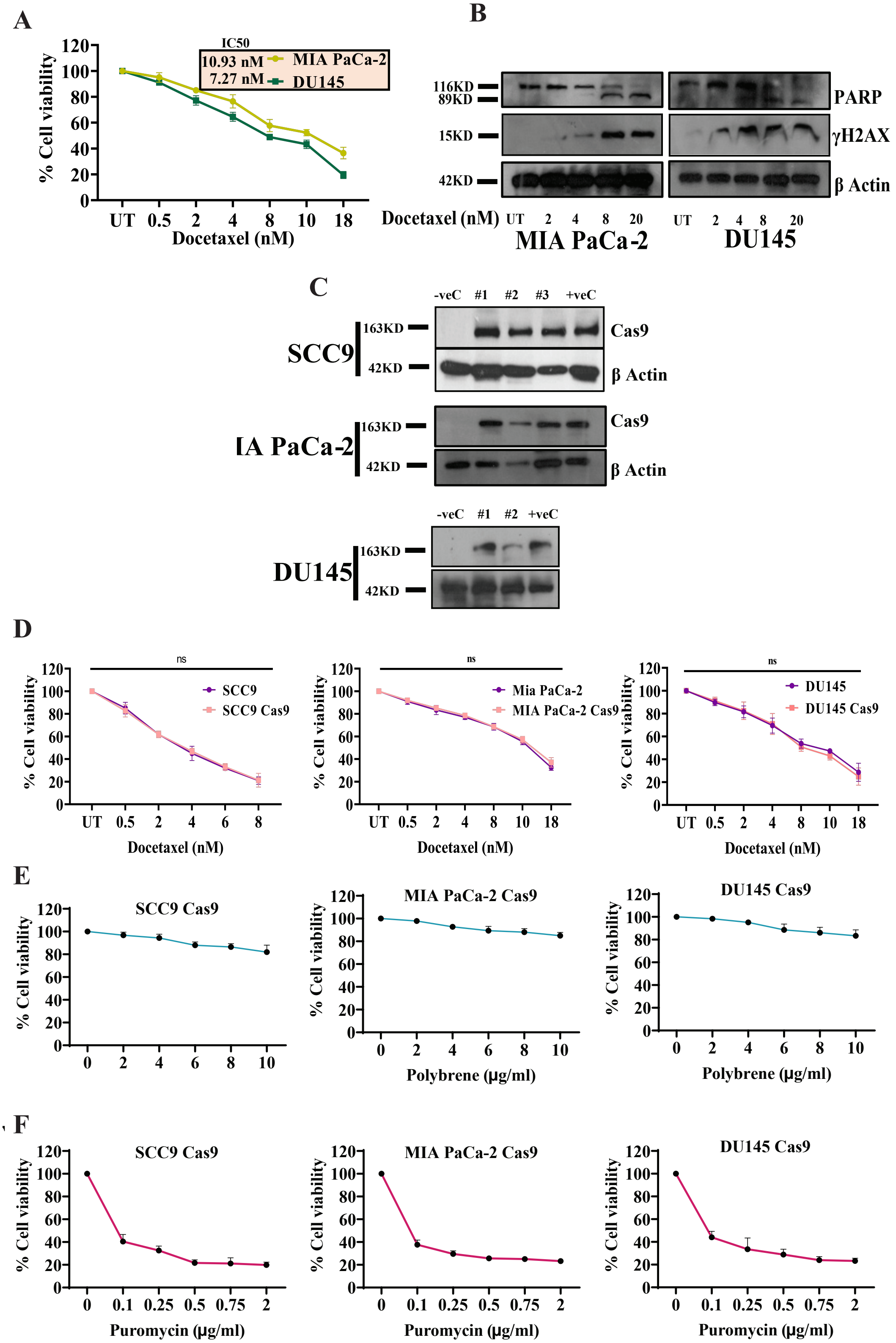

### Supplementary figure 03

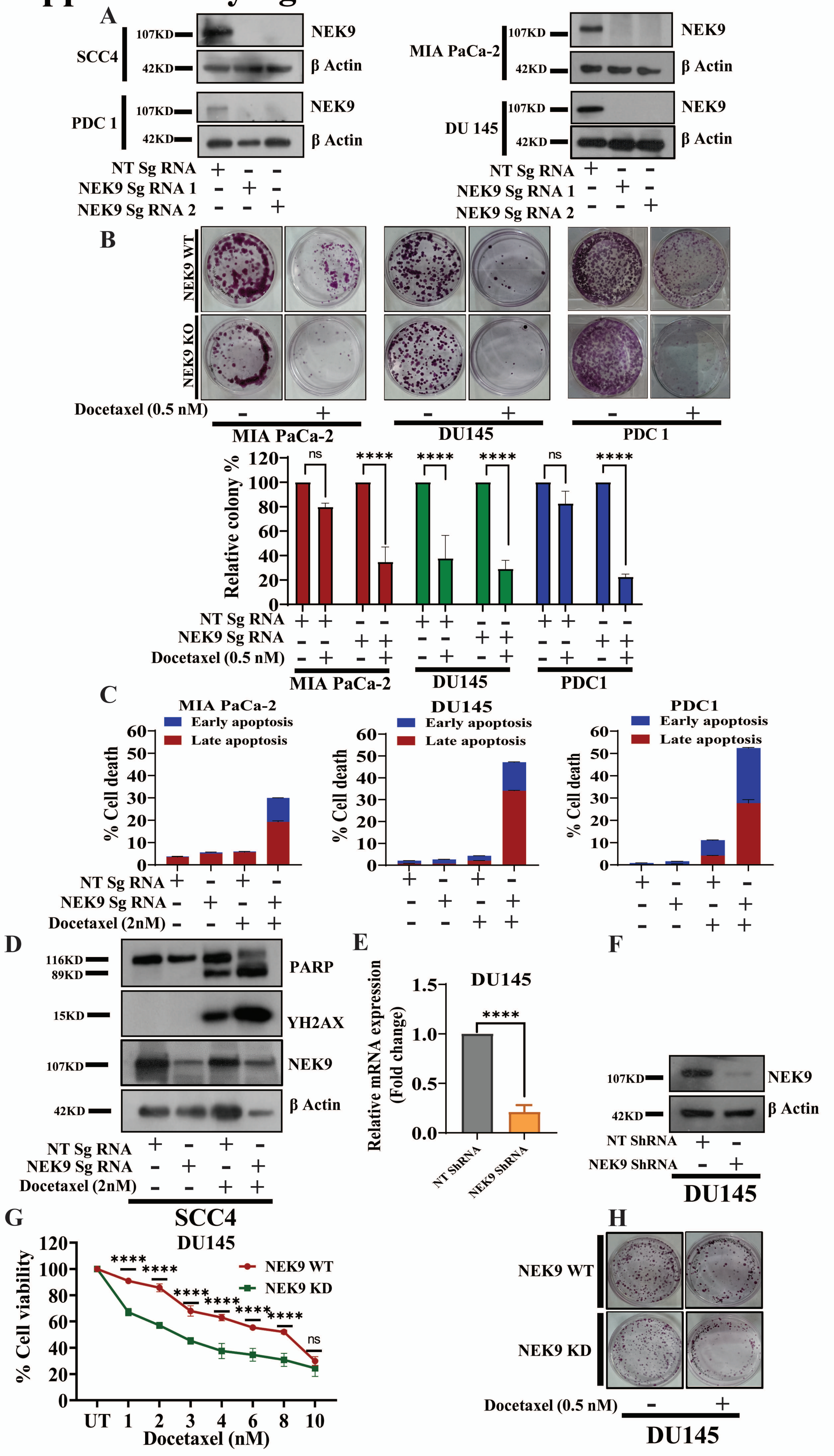

### Supplementary figure 04

**A**

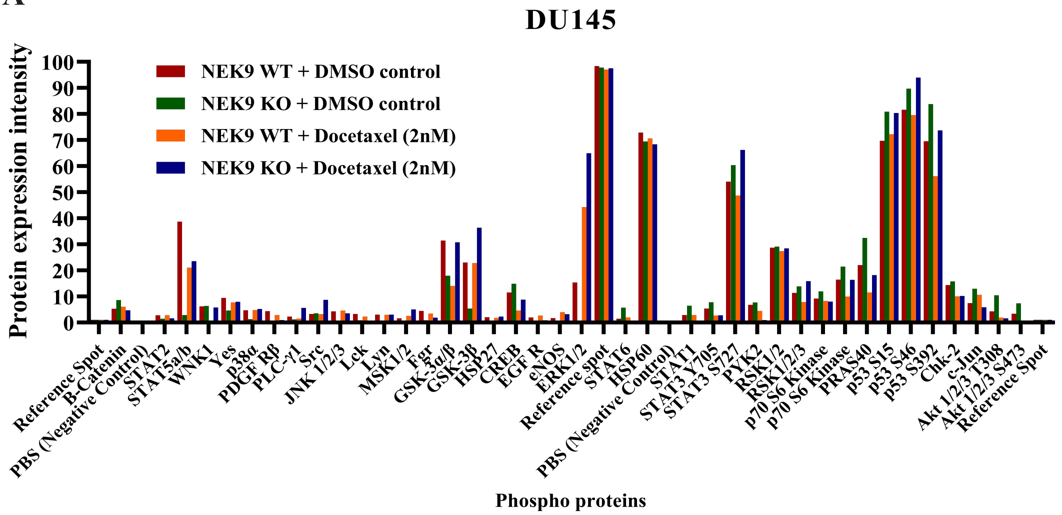

### Supplementary figure 05

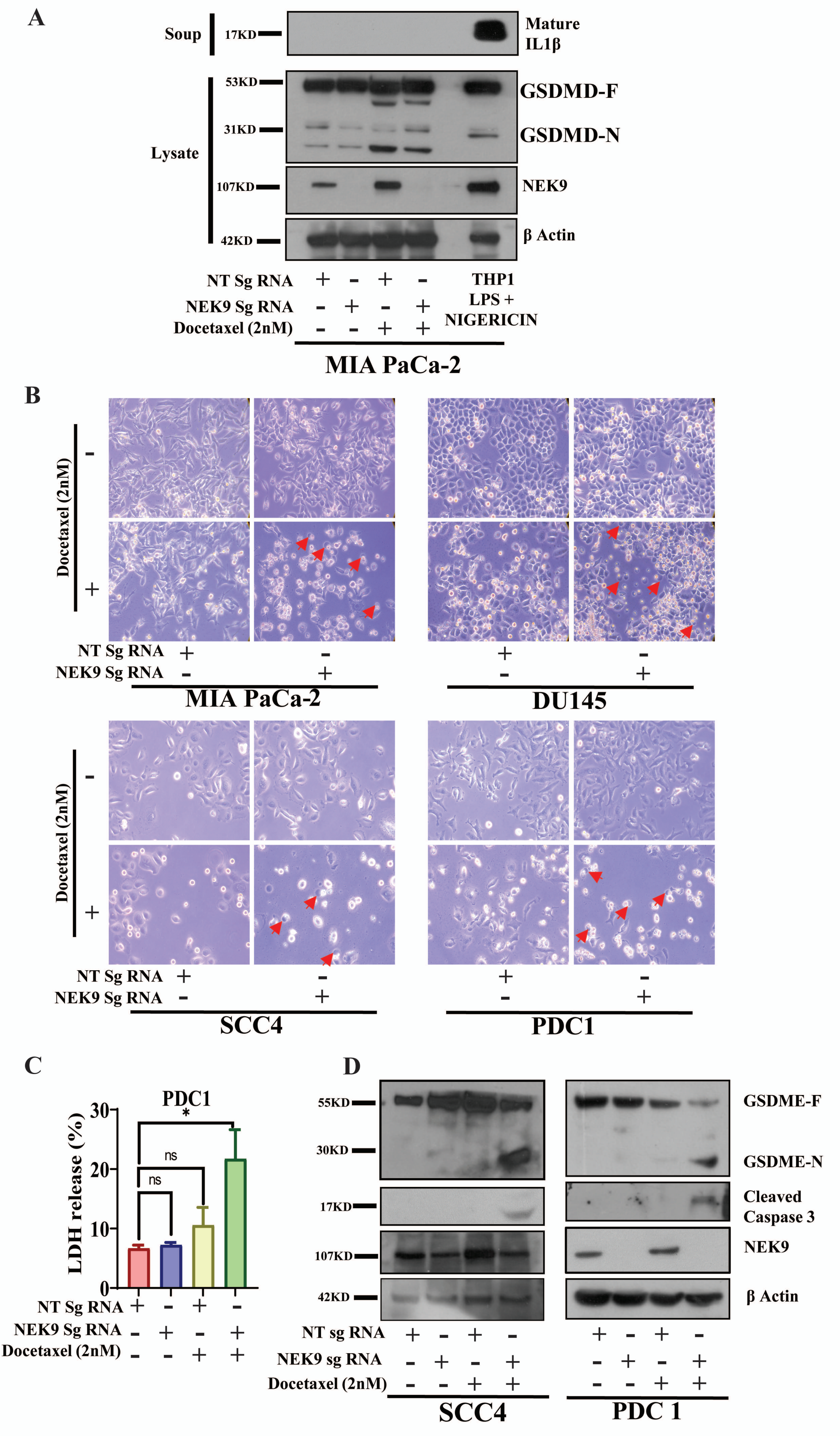

**Fig. S1 Comparative viability assays determine SCC4 cell line as the suitable OSCC candidate for primary kinome screening. A.** SCC4, SCC9 and H357 cell lines were treated with indicated concentration of docetaxel for 48hrs and cell viability was determined by MTT assay (n=3). **B.** Indicated OSCC cells were treated with docetaxel in dose dependent manner for 48 hrs, after which lysates were collected and immunoblotting was performed with indicated antibodies. **C.** The indicated cell types were treated with docetaxel for 48hrs and then stained with live dead viability/toxicity kit. Live and dead cells count and photographs in individual wells were taken with the help of Cell Insight CX7 High Content Screening (HCS) Platform (20 fields per well using 10X objective lens) on the basis of intensity (n=3 and 2-way ANOVA, *P<0.05, **P<0.01, ***P<0.001, ****P<0.0001). **D.** Representative images of docetaxel (2.5nM) treated and untreated cells from Cell Insight CX7 High Content Screening (HCS) Platform. **E.** SCC4 cells were stably transfected with LentiCas9-Blast and lysates from 3 clones along with negative and positive controls were subjected to immunoblotting with indicated antibodies. **F.** SCC4 and SCC4 Cas9 overexpressing cells were treated with indicated concentrations of drug for 48h and cell viability was accessed using MTT assay (n=3 and 2-way ANOVA, *P<0.05). **G.H.** SCC4 Cas9 overexpressing cells were treated with indicated concentrations of polybrene and puromycin respectively for 48hrs and MTT assay was performed to check the cell viability (n=3). **I.** Representative images of indicated control groups from the primary screening in SCC4 Cas9 overexpressing cells. The images were taken by cell Insight CX7 High Content Screening (HCS) system. Fig S1 results are the representatives of 2-3 independent experiments performed with triplicates.

**Fig. S2 Overexpression of Cas9 in cells and their characterization for secondary kinome screening. A.** Cell viability of MIA PaCa-2 and DU145 cells were treated with docetaxel in dose dependent manner and cell viability was measured by MTT assay (n=3). **B.** Lysates from MIA PaCa-2 and DU145 cells treated with indicated concentration of docetaxel for 48h were subjected to immunoblotting (n=3) with indicated antibodies. **C.** Cas9 was stably overexpressed in SCC9, MIA PaCa-2 and DU145 cells and overexpression was confirmed in the indicated no. of clones using immunoblotting technique (n=3) against indicated antibodies. **D.** Indicated Cas9 overexpressing cell lines and their parental counterparts were treated with indicated concentrations of docetaxel for 48hrs and cell viability was assessed with MTT assay (n=3 and 2-way ANOVA, *P<0.05). **E.** Indicated Cas9 overexpressing cells were treated with indicated concentrations of polybrene for 48h and cell viability was determined by MTT assay (n=3). **F.** Indicated Cas9 overexpressing cells were treated with indicated concentrations of puromycin for 48h and cell viability was determined by MTT assay (n=3).

**Fig. S3 Invitro validation of kinome screening results for NEK9. A.** Indicated Cas9 overexpressing cells were transduced with NT sgRNA and 2 different NEK9 sgRNA lentiviruses. NEK9 KO was confirmed by immunoblotting (n=3) against indicated antibodies. **B.** Left panel shows representative images of colony forming assay for NEK9 WT and KO cells treated with 0.5nM docetaxel for 12-14 days, and right panel represents the bar diagram for the relative colony % (n=2, 2-way ANOVA, ****P<0.0001). **C.** Indicated cells stably expressing NT SgRNA or NEK9 SgRNA were treated with docetaxel for 48h, after which cell death was measured by staining with annexin V/7AAD (n=2). **D.** SCC4 NEK9 WT and KO cells were treated with docetaxel for 48 hours and immunoblotting was performed with the indicated antibodies. **E. F.** DU145 cells were transduced with indicated NEK9 shRNA lentiviruses and KD was confirmed by qRT-PCR (**A**) (n=3, unpaired t-test, ****P<0.0001), and immunoblotting (**F**) with indicated antibodies. **G.** DU145 NEK9 WT and KD cells were treated with indicated concentrations of docetaxel for 48h and cell viability was accessed by MTT assay (n=3 and 2-way ANOVA, ****P<0.0001). **H.** Representative images of colony forming assay for DU145 NEK9 WT and KD cells treated with 0.5nM docetaxel for 12-14 days.

**Fig. S4 NEK9 depletion enhances phosphorylation of ERK1/2 T202/Y204 upon docetaxel treatment. A.** Lysates from DU145 NEK9 WT and KO cells prior treated for 48 hrs with docetaxel were subjected to Human Phospho Kinase array, to detect relative levels of phosphorylation of 37 kinase phosphorylation sites and 2 related total proteins. The bars represent the relative signals for the phospho-proteins from the indicated cells.

**Fig. S5 Pyroptosis in NEK9 ablated cells do not follow canonical ICP pathways but undergoes cell death by cancer cell pyroptosis. A.** Proteins from cultured media supernatants were isolated by methanol/chloroform precipitation assay from the indicated cells. Immunoblotting was performed with mature IL1β antibody. Whole cell lysates were isolated from the indicated cells and immunoblotting was carried out for indicated antibodies. THP1 monocytic cells stimulated with LPS in combination with Nigericin were used as positive control. **B.** Morphological changes in indicated NEK9 KO cells upon docetaxel treatment. Arrow indicates cell swealing. **C.** LDH release % was detected in the indicated cells after 48hrs of drug treatment (n=2, Ordinary one-way ANOVA, *P=0.0213). **D.** NEK9 WT and KO cells were treated with docetaxel for 48 hours and immunoblotting was performed with the indicated antibodies.
